# Comparative genomics reveals electron transfer and syntrophic mechanisms differentiating methanotrophic and methanogenic archaea

**DOI:** 10.1101/2021.09.25.461819

**Authors:** Grayson L Chadwick, Connor T Skennerton, Rafael Laso-Pérez, Andy O Leu, Daan R Speth, Hang Yu, Connor Morgan-Lang, Roland Hatzenpichler, Danielle Goudeau, Rex Malmstrom, William J Brazelton, Tanja Woyke, Steven J Hallam, Gene W Tyson, Gunter Wegener, Antje Boetius, Victoria J Orphan

**Author notes:** Address correspondence to Grayson L Chadwick,; Victoria J Orphan. These authors contributed equally to this work (Grayson L Chadwick, Connor T Skennerton). Department of Chemistry and Biochemistry, Thermal Biology Institute, and Center for Biofilm Engineering, Montana State University, Bozeman, MT, USA 59717. Department of Molecular and Cell Biology, University of California Berkeley, Berkeley, CA, USA 94720-3220.

## Abstract

The anaerobic oxidation of methane coupled to sulfate reduction is a microbially mediated process requiring a syntrophic partnership between anaerobic methanotrophic (ANME) archaea and sulfate reducing bacteria (SRB). Based on genome taxonomy, ANME lineages are polyphyletic within the phylum *Halobacterota*, none of which have been isolated in pure culture. Here we reconstruct 28 ANME genomes from environmental metagenomes and flow sorted syntrophic consortia. Together with a reanalysis of previously published datasets, these genomes enable a comparative analysis of all marine ANME clades. We review the genomic features which separate ANME from their methanogenic relatives and identify what differentiates ANME clades. Large multiheme cytochromes and bioenergetic complexes predicted to be involved in novel electron bifurcation reactions are well-distributed and conserved in the ANME archaea, while significant variations in the anabolic C1 pathways exists between clades. Our analysis raises the possibility that methylotrophic methanogenesis may have evolved from a methanotrophic ancestor.

## Introduction

Anaerobic oxidation of methane (AOM) coupled to sulfate reduction is a key microbiological process in ocean sediments that controls the amount of methane released into overlying waters and the atmosphere. However, despite the global relevance and distribution of this process, there are currently no strain isolates that will carry out AOM with sulfate. Thus, our understanding of the physiological and biochemical basis for AOM has advanced much more slowly than it has for many other microbially-mediated biogeochemical processes. Twenty years ago, strong evidence emerged that archaea may be involved in AOM based on stable isotope measurements of archaeal lipids and small subunit ribosomal RNA (SSU or 16S rRNA) gene clone libraries from marine methane seeps (1). Shortly thereafter fluorescence *in situ* hybridization from methane seep environments provided microscopic evidence for the existence of a prevalent interdomain consortia consisting of an archaeon related to known methanogens and a bacterium related to sulfate reducing bacteria (2, 3). These discoveries led to the current paradigm that sulfate-dependent AOM is carried out by anaerobic methanotrophic (ANME) archaea in a syntrophic partnership with sulfate reducing bacteria (SRB).

Subsequent work has expanded our understanding diversity and activities of the ANME archaea and lead to various hypotheses pertaining to the biochemical mechanisms underlying the syntrophic interactions between ANME and SRB. Diversity surveys have suggested that ANME are polyphyletic with three distinct clades (ANME-1, 2, and 3) within the *Halobacterota*. Investigations of 16S rRNA gene phylogenies support ANME-1 as a family-level clade, while ANME-2 is comprised of two distinct families within the *Methanosarcinales*, and members of ANME-3 are a novel genus closely related to *Methanococcoides* within the family *Methanosarcinaceae* (4, 5). Initial ‘omic analysis of fosmid libraries from ANME organisms supported “reverse methanogenesis” as the biochemical model for the methane oxidation pathway in AOM (6–8). Subsequent analysis of more complete ANME genomes from enrichment cultures (9–11), or metagenome assembled genomes (MAGs) from AOM habitats have added to our understanding of some of the major groups of ANME, and further refined the “reverse methanogenesis hypothesis” (12, 13).

The limited number of ANME genomes currently available relative to their 16S rRNA gene diversity leads to questions about whether the observations made in previous studies represent conserved features of the ANME archaea, or are skewed by the relatively small sample size and the incomplete or biased nature of metagenomic binning methods. In order to develop a better model for the evolution and metabolic capabilities of the ANME archaea we performed a large comparative analysis of the most complete set of ANME genomes to date, encompassing 39 reconstructed MAGs, binned fosmid libraries, amplified single aggregate genomes (ASAGs), and combine-assembled single amplified genomes (Co-SAGs), more than doubling the previously available genomic information. This analysis includes representatives from all recognized marine ANME groups including ANME-1, 2a, 2b, 2c, and 3. Using this expanded dataset we construct a more robust phylogenetic framework for the ANME archaea and analyze the differences in metabolic potential that exist within and among ANME clades, as well as between ANME and their methanogenic relatives with emphasis on energy conservation and potential adaptations to syntrophic associations with SRB. Based on these genomic observations we present a model for ANME metabolism for the different clades and highlight important aspects of their physiology that remain uncertain.

## Results

### Genome-resolved diversity of the methanotrophic archaea

We reconstructed 28 new ANME genomes from metagenomic datasets and fluorescence-activated cell sorting of single aggregates (14), as well as from reanalyzed publicly available metagenomic data from the sequence read archive (SRA) and the MG-RAST analysis server (see **Materials and Methods**). These genomes were combined with 11 previously published marine ANME genomes recovered from diverse environments (7, 9, 11, 15–17) to generate a set of 39 ANME genomes for comparative genomics, representing all of the currently described clades (ANME-1, 2, 3) and subclades (e.g. ANME-2a, b, c and multiple clades in ANME-1) frequently detected in marine sediments (**Table 1**)(4). In most analyses we also include the recently described alkane-oxidizing “*Candidatus* Syntrophoarchaeum” (18), as well as the nitrate-reducing freshwater ANME relatives known as ANME-2d or “*Candidatus* Methanoperedens” (19, 20) and their marine relatives known as “*Candidatus* Argoarchaeum” (15, 21). At least one genus from each ANME subclade was assigned a name and formal etymology can be found in **Materials and Methods**.

**Table 1.**
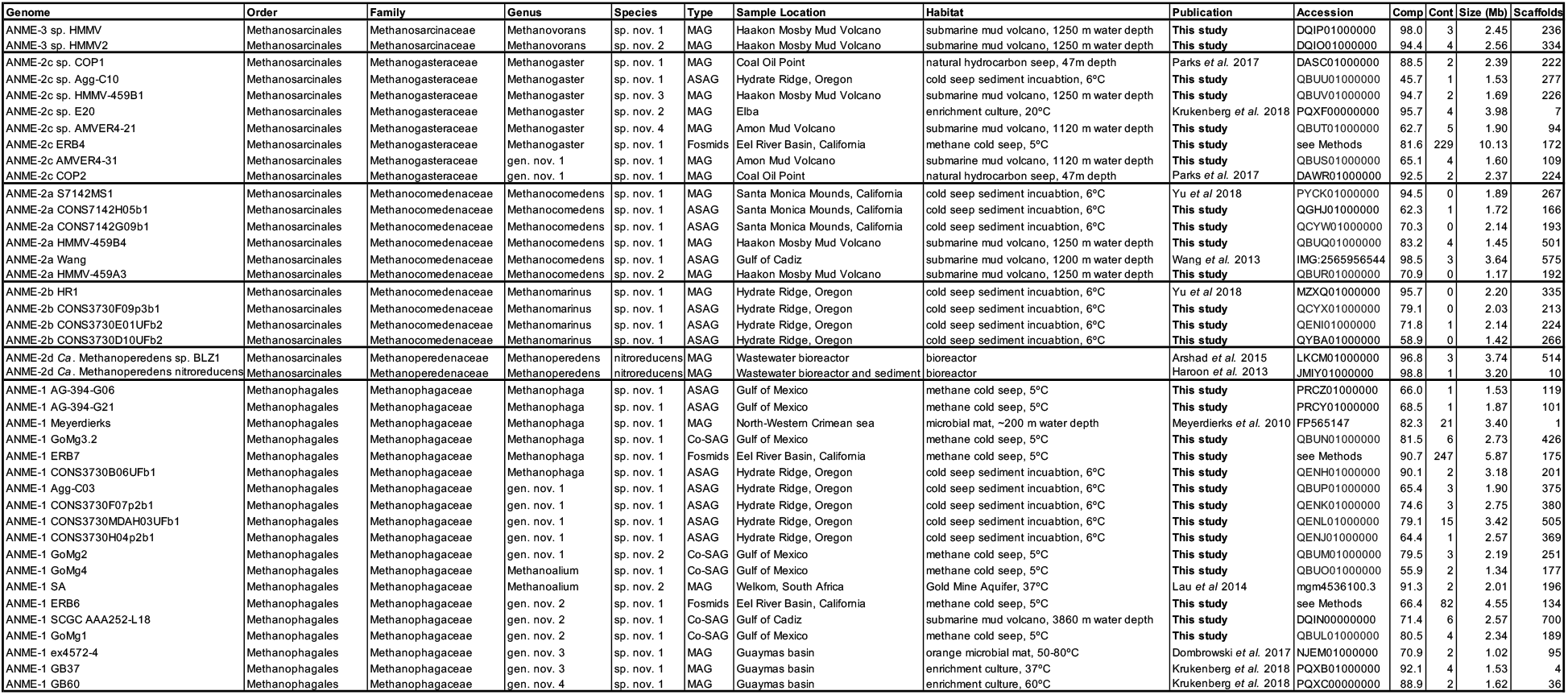
ANME genome statistics. Summary of genome characteristics, sources and accession numbers of the analyzed ANME genomes. Major ANME clades are separated by bold lines and proposed taxonomy adheres to the GTDB framework. At least one genus within each clade has been given a proposed name, and genomes that fall within other genera or species are simply given numerical placeholders (Genus and Species columns, respectively). Type indicates the method used for genome reconstruction, including: metagenome assembled genome (MAG), amplified single aggregate genome (ASAG), combine-assembled single amplified genomes (Co-SAGs), or fosmid library. Estimated levels of completeness (comp) and contamination (cont) are reported as percent as evaluated by CheckM. Note: ERB4, ERB6 and ERB7 are fosmid libraries with highly similar strains combined which has resulted in high levels of “contamination”. See **Materials and Methods** for complete details of assembly and binning for each genome.

The draft ANME genomes sequenced and assembled here were between 46% and 98% complete (mean: 78%, median: 80%) as determined by the presence of a set of marker proteins common to *Halobacterota*. Many of the less complete genomes originated from sequencing of individual flow sorted aggregates (**Table 1**). Most of the genomes did not contain duplicated marker genes, with the exception of the three genomes derived from sequenced fosmids, which encoded many duplicates (**Table 1**). The fosmids representing these genomes have consistent nucleotide signatures and, in many cases, contain overlapping regions that suggests that they are derived from multiple strains rather than from a single clonal population. While genome incompleteness can impact the accurate reconstruction of ANME metabolism, we found reproducible trends in gene presence, absence, and synteny across the different ANME lineages, with multiple genomes for each ANME clade, often originating from different studies and habitats. Taxonomic assignments in **Table 1** were made consistent with GTDB release 89 using analysis of relative evolutionary divergence (RED) (22).

Members of the ANME-1 were originally described in 16S rRNA gene surveys of methane seep sediments (1) and representatives of the ANME-1 were among the first to be genomically characterized (6). ANME-1 16S rRNA genes have since been identified in marine cold seep environments, diffusive margin sulfate-methane transition zones, deep sea hydrothermal vents, and select anoxic terrestrial ecosystems (23). With our updated genomic dataset there are now 19 representative genomes from ANME-1, recovered from eight locations, including deep-sea methane cold seeps, hydrocarbon-impacted hydrothermal vents and cold seeps, a mud volcano and a hot, deep gold mine aquifer. Collectively, these genomes are highly diverse at the sequence level with the majority being at most 60% similar to each other, based on pairwise sequence similarity and non-homologous gene content between genomes (**S1 Fig**). These genomes represent six genera as determined by analysis of their RED values. Recently proposed nomenclature based on a single ANME-1 genome from fosmid sequences placed ANME-1 within their own order *Methanophagales* (24), and this is consistent with GTDB release 89. The 19 ANME-1 genomes analyzed here represent a single family-level division within that order. We propose to conserve the family and genus-level designation implied by “*Methanophagales*” with the ANME-1 genome from Meyerdierks *et al*. 2010 belonging to the genus “*Candidatus* Methanophaga” within the family *Methanophagaceae* (**Table 1**).

Marine members of the ANME-2 are within the order *Methanosarcinales* and are comprised of subclades a, b, and c, first designated by16S rRNA gene sequences (5). The ANME-2a and 2b together form a family-level division with two genus-level clades corresponding to ANME-2a and 2b, recovered from six different locations including two methane cold seep sites, two submarine mud volcanoes, a hydrocarbon seep and shallow coastal sandy sediments (**Table 1**). We propose the name “*Candidatus* Methanocomedens” for the dominant genera corresponding to ANME-2a with name propagating to the family level as *Methanocomedenaceae*. Recent coupled fluorescence and electron microscopy analyses have revealed distinct ultrastructural features of the ANME-2b (25), and in conjunction with our phylogenomic information, we propose the genus name “*Candidatus* Methanomarinus” for ANME-2b. The six “*Ca.* Methanocomedens” genomes have average amino acid identity (AAI) values which range between 70% and 97% indicating that there are multiple distinct species, whereas the three “*Ca.* Methanomarinus” genomes have >98% AAI indicating that they represent strains of the same species (**S1 Fig**). The ANME-2c form a separate family (84% AAI among the 8 genomes analyzed in this study) representing two genus-level clades. For the dominant ANME-2c we propose the genus name “*Candidatus* Methanogaster” with name propagating to the family level as *Methanogasteraceae*.

“*Ca.* Methanoperedens” sp. BLZ1 and nitroreducens are representatives representative of the family *Methanoperedenaceae* formerly known as ANME-2d, within the order *Methanosarcinales*. These are the only known ANME that do not couple AOM in syntrophy with partner SRB, instead coupling the oxidation of methane with the reduction of nitrate, iron oxides or manganese oxides in freshwater environments (20, 26, 27). The marine sister group of “*Ca.* Methanoperedens”, GoM Arc I (also known as AAA), was recently described as an anaerobic ethane degrader, and contains two genera “*Candidatus* Argoarchaeum” (15, 21) and the thermophilic “*Candidatus* Ethanoperedens”(28). These clades are not specifically considered in the present work as they do not appear to be marine methanotrophs, and have been extensively discussed in a recent study (29).

The ANME-3 clade is the ANME group most recently diverged from known methanogens. They are closely related by their 16S rRNA genes (94-96% similarity) to *Methanococcoides*, *Methanosalsum*, and *Methanolobus*, with 65% average AAI, indicating these organisms represent a novel genus within the family *Methanosarcinaceae*, for which we propose the name “*Candidatus* Methanovorans” (**Table 1, S1 Fig**). The two genomes were both recovered from the Haakon Mosby Mud Volcano where this clade was originally described on the basis of 16S rRNA gene sequences (30). ANME-3 forms consortia with bacteria from the *Desulfobulbus* group (31).

Phylogenetic reconstructions of 16S rRNA gene sequences, concatenated marker genes, DNA-directed RNA polymerase subunit beta (RpoB) and the methane activating enzyme methyl-coenzyme M reductase subunit A (McrA) were performed to demonstrate the evolutionary relationship between ANME and other related archaea (**Fig 1**). All marker gene sets show remarkably similar phylogenies, with ANME-1 forming the deepest branching ANME clade, while ANME-2 and 3 grouped within the order *Methanosarcinales*. Importantly, in agreement with previous studies, ANME-3 reproducibly branches well within the family *Methanosarcinaceae*, in agreement with the AAI analysis described above. The McrA phylogeny deviates somewhat from this pattern, with ANME-1 falling further outside of the traditional methanogens, but grouping with some McrA from the H_2_-dependent methylotrophic methanogens as has been previously reported (12, 32, 33). The McrA in all ANME clades are more similar to methanogens than the recently described divergent McrA homologs in various uncultured archaea that are thought to utilize longer chain alkane substrates (15, 18, 21). With this general phylogenetic framework, we set out to characterize the major conserved features of the ANME energy metabolism.

**Fig 1.**
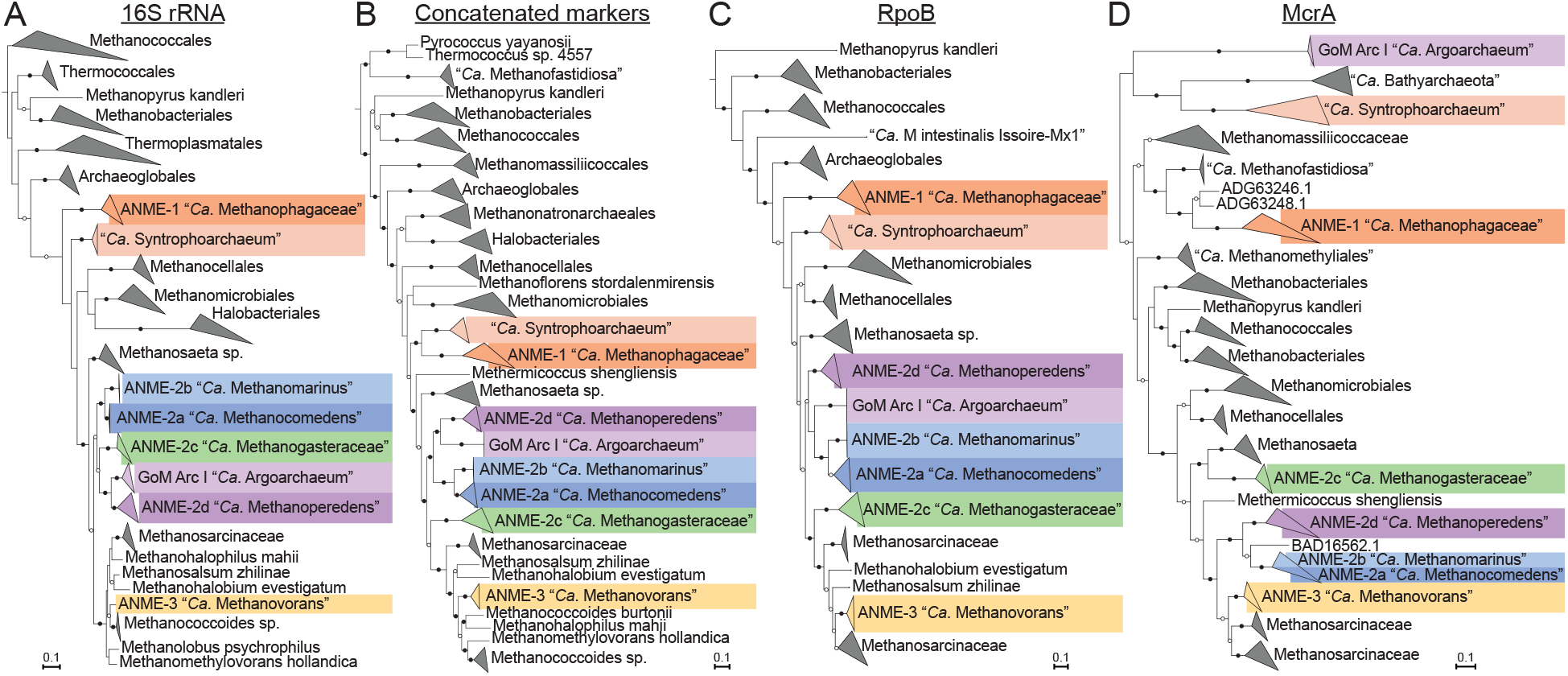
Phylogeny of ANME and related archaea. Phylogenetic trees constructed from ANME genomes, sequences of closely related archaea, and a selection of sequences derived from clone libraries demonstrate the relationship between ANME and methanogens of the *Halobacterota*. (**A**) Phylogenetic tree built with 16S rRNA gene sequences, root leads to sequence from *Sulfolobus solfataricus* p2. (**B**) Phylogenomic tree built with concatenated marker set 4 (see **S1 Table** for list), root also to *S. solfataricus* p2. (**C**) Phylogenetic tree built with protein sequence of RpoB, root leads to sequences from “*Ca*. Methanomethyliales” and “*Ca*. Bathyarchaeota”. (**D**) Phylogenetic tree of McrA protein sequences. Note the divergence of proposed alkane oxidizing McrA genes in “*Ca*. Syntrophoarchaeum”, “*Ca*. Bathyarchaeota” and “*Ca.* Argoarchaeum”. Branch support values of 100% are labelled with closed circles, >50% with open circles. Tree scales represent substitutions per site. Tree construction parameters are found in the **Materials and Methods** section. For tree files and alignments see **S1 Data**. Detailed tree figures are presented in **S2-S5 Figs**.

### ANME energy metabolism

Biochemically it is assumed that the ANME archaea oxidize methane to CO_2_ and pass electrons through an unconfirmed mechanism to their SRB partners to reduce sulfate. Metabolic reconstruction of a limited number of ANME draft genomes from environmental samples and enrichment cultures using multiple ‘omics approaches have shown that they contain and express the same genes used for methanogenesis (6, 7, 9, 10, 20, 34). Detailed enzymatic studies of the methanogenic pathway have shown that all 7 steps are reversible (35–37), including the methane producing step catalyzed by Mcr (38), which had previously been predicted to be irreversible. These findings reinforce the ‘reverse methanogenesis’ hypothesis (6, 39–41), in which ANME use the same methanogenic enzymes to oxidize, rather than produce, methane. This model offers the most likely pathway for carbon oxidation in ANME, however the mechanisms by which these archaea conserve energy from this process, and how methane-derived electrons are transferred to their syntrophic sulfate-reducing bacterial partners remain open questions.

It has been pointed out in recent reviews that a simple reversal of the methanogenesis pathway does not represent a viable basis for an energy metabolism in the ANME archaea, as the exact reversal of a process that results in net ATP generation must result in ATP loss (13, 42). Bioenergetic novelty beyond a wholesale reversal of methanogenesis therefore must exist. In the following discussion we break down the ANME energy metabolism into three phases and discuss the conserved features of these phases across our collection of ANME genomes (**Fig 2**). In the first phase, methane is oxidized to CO_2_ and all eight electrons are deposited on cytoplasmic electron carriers in an endergonic process that requires an investment of energy. In the second phase, cytoplasmic electron carriers are re-oxidized in an exergonic process that reduces a set of intermediate electron carriers, recovering the energy invested in the first phase and conserving additional energy as an ion motive force. In the final phase, electrons must be discarded, likely not involving energy gain or loss. We use this division of energy metabolism as an organizing framework throughout this work.

**Fig 2.**
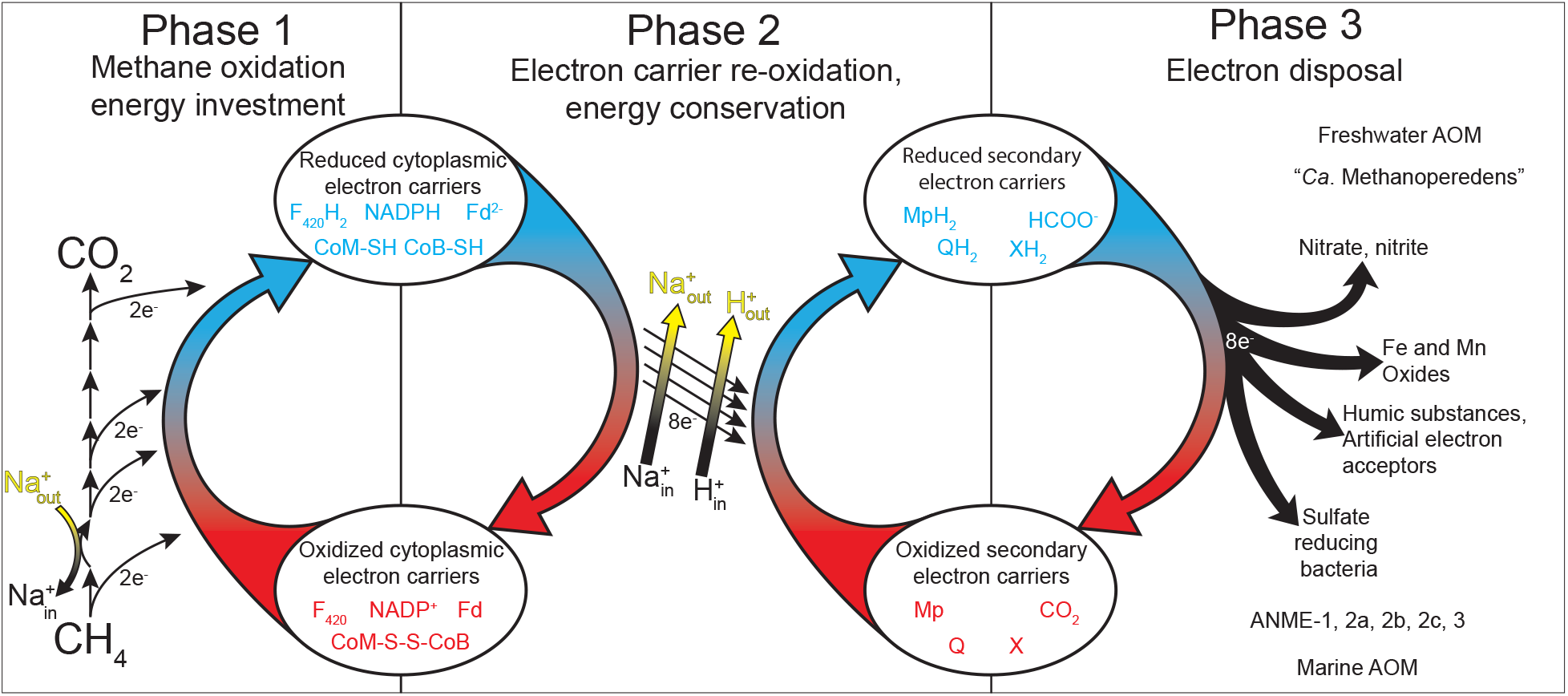
Summary of ANME energy metabolism. Schematic representation of the three phases of ANME energy metabolism in our current model. In Phase 1 methane is oxidized to CO_2_ through the reversal of the canonical seven step methanogenesis pathway. Energy is invested in this phase in the form of sodium ion translocation from the outer face of the cytoplasmic membrane to the inner face (yellow arrow). As C1 moieties are sequentially oxidized, eight electrons are transferred to soluble electron carriers such as F_420_H_2_, NADPH, Fd^2-^, and CoM-SH/CoB-SH. In Phase 2 eight electrons on these primary electron carriers are transferred to secondary electron carriers in a process that conserves energy needed for cell growth in the form of sodium and proton motive forces (yellow arrows). These secondary electron carriers may be quinols (QH_2_) methanophenazine (MpH_2_) or possibly soluble electron carriers such as formate (HCOO-) or an unknown electron shuttle (XH_2_). In Phase 3 the secondary electron carriers are relieved of their electrons in various ways depending on the environmentally available electron acceptors, which can include sulfate reducing bacteria in the case of marine ANME-SRB consortia, iron, manganese or oxidized nitrogen species in the case of “*Ca.* Methanoperedens”. Humic substances and artificial electron acceptors (AQDS) have also served as electron acceptors in laboratory experiments for a variety of different ANME from fresh and marine environments.

### Energy metabolism phase 1: The conserved C1 machinery of the methanogenesis pathway in ANME archaea

#### Function of the methanogenesis pathway in ANME archaea

Our analysis of this expanded set of ANME genomes is consistent with early genomic work that identified reverse methanogenesis as the most likely pathway of carbon oxidation in the ANME archaea (6, 7, 11). Genes for all seven steps of the methanogenesis pathway were found in all ANME clades (**Fig 3**). The only consistent exception is F_420_-dependent methylenetetrahydromethanopterin reductase (Mer), which is absent from all 19 ANME-1 genomes as well as the “*Ca.* Syntrophoarchaeum”, as observed previously (6, 7, 10, 18). This modification of the canonical methanogenesis pathway is a common feature of the entire class Syntrophoarchaeia. Some ANME contain paralogous copies of enzymes carrying out certain steps of the pathway and these are indicated in **Fig 3**. Notably, none of the ANME genomes contain the specific methyltransferases for methanol (43), methylamines (44) or methylated sulfur compounds (45) used for methylotrophic methanogenesis in the *Methanosarcinaceae*. This strongly argues against ANME archaea using methylated compounds as intermediates or end products of methane oxidation.

**Fig 3.**
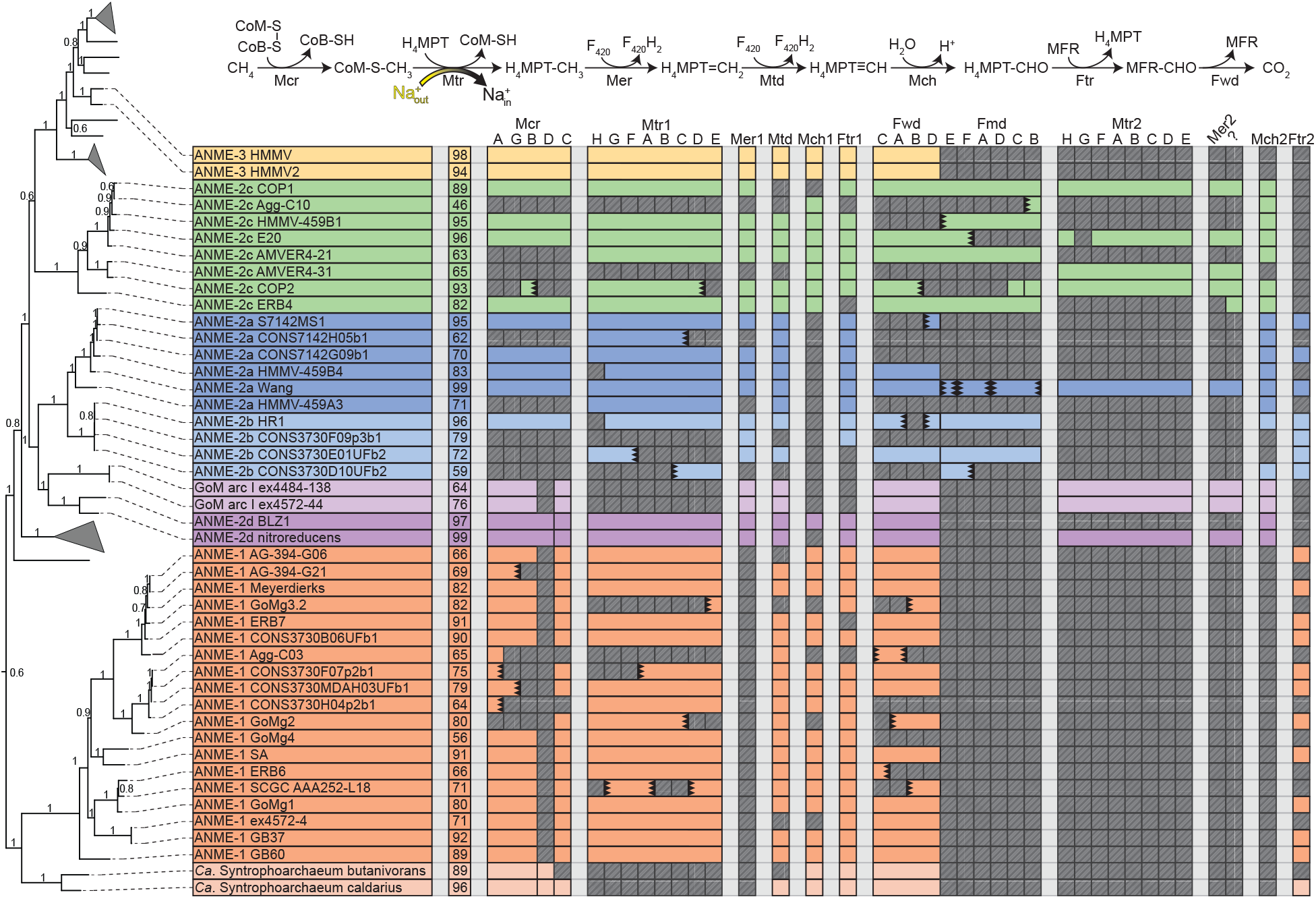
Presence of methanogenesis pathway genes in ANME archaea. The proteins responsible for the seven steps of methanogenesis from CO_2_. Mcr: Methyl-coenzyme M reductase; Mtr: N^5^-methyl-H_4_MPT:coenzyme M methyltransferase; Mer: methylene-H_4_MPT reductase; Mtd: F_420_-dependent methylene-H_4_MPT dehydrogenase; Mch: N^5^,N^10^-methenyl-H_4_MPT cyclohydrolase; Ftr: formylmethanofuran-H_4_MPT formyltransferase; Fmd/Fwd: formyl-methanofuran dehydrogenase. Colored boxes represent presence of homologs of these proteins in ANME genomes. Missing genes are represented by gray boxes with diagonal line fill. Numbers in the second column represent estimated genome completeness. When genes are together in a gene cluster their boxes are displayed fused together. If a gene cluster appears truncated by the end of a contig it is depicted by a serrated edge on the box representing the last gene on the contig. Numbers following protein names indicate whether the enzyme is closely related to those found in *Methanosarcinaceae* (1) or are distantly related homologs (2). Question mark represents hypothetical protein of unknown function found clustered with Mer2. Tree orienting genome order is the same as found Fig 1B. For details on paralog phylogenetic relations see Fig 4. Gene accession numbers can be found in **S2 Data**.

To understand whether the transition to a methanotrophic lifestyle is accompanied by a significant diversification of the enzymes catalyzing the reactions of the reverse methanogenesis pathway we built phylogenetic trees of the enzymes involved in each step. With regard to the first step, i.e. the presumed involvement of the McrA in methane activation, the McrA phylogeny largely tracks with phylogenetic marker trees, except for ANME-1 (**Fig 1**). The second step of the pathway is carried out by the N^5^-methyl-H_4_MPT:coenzyme M methyltransferase (Mtr) complex (**Fig 3**). All ANME clades contain Mtr homologs that are phylogenetically consistent with their genome phylogeny (**Fig 4A**). However, a highly divergent second copy of the entire Mtr complex exists that was first observed in the genome of an ANME-2a from an enrichment culture (11). We find this second “Mtr2” to be sporadically distributed through the ANME-2. Absent from the other ANME-2a genomes, Mtr2 is present in ANME-2c, one ANME-2d and both “*Candidatus* Argoarchaeum”. These Mtr2 complexes form a monophyletic group that is phylogenetically distinct from all previously described methanogens yet still contains the same gene synteny in the eight-subunit cluster. In addition, all the Mtr2 gene clusters are upstream of a highly divergent Mer2, which catalyzes the third step in the methane oxidation pathway. All ANME-2 and 3 clades contain a less divergent copy of Mer that tracks with their genome phylogeny (Mer1) (**Fig 4B**).

**Fig. 4.**
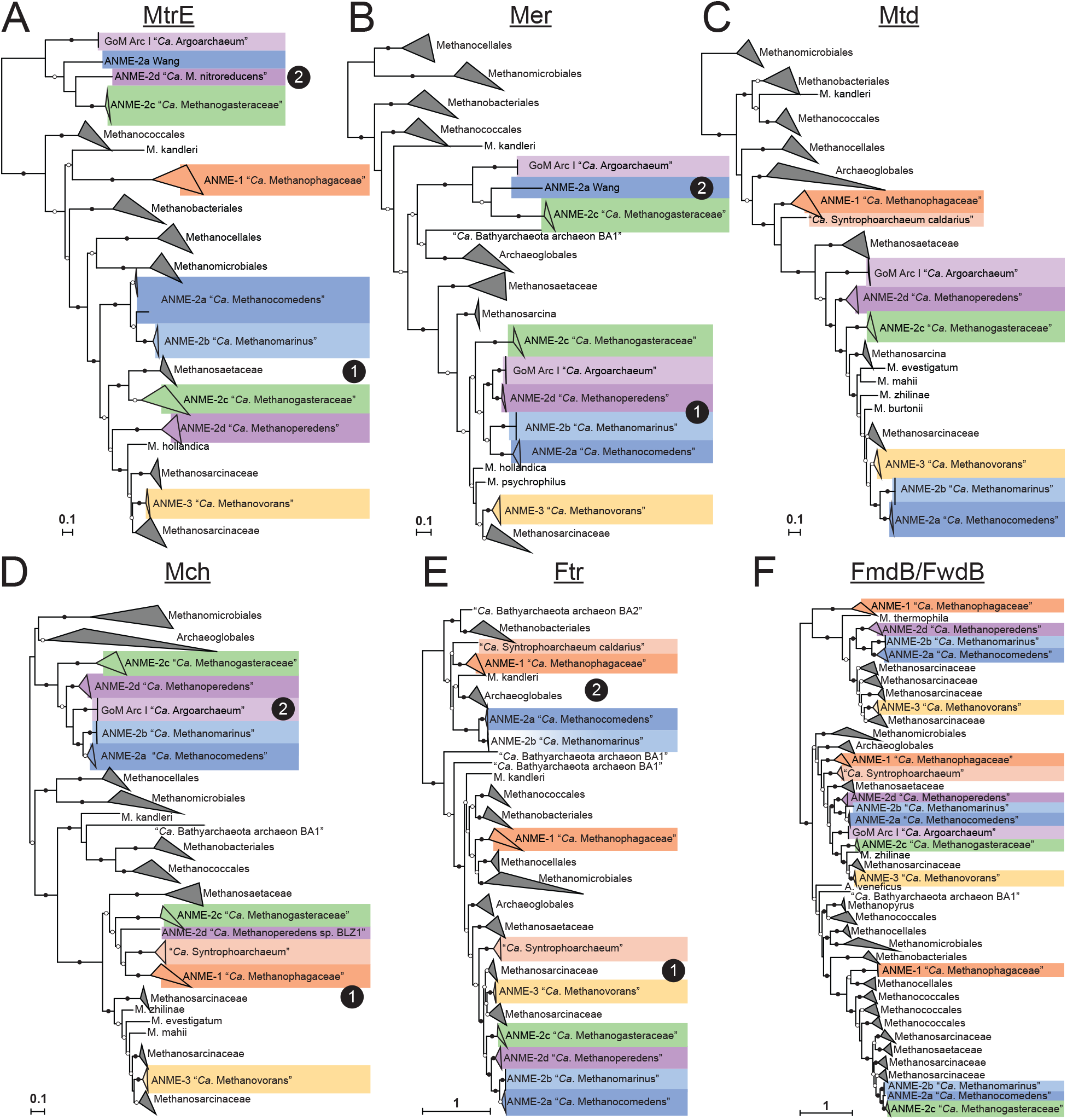
Phylogeny of enzymes in the methanogenesis pathway. Phylogenetic trees constructed from protein sequences of enzymes involved in the methanogenesis pathway in ANME and related archaea. Mcr phylogeny is presented in Fig 1. Numbers next to clades indicates whether the cluster is closely related to those found in *Methanosarcinaceae* (1) or are distantly related homologs (2), matching labels in Fig 3. **A)** MtrE: N^5^-methyl-H_4_MPT:coenzyme M methyltransferase subunit E; **B)** Mer: methylene-H_4_MPT reductase; **C)** Mtd: F_420_-dependent methylene-H_4_MPT dehydrogenase; **D)** Mch: N^5^,N^10^-methenyl-H_4_MPT cyclohydrolase; **E)** Ftr: formylmethanofuran-H_4_MPT formyltransferase; **F)** FmdB/FwdB: formyl-methanofuran dehydrogenase subunit B, molybdenum/tungsten variety, respectively. Branch support values of 100% are labelled with closed circles, >50% with open circles. Tree scales represent substitutions per site. Tree construction parameters are found in the **Materials and Methods** section. Alignments and tree files can be found in **S1 Data.**

The fourth step of the pathway is catalyzed by the F_420_-dependent N^5^,N^10^-methylene-H_4_MPT dehydrogenase (Mtd) enzyme (**Fig 3**). Only a single copy of the gene encoding Mtd was found in any genome, and the phylogeny of the predicted protein sequence is largely congruent with the genome phylogenies (**Fig 4C**). The fifth step in the pathway is catalyzed by N^5^,N^10^-methenyl-H_4_MPT cyclohydrolase (Mch). The analysis of early fosmid libraries revealed a gene on an ANME-2c-assigned fosmid encoding Mch (8), which was highly divergent from an Mch located on an ANME-2-assigned fosmid from a previous study (6). This led the authors to question whether the Mch had diverged rapidly between ANME-2c and different ANME-2 groups, or whether multiple Mch copies were present as paralogs within ANME-2c genomes. Interestingly, ANME-2a, 2b, 2c and 2d all share a well-supported monophyletic group of Mch genes (Mch2) that are very different from those of closely related methanogens (Mch1) (**Fig 4D**). The Mch2 cluster corresponds to the gene identified as being closely related to Mch found in *Archaeoglobus* (8). The ANME-2c genomes analyzed here also contain a copy of Mch1 that is closely related to those found in the methanogenic *Methanosarcinaceae*, which corresponds to the gene found in the first set of ANME-2-assigned fosmids (6). Apparently ANME-2c contain both copies of this gene, while the ANME-2a, 2b, and 2d only contain the divergent Mch2. ANME-3 contain a copy very similar to those of their close methanogenic relatives.

The reaction catalyzed by Mch is a curious step of the methanogenesis pathway to have a strongly supported, ANME-specific clade of enzymes. The cyclohydrolase reaction is thought to occur essentially at equilibrium (37), so it is unclear what evolutionary pressure would result in such a stark difference between Mch2 enzymes in some ANME-2 and their closely related methanogenic relatives. The pterin moiety of the H_4_MPT analog could vary between ANME and the methanogens in the *Methanosarcinaceae*. At least five pterins are known to be found in H_4_MPT analogs in methanogenic archaea: methanopterin, sarcinopterin, tatiopterin-I, tatiopterin-O and thermopterin (46). However, this level of sequence variation is not observed in other enzymatic steps of the pathway, which might be expected if the divergent Mch2 was the result of a significantly different form of C1-carrier in ANME.

Divergent paralogs of formylmethanofuran-tetrahydromethanopterin N-formyltransferase (Ftr) were found in ANME-1, 2a and 2b. These Ftr2 clustered together in the phylogenetic tree with Ftr genes from *Archaeoglobales* and deeper branching hydrogenotrophic methanogens such as *Methanopyrus kandleri* and *Methanothermobacter marburgensis* (**Fig 4E**). These archaea all contain both Ftr1 and Ftr2, and only the Ftr1 versions have been biochemically characterized to our knowledge (47). In the cases where transcriptomic information is available, Ftr1 is more highly expressed in ANME, and is therefore expected to be the dominant version utilized in ANME energy metabolism under the AOM conditions tested (**S3 Data**).

The seventh and final step of the methanogenesis pathway is carried out by formyl-methanofuran dehydrogenase (**Fig 3**). Two major variants of formylmethanofuran dehydrogenase are present in methanogens that contain either tungsten or molybdenum metal centers in their active sites (Fwd and Fmd, respectively), and multiple paralogs of both can be found in methanogens such as *Methanosarcina acetivorans* (48). Based on homology to versions of these enzymes in *M. acetivorans*, it appears that ANME largely contain the Fwd version, which have been detected in proteomic analyses of methane seeps (49) (**Fig 4F**). Some members of the ANME-2a, 2b, and 2c have the Fmd version as well, and in the ANME-2c the genes encoding Fmd and Fwd occur together in a single large gene cluster. As was observed in previous ANME-1 genomes, the FwdFG subunits are not present in any ANME-1 (6, 7, 12).

Based on the above observations we conclude that the transition from methanogenesis to methanotrophy required relatively little biochemical novelty within the central C1-carrying pathway of ANME energy metabolism. The loss of Mer in ANME-1 (**Fig 3**) remains the single major variation to the central C1-carrying pathway of ANME energy metabolism. Some paralogs of other steps in the pathway exist, but with the exception of Mch2 in some ANME-2, these are less well conserved and less well expressed than those previously found in methanogenic archaea. While McrA in ANME-1 is slightly incongruous with its genome phylogeny, and was found to bind a modified F_430_ cofactor (50), we see little evidence for significant changes in ANME-2 or 3, suggesting these modifications are not necessary for using Mcr to activate methane. These results are broadly consistent with biochemical studies that have demonstrated the reversibility of the enzymes in this pathway (36–38), suggests that there has likely been little specialization in terms of their directionality during the evolution of the ANME archaea.

#### Differing roles for MetF in ANME archaea

Methylenetetrahydrofolate reductase (MetF) has been proposed to act in the third step as a replacement for Mer in ANME-1 (10, 42) and is expressed at similar levels as other genes in the reverse methanogenesis pathway (9) (**S3 Data**). MetF is structurally similar to Mer and completes the same step of the Wood-Ljungdahl pathway in bacteria but uses NADPH as the electron donor rather than F_420_H_2_, and interacts with C1-bound tetrahydrofolate (H_4_F) instead of tetrahydromethanopterin (H_4_MPT).

MetF is not only found in ANME-1, but also in other ANME groups and methanogenic members of the *Methanosarcinaceae*, where it is expected to function as a methylene-H_4_F-interacting enzyme involved in anabolic processes (51). Since the potential Mer/MetF switch in ANME-1 appears to be the only significant modification to the central carbon oxidation pathway in ANME we investigated the distribution, phylogenetic placement, and genomic context of MetF in ANME in order to try and better understand the evolutionary history of these proteins. MetF homologs found in ANME-1 were clearly very different at the sequence level from those found in other ANME genomes (**Fig 5A**), and, interestingly, the ANME-2c universally lack MetF of either type. All ANME MetF are found next to a MetV gene which is a common feature of MetF in acetogenic bacteria, and evidence suggests a complex forms between the proteins encoded by these two genes (52).

**Fig. 5.**
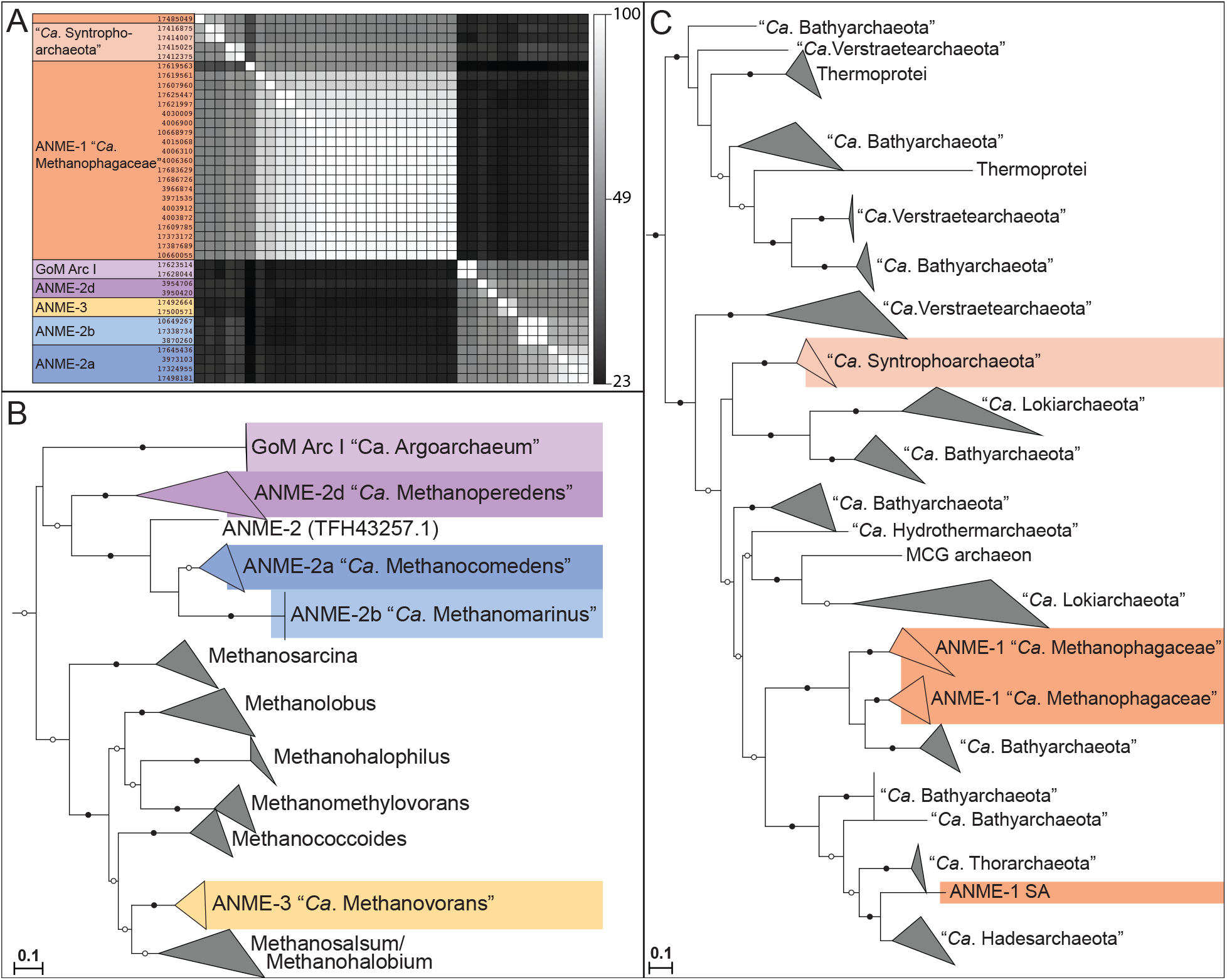
MetF in ANME Archaea. (**A**) Amino acid sequence identity of MetF homologs found in ANME, “*Ca.* Argoarchaeum” and “*Ca.* Syntrophoarchaeum”. ANME-1 and “*Ca.* Syntrophoarchaeum” form one cluster based on sequence similarity, while ANME-2a, 2b, 2d, 3 and “*Ca.* Argoarchaeum” form a second. Grayscale values represent percent identity. Sequences similar to the ANME-2/3 or ANME-1 clusters were retrieved via BLAST search of the NCBI nr database and used to construct phylogenetic trees of these two groups. (**B**) ANME-2, 3 and “*Ca.* Argoarchaeum” cluster together with closely related members of the *Methanosarcinaceae*. (**C**) ANME-1 and “*Ca.* Syntrophoarchaeum” form a polyphyletic group within a diverse group of sequences derived from MAGs of uncultured archaea. Notably the ANME-1 sp. SA is significantly different than the rest of the ANME-1. Roots for both trees lead to closely related MetF sequences from bacteria. Branch support values of 100% are labelled with closed circles, >50% with open circles. Tree scales represent substitutions per site. Tree construction parameters are found in the **Materials and Methods** section. Alignments and tree files can be found in **S1 Data**.

Two phylogenetic trees were constructed with sequences of each of these MetF groups along with the most closely related homologs in the NCBI NR database (**Fig 5B and 5C**). The ANME-1 MetF homologs branched together with a diverse group of proteins found exclusively in uncultivated archaeal genomes, most of them identified as “*Ca.* Bathyarchaeota”, “*Ca.* Verstraetearchaeota” and various members of the “*Ca*. Asgardarchaeota”. The MetF from the other ANME were all found within a monophyletic group containing other closely related *Methanosarcinaceae*. MetF and MetV in the *Methanosarcinaneae* are found in gene clusters with other H_4_F-interacting enzymes that carry out important C1 reactions in biosynthesis, which led to the conclusion that H_4_F is used for biosynthesis in the *Methanosarcinaceae* (51). This clustering of anabolic C1 genes is preserved in many of the ANME-2 and 3, and we infer a similar function. For a detailed discussion of C1 anabolic metabolism in ANME, see below.

The phylogeny of MetF proteins in ANME is best explained by two separate acquisitions of these genes. MetF in ANME-2 and 3 were likely acquired in the common ancestor of these organisms and the other *Methanosarcinaceae* along with other enzymes to utilize H_4_F-bound C1 moieties for the purpose of biosynthesis. Based on its distribution in uncultured archaea of different phyla, as well as the paucity of other H_4_F-interacting proteins in ANME-1, there is good support for a different, possibly catabolic, function of MetF in ANME-1. Due to the structural similarity between H_4_F and H_4_MPT (53), it is possible that the ANME-1-type MetF has evolved to react with C1 moieties bound to H_4_MPT instead of H_4_F. A switch between H_4_F and H_4_MPT as a carbon carrier is not unprecedented, as serine hydroxymethyl transferase (GlyA) has different versions specific to either H_4_F (51) or H_4_MPT (54, 55). Additionally, MtdA in methylotrophic bacteria has been shown to react with either H_4_F or H_4_MPT (56).

### Energy metabolism phase 2: Cytoplasmic electron carrier oxidation and energy conservation

After methane is oxidized to CO_2_ in the pathway described above, four electrons will be found on two molecules of F_420_H_2_, two electrons on reduced ferredoxin (Fd^2-^), and two electrons on the reduced forms of coenzyme M and coenzyme B (CoM-SH, CoB-SH). If the proposed Mer/MetF switch in ANME-1 is correct, then it is possible that MetF may have produced a reduced NADPH instead of one of the F_420_H_2_ (**Fig 2**). Importantly, these C1 oxidation reactions do not produce energy for the cell and will in fact consume sodium motive force at the step catalyzed by Mtr. This investment of energy helps drive the reaction in the oxidative direction, and the effect of energy investment can be seen in the decreased redox potential of the eight methane-derived electrons: the CH_4_/CO_2_ redox couple has a standard state midpoint potential of -240mV, and the average midpoint potential of the electrons once transferred to their cytoplasmic electron carriers is approximately -340mV (two on Fd^2-^ (-500mV), four on F_420_H_2_ (-360mV) and two on CoM-SH/CoB-SH (-145mV)) (37). The next phase of ANME energy metabolism, the re-oxidation of these reduced cytoplasmic electron carriers, must conserve sufficient energy to overcome the loss at Mtr and support growth.

This is very similar to the situation in the energy metabolism of methylotrophic methanogens in the *Methanosarcinaceae* which are close relatives of ANME-2 and 3. In these methanogens methyl groups are transferred from substrates such as methanol or methylamines onto CoM via substrate-specific methyltransferases. The methyl group is subsequently oxidized to CO_2_ via a reversal of the first six steps of the methanogenesis pathway, consuming sodium motive force at Mtr and producing two F_420_H_2_ and one Fd^2-^ that must be re-oxidized coupled to energy conservation. In most cases, these electrons are transferred onto membrane-bound methanophenazine as an intermediate electron carrier in processes that lead to ion motive force generation. This ion motive force is then used to produce ATP via ATP synthase. The only oxidation reaction in ANME which does not normally occur in any methanogen is the production of the heterodisulfide (CoM-S-S-CoB) from the free sulfides CoM-SH and CoB-SH. Many of the strategies of energy conservation coupled to cytoplasmic electron carrier oxidation that have been characterized in the methylotrophic methanogens appear to be conserved in the ANME archaea.

#### F_420_H_2_ oxidation

F_420_H_2_ oxidation for the purpose of energy conservation in methylotrophic methanogens can occur via the F_420_H_2_:methanophenazine oxidoreductase complex (Fpo) or the F_420_-reducing hydrogenase (Frh) (**Fig 6A**). F_420_H_2_ oxidation by the cytoplasmic Frh produces H_2_ which diffuses out of the cell and is subsequently oxidized on the positive side of the membrane by a membrane-bound hydrogenase (Vht) (57). Our comparative analysis of ANME genomes does not support this mechanism of electron flow given the lack of both the NiFe hydrogenase subunit of Frh and Vht-like hydrogenases.

**Fig 6.**
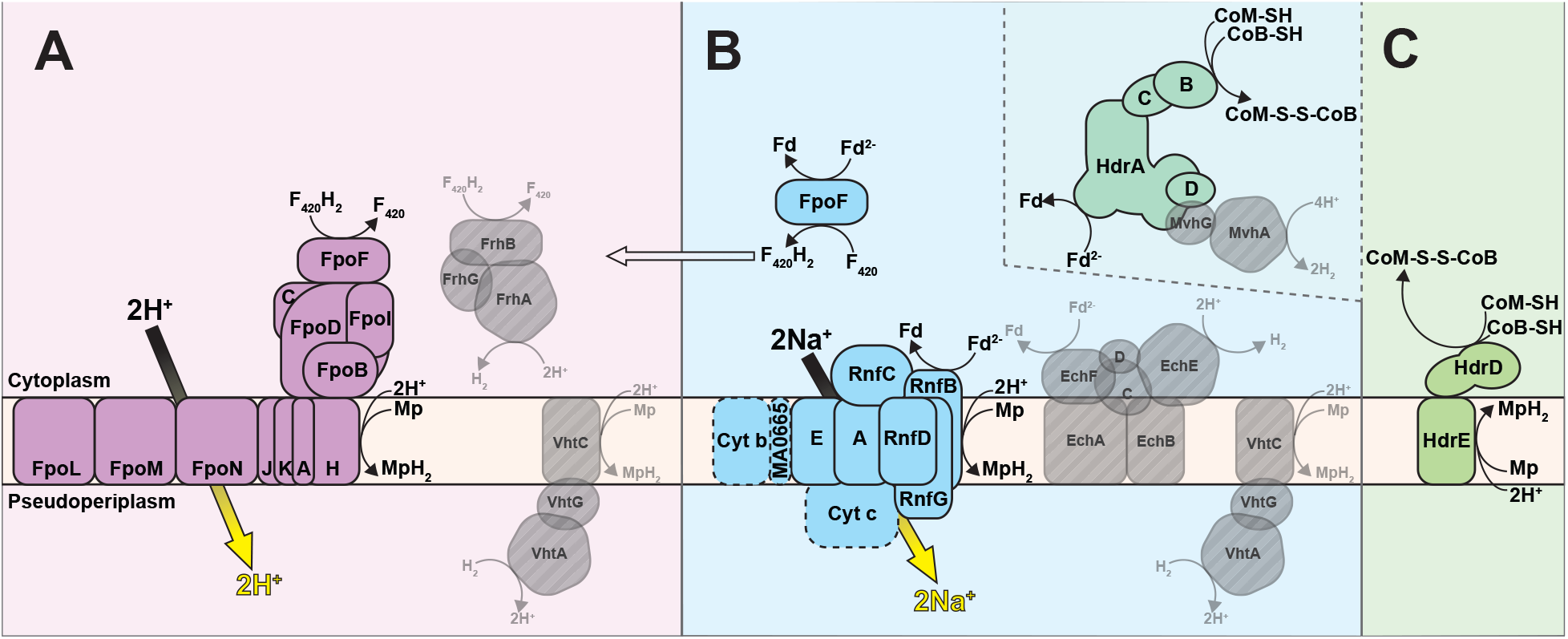
Cytoplasmic electron carrier oxidation. Some energy conservation systems discovered in methanogenic archaea are conserved in ANME archaea (colored fill) while others appear absent (transparent gray with diagonal line fill). (**A**) F_420_H2 oxidation is coupled to proton translocation in methylotrophic methanogens via the Fpo/Fqo complex or by the production of H_2_ by Frh and subsequent oxidation by Vht. In either case, electrons are ultimately deposited on MpH_2_. Neither Frh or Vht complexes have been observed in any ANME genomes analyzed here. (**B**) Fd^2-^ oxidation can be coupled either to sodium motive force or proton motive force in methylotrophic methanogens. The Rnf complex catalyzes Fd^2-^:Mp oxidoreductase reaction coupled to sodium translocation and is found in a number of methanogens and ANME. The ANME-2c contain most of the complex, but lack the cytochrome *c*, cytochrome *b* and MA0665 subunits, so their activity is difficult to predict. Ech and Vht can combine to produce net proton translocation via H_2_ diffusion in methylotrophic methanogens, but neither complex is found in ANME. FpoF can catalyze a Fd^2-^:F_420_ oxidoreductase reaction and F_420_H_2_ could then pass through the Fpo/Fqo complex. Various HdrABC complexes are present in all ANME genomes, and could in principle oxidize Fd^2-^ ad CoM-SH/CoB-SH through a reversal of electron bifurcation reaction. The electron acceptor in this process is likely to not be H_2_ in most ANME groups due to the absence of MvhG and MvhA. (**C**) Besides the HdrABC complexes mentioned above and second possible CoM-SH/CoB-SH oxidation strategy would be a reversal of the HdrDE reaction found in methylotrophic methanogens. In ANME the reaction would have to proceed in the direction illustrated, and therefore would dissipate proton motive force by consuming a proton on the positive side of the membrane. For presence/absence of these systems in ANME genomes analyzed here see **S6 Fig.**

This leaves Fpo as the much more likely candidate for F_420_H_2_ oxidation in ANME. Fpo is a homolog of respiratory complex I, and in the *Methanosarcinaceae* it couples the transfer of electrons from F_420_H_2_ to the membrane soluble electron carrier methanophenazine with the translocation of protons across the cell membrane. In sulfate-reducing *Archaeoglobales* the homologous F_420_H_2_:quinone oxidoreductase complex (Fqo) utilizes a membrane-soluble quinone acceptor instead of methanophenazine (58, 59). These complexes are conserved across all ANME clades (**S6 Fig**), and phylogenetic analysis of Fpo shows that homologs from ANME-2a, 2b, 2c and 3 are most similar to the *Methanosarcinaceae*, suggesting they also may utilize methanophenazine. The homologs from ANME-1 were distinct from those of the other ANME clades and were most similar to the Fqo described from *Archaeoglobales* as previously reported (6) (**S7 Fig**). Consistent with this possibility ANME-1 genomes also contain homologs of the futalosine pathway used for menaquinone biosynthesis that are absent in the other marine ANME clades. This suggests that ANME-1 use a quinone as their membrane-soluble electron carrier as was previously suggested (42). In either case, Fpo and Fqo are expected to be important points of F_420_H_2_ oxidation and membrane energization in the ANME archaea.

#### Ferredoxin oxidation

Similar to F_420_H_2_ oxidation, several mechanisms are known for coupling ferredoxin oxidation to energy conservation in methanogens. Four major pathways have been proposed: 1) a hydrogen cycling mechanism using energy conserving hydrogenase (Ech) (57), 2) oxidation with a modified version of the Rhodobacter nitrogen fixation (Rnf) complex (60), 3) an Fpo-dependent pathway (61), and 4) a heterodisulfide reductase (Hdr)-mediated electron confurcation (62)(**Fig 6B**). The Ech model is easily ruled out by the lack of Ech homologs in all marine ANME genomes, although these are present in the freshwater ANME-2d genomes (19, 20).

Genomes from many members of the ANME-2a, 2b, 2c and 3 contained homologs of the Rnf complex (**S6 Fig**). Rnf was first characterized in bacteria as an enzyme that performed the endergonic transfer of electrons from NADH to ferredoxin by dissipating sodium motive force (63). In methanogens however, Rnf is thought to couple the exergonic electron transfer from ferredoxin to methanophenazine with the endergonic translocation of sodium ions to the positive side of the cytoplasmic membrane. This activity in methanogens involves a novel multiheme cytochrome *c* subunit encoded in their Rnf gene clusters (60), along with a small conserved integral membrane protein (MA0665 in *M. acetivorans*). This enzyme complex operates in methylotrophic members of the *Methanosarcinaceae* that do not conduct hydrogen cycling, and in some cases is required for energy conservation during acetoclastic methanogenesis (64, 65).

Rnf was not found in the ANME-1 genomes analyzed here, with the exception of two genomes from a genus-level subclade recovered from a South African gold mine aquifer (SA) and from a hydrocarbon seep from the Gulf of Mexico (GoMg4) (**Table 1**; **S6 Fig**). Rnf gene clusters in ANME-2a, 2b and 3 contain homologs of the cytochrome *c* subunit and MA0665 normally found in methanogens, but surprisingly both were missing in the Rnf gene clusters from all ANME-2c and the two ANME-1. Homologs for these subunits were not identified elsewhere in any of these genomes. Based on the current information, it is unclear what reaction the ANME-2c and ANME-1 Rnf are performing since it has been demonstrated that this cytochrome *c* is involved in the electron transfer to methanophenazine (66). It is possible that Rnf retains the ability to transfer electrons from ferredoxin to a membrane bound electron carrier in these lineages, or alternatively they could function as ferredoxin:NAD^+^ oxidoreductase, as found in bacteria. It is unclear what role the latter function could play in our current model of ANME metabolism.

An additional ANME-specific modification to the *rnf* gene cluster is the inclusion of a membrane-bound b-type cytochrome in ANME-2a, 2b and 3. This type of cytochrome is generally involved in electron transfer between membrane-bound and soluble electron carriers, and has no closely related homologs in methanogens. If the protein encoded by this gene is incorporated into the Rnf complex, it could be of great importance to the flow of electrons in these groups. This observation is particularly striking in ANME-3 since their close methanogenic relatives in the *Methanosarcinaceae* lack this b-type cytochrome. This means that ANME-3 has acquired this subunit by horizontal gene transfer from an ANME-2a or 2b, or that all of the related methanogens have lost it. Whichever evolutionary scenario is correct, this represents an important ANME-specific modification to a key bioenergetic complex.

Because the majority of ANME-1 do not contain an Rnf complex an alternative mechanism is needed to explain how ferredoxin is recycled. One option is the Fpo-dependent mechanism proposed in *Methanosaeta thermophila* where ferredoxin is oxidized using an Fpo complex without the FpoF subunit (61). If this pathway operates in ANME-1 the reduced ferredoxin would donate electrons directly to the iron sulfur clusters found in the FqoBCDI subunits. Since FqoF homologs are encoded in the ANME-1 genomes, and are expected to be used in complete Fqo complexes to oxidize F_420_H_2_ as described above, this Fd^2-^ oxidation strategy would require some Fqo complexes to have FqoF bound, while others do not.

Another possibility would be for soluble FqoF to act as a Fd^2-^:F_420_ oxidoreductase, and then the subsequent oxidation of F_420_H_2_ by a normal Fqo complex (**Fig 6B**). This pathway has been proposed in *Methanosarcina mazei* based on the Fd^2-^:F_420_ oxidoreductase activity of soluble FpoF (67). In either of these Fpo/Fqo-dependent Fd^2-^ oxidation pathways, the oxidation of Fd^2-^ would result in the reduction of a membrane-bound electron carrier coupled to energy conservation in the form of proton translocation by the Fpo/Fqo complexes.

A final possible mechanism of Fd^2-^ oxidation is through soluble heterodisulfide reductase (Hdr)-mediated flavin-based electron confurcation. This process has some biochemical support for operating in this direction (62), but would constitute a reversal of the well accepted electron bifurcation mechanism used by many hydrogenotrophic methanogens. In the best characterized examples of this process, an enzyme complex of soluble Hdr and a hydrogenase (hdrABC-mvhADG) reduce ferredoxin and heterodisulfide with four electrons sourced from two hydrogen molecules (68). If reversed, this reaction could potentially oxidize Fd^2-^. As this mechanism would also involve CoM-SH/CoB-SH oxidation it is discussed in detail in the following section.

#### CoM-SH/CoB-SH oxidation

The last oxidation that needs to occur during this part of ANME energy metabolism is CoM-SH/CoB-SH oxidation to the heterodisulfide (CoM-S-S-CoB). This oxidation is the most energetically challenging because the relatively high midpoint potential of the heterodisulfide (E_0_’= -145mV) rules out most methanogenic electron carriers as acceptors without the input of energy to force electrons “uphill” to a lower redox potential. It also represents the second net reaction in the ANME energy metabolism that does not occur during canonical methanogenesis (the first step being methane activation through Mcr). In methylotrophic methanogens heterodisulfide is the terminal electron acceptor for the six electrons that have passed through F_420_H_2_ and Fd^2-^ discussed in the previous sections.

Two enzyme systems from methanogens could potentially produce heterodisulfide by running in reverse: soluble HdrABC complexes using a confurcation mechanism as mentioned above, or HdrDE, a membrane-bound system which would deposit electrons on methanophenazine (**Fig 6C**). ANME-2a, 2b, 2c, 2d, and 3 genomes contained HdrDE genes similar to ones from closely related methanogens. The reaction carried out by this complex in methanogens is expected to be readily reversible, with electrons from CoM-SH/CoB-SH oxidation being deposited on methanophenazine. This electron transfer reaction is slightly endergonic at standard state due to the redox potential of methanophenazine (E_0_’ = -165mV) being lower than that of heterodisulfide by 20mV. This reaction may be exergonic under physiological concentrations of oxidized and reduced species. Alternatively, the slightly endergonic nature of the electron transfer could be overcome by a quinol loop-like mechanism, where two protons are released in the cytoplasm from CoM-SH/CoB-SH oxidation and two protons are consumed on the outer face of the cytoplasmic membrane to form reduced MpH_2_, thus dissipating proton motive force (69). Electron transfer from the cytoplasm to the outer face of the membrane occurs through the b-type cytochromes in the HdrE subunit. The HdrDE complexes are well expressed in the ANME-2a, 2c and 2d (**S3 Data**).

HdrDE gene clusters are absent in all ANME-1 genomes. This, combined with the lack of Rnf and Ech, suggests that ANME-1 in particular could rely on HdrABC complexes for both CoM-SH/CoB-SH and ferredoxin oxidation. Since its discovery just over ten years ago (70), flavin-based electron bifurcation and confurcation have been shown to be a key energy conversion point in many anaerobic metabolisms, with a great diversity of different electron donors and acceptors (71). In the complete HdrABC-MvhADG complex from methanogens the endergonic reduction of Fd (E_0_’ = -500mV) with H_2_ (E_0_’ = -420mV) is driven by the exergonic reduction of heterodisulfide (E_0_’ = -145mV) with H_2_.

In most cases a strict reversal of H_2_ electron bifurcation is not possible in ANME due to a lack of MvhAG genes that encode the NiFe hydrogenase large and small subunits. The exceptions are two small subclades within ANME-2c and ANME-1 that contain NiFe hydrogenases similar to Mvh. These two subclades contain genomes recovered from different environments; a South African gold mine (SA) and the Gulf of Mexico (GoMg4) for ANME-1, versus a hydrocarbon seep off of Santa Barbara, CA (COP2) and a mud volcano from the Mediterranean (AMVER4-31) for ANME-2c. The majority of ANME-2c and ANME-1 however lack these genes, and they are completely absent in ANME-2a, 2b and 3, suggesting confurcation of electrons from Fd^2-^ and CoM-SH/CoB-SH to H_2_ is not a dominant process in most ANME lineages.

#### Novel gene clusters encoding electron bifurcation/confurcation complexes

While a strict reversal of hydrogen-dependent electron bifurcation seems unlikely in ANME, the potential involvement of alternative bifurcation/confurcation reactions in ANME metabolism is supported by the broad distribution and conservation of unusual HdrABC homologs, even in those ANME genomes containing HdrDE. Alternative bifurcation reactions have been proposed in methanogens, where formate or F_420_H_2_ can serve as the electron donor in place of hydrogen. In the case of formate, MvhA and G are replaced by the F_420_H_2_-dependent formate dehydrogenase genes FdhAB (72, 73). In the case of F_420_H_2_, certain HdrABC complexes might be able to interact with F_420_H_2_ without any additional protein subunits (62, 74).

A comparison of ANME HdrA homologs to crystal structures of the entire HdrABC-MvhADG complex purified from *Methanothermococcus wolfeii* (75), reveal some stark differences in how these complexes may facilitate electron bifurcation. HdrA in methanogens normally consists of four domains, an N-terminal domain with an iron sulfur cluster, a thioredoxin reductase domain which binds the bifurcating FAD, and two ferredoxin domains (**Fig 7**). The MvhAG hydrogenase feeds electrons through MvhD and the C-terminal ferredoxin domain to the FAD where they are bifurcated, one pair passing up through HdrBC onto CoM-S-S-CoB, and the other through the inserted ferredoxin domain to a free soluble ferredoxin. Interestingly, the heterohexameric HdrABC-MvhADG forms dimers, and one of the cysteine ligands for the N-terminal iron sulfur cluster comes from the thioredoxin-reductase domain of the other copy of HdrA (Cys197), indicating an obligate dimeric nature of the complex in *M. wolfeii* (**Fig 7C**).

**Fig 7.**
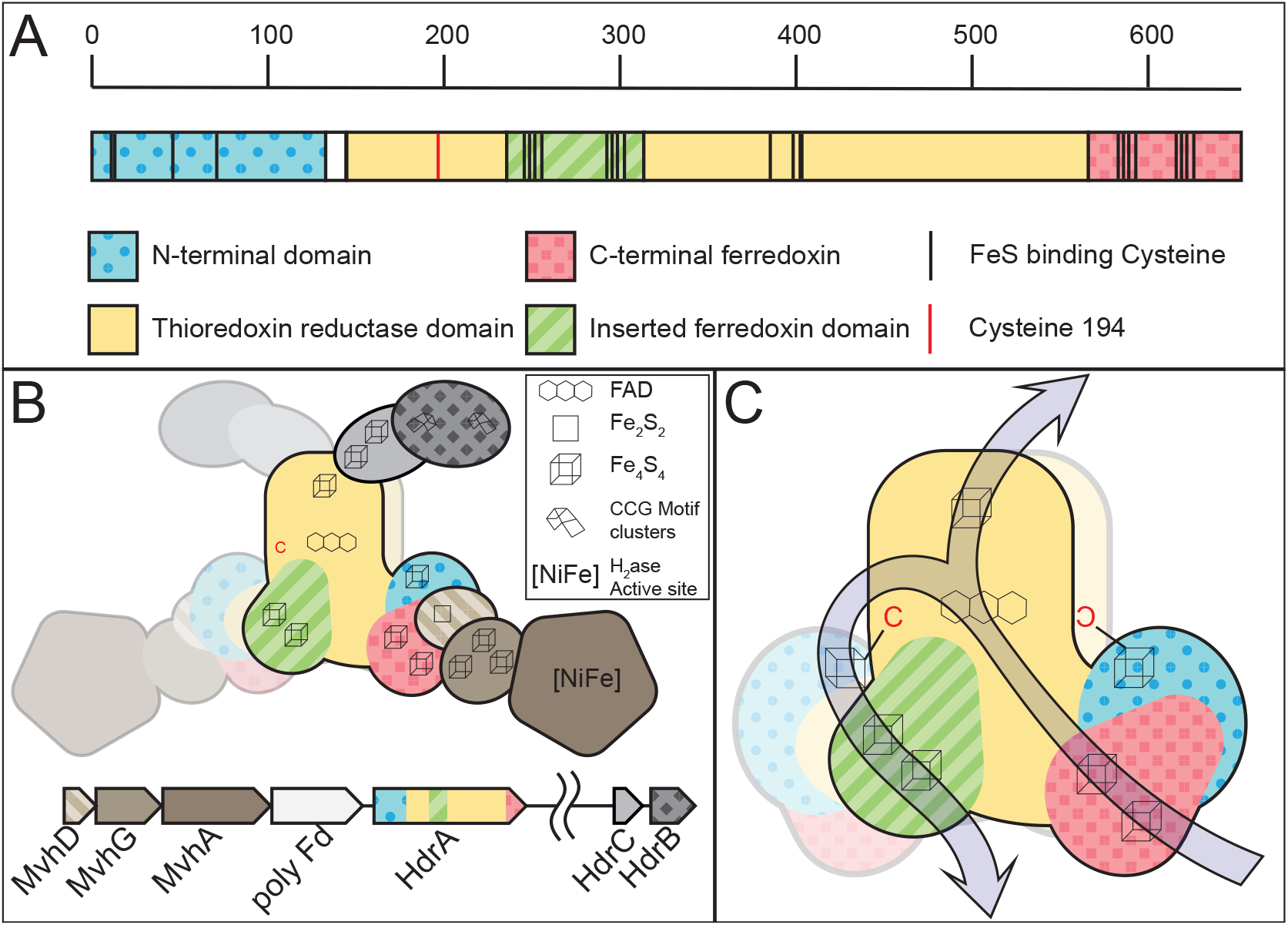
HdrABC structure overview. Depiction of the primary structure of HdrA and the quarternary structure of the HdrABC-MvhADG complex based on the structure from *M. wolfeii*. (**A**) HdrA can be broken down into four domains, the positions of these domains and key iron-sulfur cluster binding cysteines are illustrated, scale denotes amino acid position in the *M. wolfeii* sequence. (**B**) Quarternary structure of the entire HdrABC-MvhABG complex illustrating the dimeric structure. Metal cofactors involved in the oxidation/reduction of substrates or electron transport through the complex are highlighted. (**C**) Detail of HdrA domain structure highlighting cofactor position and proposed electron flow from MvhD in through the C-terminal ferredoxin, bifurcation through the FAD cofactor, with two electrons flowing out through HdrBC via the Thioredoxin reductase domain’s FeS cluster, while two other electrons flow out through the inserted ferredoxin domain, presumably to free ferredoxin (Fd^2-^). Importantly for the proposed heterodimeric HdrA discussed here, this latter electron flow passes through the FeS cluster bound through a combination of Cys residues in the N-terminal domain, combined with a single Cys from the other HdrA subunit (Cys197 highlighted in red).

By aligning the ANME HdrA paralogs and comparing the presence of domains, conserved residues, sequence similarity, and their genomic context, we clustered the dominant HdrA homologs into 14 groups (**Fig 8A, S8 Fig**). Most methanogen genomes contain 1-2 copies of *hdrABC* gene clusters, but ANME, and in particular ANME-1, have a greater abundance of HdrA homologs that exceed the number of *hdrBC* homologs (**S9 Fig**). Some of these HdrA homologs exceed 1000 amino acids in length. In comparison, HdrA from methanogens is usually 650 amino acids in length, with some slightly larger homologs occurring due to the fusion of *hdrA* and *mvhD* (76). The gene clusters containing HdrA homologs often contained HdrBC and MvhD as expected, but more unexpected was the co-occurrence of multiple copies of HdrA and the occasional presence of HdrD-like proteins (**Fig 8A, S10 Fig**). Although HdrD is a fusion of HdrB and HdrC, it is not common to find the fused form in gene clusters with HdrA in methanogens. Distant homologs of the F_420_-dependent formate dehydrogenase FdhAB were also found in HdrA gene clusters in all ANME groups. The gene cluster containing HdrA2 and 3 along with FdhAB-like genes in ANME-2a was discussed in detail in one of the earliest studies of ANME fosmid libraries (8), and the significance of this cluster has now been substantiated by its broad conservation as well as reasonably high transcription levels (**S3 Data**).

**Fig 8.**
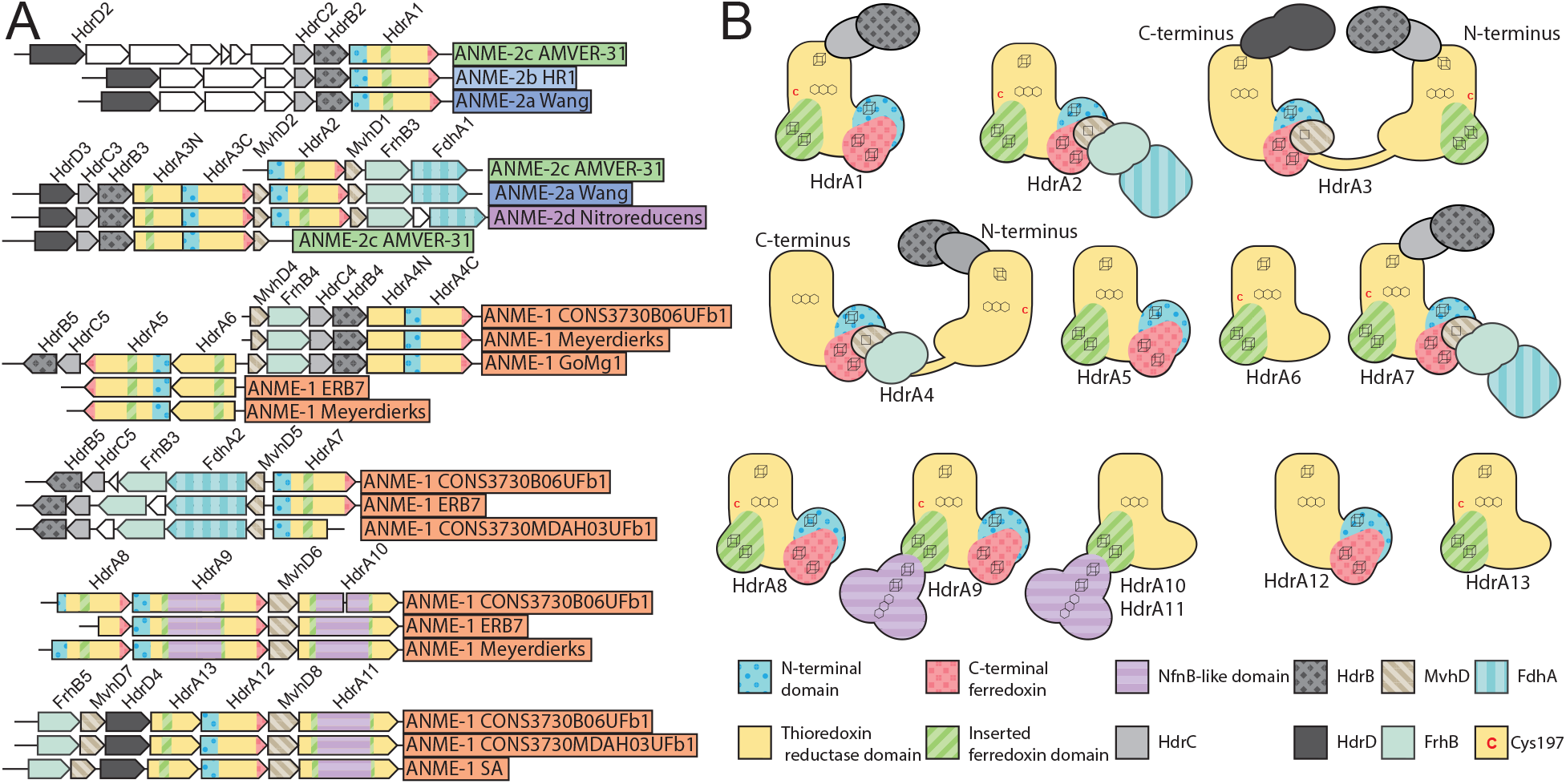
Hdr Operons and Domain heterogeneity. (**A**) Examples of gene clusters containing HdrA genes from select ANME genomes. HdrA paralogs present in ANME have extensive modification to the domain structure as compared to the HdrA crystalized from *M. wolfeii* (see Fig 7 for details of HdrA structure). These domains and associated protein subunits are illustrated with the gene context and orientation. (**B**) Illustration of conserved domains and cofactor binding residues in the 13 HdrA clusters defined here. All HdrA appeared to retain residues responsible for interaction with FAD, however the presence of FeS-binding cysteine residues, and entire domains as defined on the *M. wolfeii* structure are variably retained. Importantly, tandem or fused HdrA appear to have complementary presence/absence of C-terminal ferredoxin domains and Cys197 suggesting the formation of a heterodimeric complex.

Many HdrA homologs in ANME were lacking various domains present in the crystalized complex from *M. wolfeii* (**Fig 8B**). An interesting pattern emerges in gene clusters with tandem HdrA genes (i.e. HdrA 5 and 6, or 12 and 13), in which one copy has the N-terminal domain but lacks the Cys197 ligand for the N-terminal FeS cluster in its thioredoxin reductase domain (Hdr5 and 12), while the second copy lacks the N-terminal domain but has Cys197 (Hdr6 and 13). It seems likely that these tandem HdrA genes form heterodimers, breaking the rotational two-fold symmetry found in the crystal structure from *M. wolfeii* (**Fig 7B**). This pattern of Cys197/N-terminal domain complementarity can also be seen in largest HdrA homologs 3 and 4, which are fusions of two HdrA genes. In both cases the N-terminal HdrA contains the N-terminal FeS cluster-binding domain, but lacks Cys197, while the C-terminal HdrA contains Cys197 and no N-terminal domain, suggesting these HdrA fusions within a single continuous peptide may act as their own heterodimeric partners. Evidence for symmetry breaking in HdrABC complexes has recently been demonstrated in *Methanococcus maripaludis,* where a hydrogenase and a formate dehydrogenase can be simultaneously bound to dimerized HdrABC (77). This appears to be the result of HdrABC’s ability to bind either hydrogenase or formate dehydrogenase in these methanogens, resulting in a mixture of subunits, whereas in the ANME case this asymmetric form may be imposed by a heterodimer of two HdrA paralogs.

Another remarkable HdrA modification is a 500 amino acid insertion in the ferredoxin domain of HdrA 9, 10, and 11, present only in some ANME-1 genomes (**Fig 8**). This insertion shares 45% amino acid sequence identity to the NfnB subunit of the bifurcating NADH-dependent reduced ferredoxin:NADP oxidoreductase crystalized from *Thermotoga maritima* (78). NfnB binds the b-FAD cofactor thought to be responsible for bifurcation in the NfnAB complex, potentially giving these HdrA-NfnB genes two electron bifurcation sites, and suggests NADPH as a possible electron carrier in these compounds.

Based on the extensive conservation of FdhAB subunits in Hdr gene clusters in nearly all ANME groups it is tempting to speculate that these enzymes may have an important role in CoM-SH/CoB-SH oxidation through electron confurcation. A reversal of electron bifurcation from formate would be a reasonable reaction to expect in ANME metabolism, as formate is a common syntrophic intermediate. Based on the ratio of formate to bicarbonate in the cell one could envision the midpoint potential to lie appropriately between that of Fd^2-^ and CoM-SH/CoB-SH to receive electrons from both. FdhA belongs to the molybdopterin oxidoreductase family, members of which act on many different substrates including formate, nitrate, and DMSO among others in assimilatory and dissimilatory processes (79). The FdhA homologs from ANME genomes are often annotated as formate dehydrogenases and retain a conserved cysteine ligand for binding the molybdenum atom (sometimes a selenocysteine in other organisms). However, they are very distantly related to any biochemically characterized enzymes, making it difficult to assign their substrate with any level of certainty. A very closely related gene cluster with HdrA and FdhAB can be found in the genomes of the methylotrophic methanogen genus *Methanolobus*, potentially providing a good opportunity to study the substrate specificity of this specific group of molybdopterin oxidoreductases in a pure culture organism. These are the only methanogens that encode this type of Hdr gene cluster, and it is worth noting in this context that the *Methanolobus* are incapable of using formate as a methanogenic substrate (80).

Another possibility is confurcation of Fd^2-^ and CoM-SH/CoB-SH electrons to F_420_H_2_. FdhB and homologous proteins carry out oxidoreductase reactions with F_420_ (81) in F_420_-dependent hydrogenases (FrhB), formate dehydrogenases (FdhB), F_420_-dependant sulfite reductase (Fsr) and the Fpo complex (FpoF). The observation of FdhB homologs in gene clusters with HdrA in ANME-2d lead to the suggestion that these complexes could confurcate Fd^2-^ and CoM-SH/CoB-SH to reduce two molecules of F_420_, although this model proposes no role for the FdhA homologs present in the gene clusters (19). Some HdrA gene clusters only contain FdhB and not an FdhA homolog, however these are found only in ANME-1 genomes (**Fig 8A, S10 Fig**).Additionally, there is evidence that certain HdrABC complexes can interact with F_420_ without any additional subunits (62, 74). If such a reaction were to produce F_420_H_2_ in ANME, its re-oxidation through the Fpo complex could be coupled to energy conservation as has been previously proposed in *M. acetivorans* (82).

We can only speculate on the functions of all these modified HdrA paralogs, but it is clear that ANME have an exceptionally diverse potential of bifurcation/confurcation complexes at their disposal, and that the flow of electrons through these complexes will necessarily be different than in the traditional HdrABC-MvhADG complex due to domain gain and loss within the HdrA homologs, as well as the replacement of hydrogenase subunits with various other input modules. While any electron confurcation schemes through HdrABC is speculative at this point, they seem to be viable option for Fd^2-^ and CoM-SH/CoB-SH oxidation in ANME due to their widespread conservation across all ANME clades, particularly in ANME-1 that lack HdrDE and most known Fd^2-^ oxidation systems. Additionally, an examination of previously reported transcriptomic information indicates that these complexes are often expressed to levels on par with other components of the reverse methanogenesis pathway (**S3 Data**). A key question that remains is what substrate the FdhA homologs in ANME act upon.

### Energy metabolism phase 3: Genomic evidence for mechanisms of syntrophic electron transfer

The most enigmatic phase of marine ANME metabolism is the interspecies electron transfer to the sulfate-reducing partner; a process which appears to necessitate the formation of conspicuous multicellular aggregates of the two organisms (4, 83). The cytoplasmic electron carrier oxidation described in the previous section will result in 8 electrons on a combination of membrane-bound MpH_2_ or QH_2_ and possibly some soluble electron carrier formed through an electron confurcation reaction oxidizing Fd^2-^ and CoM-SH/CoB-SH. Based on energetic considerations and precedent from other known syntrophies, the hypothesized mechanisms for the AOM syntrophy have included the diffusive exchange of small molecules such as hydrogen, formate, acetate, methanol, methylamine (84), methyl-sulfides (85), zero-valent sulfur (86), and direct electron transfer using multiheme cytochrome *c* proteins (7, 87, 88). We assessed the genomic potential for each of these syntrophic electron transfer strategies across our expanded sampling of ANME clades.

#### Hydrogen transfer

Hydrogen transfer is one of the most common forms of syntrophic electron transfer (89). A classic mode of syntrophic growth involves hydrogenotrophic methanogens consuming H_2_ produced by the fermentative metabolism of a syntrophic partner. A direct reversal of the methanogenic side of this syntrophy is not possible as the majority of ANME genomes lack any identifiable hydrogenases with the notable exception of MvhA homologs in a small set of ANME-1 and 2c genomes as mentioned in the preceding section (**S10 Fig**). The first report of an ANME-1 genome from fosmid libraries contained a gene that appeared to encode an FeFe hydrogenase (7), however, homologs of this gene were not found in any of the other ANME-1 genomes analyzed here.

This lack of hydrogenases is consistent with the lack of energy-conserving hydrogenases in the genomes of their syntrophic sulfate-reducing bacterial partners from cold seeps (90) and previous experimental results showing that the addition of excess hydrogen does not inhibit AOM (91, 92). Hydrogen has been shown to stimulate sulfate reduction in AOM sediments suggesting that at least some portion of the sulfate-reducing community can utilize this electron donor (84, 91, 93). In the case of the syntrophic thermophilic ANME-1-“*Candidatus* Desulfofervidus auxilii” consortium, hydrogen amendment suppresses growth of ANME-1, because the “*Ca*. D. auxilii*”* can grow alone on hydrogen and stops investing in electron transferring structures (94). Together, genomic and physiological data suggests that the vast majority of ANME are incapable of producing hydrogen as an electron shuttle.

#### Formate transfer

Formate is another possible small molecule intermediate commonly involved in syntrophic electron transfer (95), and was predicted to be a possible intermediate in the ANME-SRB syntrophy based on early modeling studies (96). As described above FdhAB-like genes in association with Hdr-containing gene clusters are broadly distributed in ANME and could potentially be used to produce formate from Fd^2-^ and CoM-SH/CoB-SH and are relatively highly expressed (**S3 Data**) but their substrate is currently unknown. In addition to these complexes, five of the ANME-1 genomes contained a membrane-bound formate dehydrogenase with signal sequences predicted to mediate their secretion to the positive side of the cytoplasmic membrane. One of the ANME-1 genomes containing this formate dehydrogenase is based on early fosmid libraries that were analyzed extensively, and it was suggested to play a role in electron transfer, depicted as either being free-floating in the pseudoperiplasm, or anchored to the membrane via a *c*-type cytochrome (7). Interestingly, the gene cluster encoding these formate dehydrogenases in four other ANME-1 genomes also contain DmsC-like genes that traditionally serve as membrane anchor subunits for members of the molybdopterin oxidoreductase family. The presence of these genes in the other ANME-1 genomes makes it likely that this is how the complex is anchored and interacts with the ANME-1 membrane-bound quinol, since this is a fairly common pattern for members of this family of oxidoreductases (97). Interestingly these formate dehydrogenase genes are distributed very sporadically through the ANME-1 genomes instead of being confined to a specific subclade (**S10 Fig**).

This genomic evidence from different subclades provides a more plausible role for formate than hydrogen in ANME metabolism. However, like hydrogen, formate addition was not shown to inhibit AOM (84, 91, 92) which would be expected if it was the major electron donor to the sulfate reducing partner in those sediments. The addition of formate to AOM sediments and enrichment cultures of ANME-SRB consortia has also had mixed results in terms of stimulating sulfate reduction (84, 91, 93). Additionally, genes encoding FdhC, a member of the formate-nitrate transporter family thought to be responsible for formate transport in many formate-utilizing methanogens (98, 99) was absent from all ANME genomes.

#### Other soluble electron carriers

Acetate has been proposed as a syntrophic intermediate in sulfate-coupled AOM (7, 100), and one could easily envision an energy conservation strategy in ANME archaea whereby methane and CO_2_ are combined in a reversal of acetoclastic methanogenesis and energy is conserved via substrate-level phosphorylation through Pta and AckA as in acetogenic bacteria. Acetate thus produced could be used by the SRB partner for sulfate reduction. Two problems exist with this model that make it unlikely in any of the ANME groups represented here. First, Pta and AckA were absent from all ANME genomes. Second, this model requires that the two electrons transferred from methane to heterodisulfide (E_0_’ = -145mV) would be used for the reduction of CO_2_ which requires ferredoxin (E_0_’ = -500mV). This would mean the single ATP produced by substrate level phosphorylation would have to overcome the sodium motive energy lost in the Mtr step, force electrons energetically uphill by ∼350mV from the free disulfides onto ferredoxin, and have energy left over for growth. Experimental addition of acetate to AOM enrichments has not been found to decouple the partners and inhibit methane oxidation, as would be expected for a soluble extracellular intermediate (84, 91, 92).

Methylsulfides were suggested to be a potential intermediate in AOM (85), but as noted above the methyltransferase genes for methylsulfides, methanol, and methylamines were not recovered in any ANME genomes, and experiments have not found methylsulfide-stimulated sulfate reduction in ANME-SRB enrichments (91). Zero-valent sulfur has also been suggested as an intermediate in marine AOM using an ANME-2 enrichment (86). However, in vitro studies with other ANME consortia did not show a similar response (88, 93) and recent investigations into sulfur utilizing genes in ANME found no evidence for dissimilatory sulfate reduction (17). If this process occurs, it is in a restricted group of ANME through a novel biochemical mechanism. Electron transfer is also possible via soluble organic shuttles. *Shewanella oneidensis* MR-1 was determined to utilize small molecule shuttles derived from menaquinone (101) or flavins (102) that help electrons pass through the external environment to their ultimate electron acceptor. The mechanism of producing these compounds is very poorly understood. Only recently a major menaquinone-like shuttle was identified and its biosynthesis elucidated (103). Such a shuttle-based electron transfer strategy could be potentially used for accepting electrons from the ANME Hdr complexes, but predicting the occurrence of these shuttling mechanisms from genomic information is not possible with our current understanding of these processes.

#### Direct interspecies electron transfer

If soluble electron carriers were produced by ANME, they could simply diffuse through the intracellular space between the ANME to the SRB without any specialized protein systems. Specific transporters or permeases would only be needed if the compounds produced in the ANME cytoplasm are not membrane permeable. On the other hand, electrons on MpH_2_/QH_2_ would require an oxidation system with no analog in methanogens. In cytochrome-containing methanogens the eventual electron acceptor for these membrane-bound electrons is the heterodisulfide located in the cytoplasm, whereas the terminal electron acceptor of marine ANME would be their partner SRB located outside of the cell.

This is a respiratory challenge that is similar to the one facing bacteria that carry out extracellular electron transfer (EET) to utilize insoluble metal oxides or electrodes as terminal electron acceptors. Instead of having a soluble terminal electron acceptor that can diffuse to the cytoplasmic membrane and be reduced by a terminal oxidase, these organisms make conduits for their electron transport chain to extend through the periplasm and outer membranes to interact directly with an electron acceptor in the extracellular space (104).

Various systems have been discovered in Gram-negative bacteria that perform EET. These generally consist of a quinol:cytochrome *c* oxidoreductase enzyme, small soluble cytochrome *c* intermediates between the inner and outer membranes, and a beta barrel/decaheme cytochrome *c* protein complex that is embedded in the outer membrane (**Fig 9A**)(105). In bacteria that form large conductive biofilms, the extracellular space is also densely packed with secreted *c*-type cytochromes which can form large conductive complexes (106) . Based on the similarity of the metabolic challenge facing ANME and EET-capable bacteria, as well as the discovery of analogs of bacterial EET systems in ANME genomes (7, 87, 88), an attractive hypothesis is that the ANME-SRB syntrophy is based on direct electron transfer between the syntrophic partners. However, in contrast to other EET-capable bacteria that use metals as terminal electron acceptors, the highly specific interspecies interaction observed in the AOM consortia may require the partner SRB to encode cognate electron-accepting systems.

**Fig 9.**
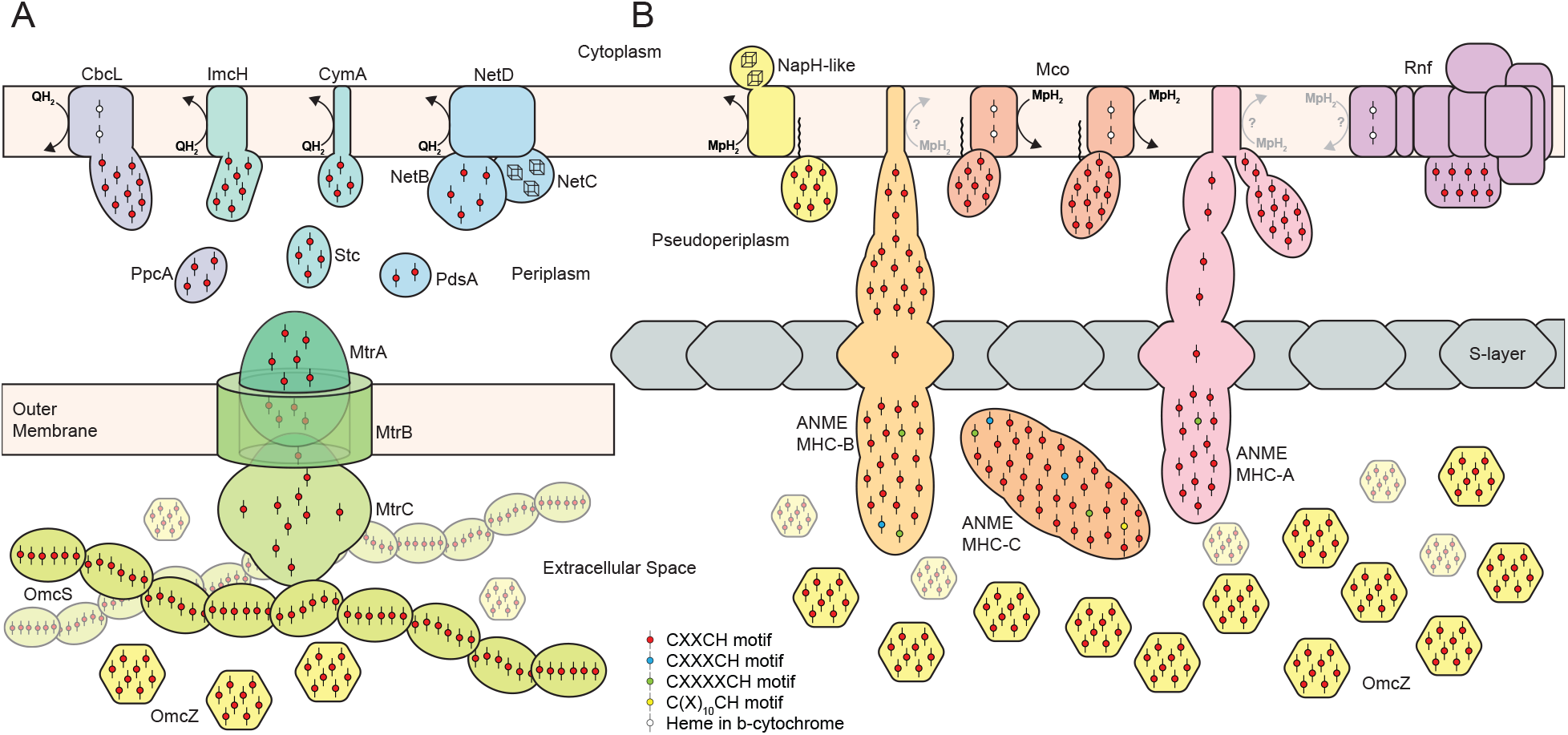
Overview of proposed extracellular electron transfer pathways. Comparison between EET systems known from Gram-negative bacteria and proposed analogous systems in ANME archaea. **A)** EET systems in Gram-negative bacteria involve membrane-bound quinol:cytochrome *c* oxidoreductases (CbcL, ImcH, CymA, NetD), small soluble cytochromes apparently involved in electron transport through the periplasmic space (PpcA, Stc, PdsA), and a beta-barrel/decaheme cytochrome *c* protein complex (MtrCAB) that acts as an electron conduit by which electrons can transit through the outer membrane to the extracellular space filled with additional cytochrome *c* such as OmcZ and filaments of OmcS. **B)** Analogous protein complexes found in ANME genomes that appear optimized for the challenges associated with EET in the archaeal cell architecture. MpH_2_:cytochrome *c* oxidoreductases are likely present in the form of gene clusters containing VhtC cytochrome *b* subunits together with large 7 or 11 heme-binding MHC proteins (Mco). Other potential options for this step could include the NapH homologs sporadically distributed through ANME genomes, or through the action of the unique cytochrome *b* gene found in ANME Rnf clusters. Electron transfer through the outer proteinaceous S-layer requires a different mechanism than the beta-barrel/decaheme cytochrome strategy evolved in the EET-capable bacteria containing an outer membrane. This step is expected to be overcome by the giant ANME-specific MHC proteins containing S-layer domains allowing them to integrate into the S-layer structure. Very close homologs of OmcZ are found in ANME (see Fig 10). For details of S-layer MHC fusions, see Fig 11.

#### Potential methanophenazine:cytochrome *c* oxidoreductase complexes

In bacteria capable of EET, the quinol:cytochrome *c* oxidoreduction step is carried out by a diverse group of non-homologous enzymes. These can be as simple as CymA in *Shewanella oneidensis* MR-1, which is a four heme-binding cytochrome *c* protein with a single transmembrane helix (107), or more complex, involving multiple subunits such as the recently described NetBCD system in *Aeromonas hydrophila* (108). These complexes are relatively easy to replace, as CymA knockouts in *S. oneidensis* MR-1 can be rescued by suppressor mutants that turn on the expression of completely unrelated quinol oxidoreductase complexes (109). In *Geobacter sulfurreducens* at least two quinol:cytochrome *c* systems coexist, ImcH and CbcL, that appear to be tuned to the redox potential of different terminal electron acceptors (110, 111). The only common features in these systems is the presence of periplasmic cytochrome *c* or FeS binding proteins associated with, or fused to, membrane anchor subunits that facilitate electron transfer from the membrane bound quinol to the periplasmic acceptor (**Fig 9A**).

Due to the non-homologous nature of these electron transport systems in bacteria, we examined the ANME genomes for genes and gene clusters with a potential for analogous function, but adapted to the specifics of archaeal cell biology (**Fig 9B**). ANME-2a, 2b and 2c were found to encode a membrane-bound cytochrome *b* that is closely related to the membrane-bound subunit of the Vht hydrogenase (VhtC), which in methanogens mediates the reduction of methanophenazine with electrons from H_2_ (112) (**S11 Fig**). In the genomes of ANME-2a and 2b, this cytochrome *b* is followed by a multiheme *c*-type cytochrome (MHC) containing between 7 and 11 heme-binding motifs (CxxCH), instead of the *vhtAG* genes encoding the hydrogenase subunits found in *Methanosarcina*. In ANME-2c a closely related homolog for this MHC protein was found elsewhere in the genome. This conspicuous gene clustering is not found in any methanogenic archaea, and the importance of this system is supported by the high expression of the cytochrome *b* subunit reported in both ANME-2a and ANME-2c (9, 11). Phylogenetic analysis shows these VhtC homologs to be a closely related sister group to those found in *Methanosarcina* (**S12 Fig**). These gene clusters are the clearest examples of biological novelty well conserved in ANME that could explain the evolution of electron transfer capabilities, and we refer to them here as methanophenazine-cytochrome c oxidoreductase (Mco).

Notably, ANME-1 and ANME-3 genomes did not contain homologs to Mco, so this does not appear to be a universal mechanism of electron transport in all ANME. ANME-1 lacked any identifiable cytochrome *b*, while the only ones apparent in ANME-3 were multiple copies of *hdrE* and the gene associated with the Rnf gene cluster mentioned above. These Rnf-associated cytochrome *b* in ANME-2a, 2b and 3 are not found in any related methanogens. It is conceivable that these genes are involved in the oxidation of MpH_2_ and in the transfer of these electrons onto the Rnf-associated cytochrome *c* for the purpose of EET. Two recent studies on *M. acetivorans* have implicated the Rnf-associated cytochrome *c* in electron transfer to Fe(III) or the artificial electron acceptor AQDS (113, 114). Such a process may be occurring in the ANME-SRB syntrophy via a similar mechanism and explain previous results of marine ANME utilizing AQDS, Fe(III) and Mn(IV) (115, 116).

Another possible MpH_2_/QH_2_ oxidizing systems could be represented by diverse homologs related to the membrane-bound periplasmic nitrate reductase subunit NapH which are sporadically distributed across ANME genomes. NapH typically contains 4 transmembrane helices, two conserved cytoplasmic FeS clusters, and mediates electron flow from a menaquinol to the periplasmic nitrate reductase NapAB (117). The NapH homologs in ANME are not found in suggestive gene clusters except in one of the ANME-2c genomes in which NapH was followed immediately by an 8 heme MHC, reminiscent of quinol:cytochrome *c* oxidoreductase gene clusters described in bacteria (**Fig 9B**). NapH homologs are found in both ANME-3, as well as two ANME-1, and are worthy of investigating further due to the lack of other obvious candidates for menaquinol/methanophenazine oxidation in those genomes. However, under standard laboratory AOM conditions (9), the NapH homolog found in ANME-2c E20 had very low expression levels (**S3 Data**). Due to their uneven distribution in ANME genomes these NapH homologs may have other non-essential functions.

While no other gene clusters could be identified that seemed likely to mediate MpH_2_/QH_2_:cytochrome *c* oxidoreduction, the example of CymA in *S. oneidensis* MR-1 highlights how simple these systems can be while still being vitally important respiratory proteins, and how easily one system can be functionally complemented by a completely unrelated one (107, 109). Numerous small multiheme cytochromes are encoded and expressed in ANME genomes that either have single membrane anchors on their C-terminus or PGF-CTERM archaeosortase motifs that are predicted to covalently link them to membrane lipids. These small MHC proteins could potentially act in a similar fashion to CymA. However, without an associated or fused large membrane anchor that is homologous to previously characterized systems it is difficult to implicate them directly in membrane bound electron carrier oxidation through genomic evidence alone.

#### Multiheme cytochrome *c* protein abundance and expression

ANME genomes contained many (5-8 times) more genes encoding MHCs than any of their methanogenic relatives (**S13 Fig**). Only the “*Ca.* Syntrophoarchaeum” and some members of the *Archaeoglobales* that are known to conduct EET contain a similar number of MHC genes among the archaea. Many homologous groups of these MHCs are only present in ANME and the aforementioned archaea. All MHC proteins are predicted to reside in the extracellular space as heme attachment to the cytochrome *c* apoprotein occurs here (118). Representatives of these MHCs are among the highest expressed proteins in multiple ANME lineages. Small MHC in ANME-2c E20, ANME-1 GB37 and ANME-1 GB60 were the 14^th^, 10^th^, and 18^th^ highest expressed proteins, respectively, and contain 8, 4 and 4 heme binding motifs, respectively (**S3 Data**). One specific group of small MHC proteins has widespread distribution in the ANME, “*Ca.* Syntrophoarchaeum”, and some members of the *Archaeoglobales*, with no relatives in methanogenic archaea. Multiple paralogs exist in most ANME groups and these include some of the highest expressed proteins in ANME, many exceeding all methanogenesis pathway genes except Mcr (**S14 Fig, S3 Data**).

As with the quinol oxidation step, the intermediate small soluble cytochromes *c* vary greatly between different EET-capable bacteria. Any number of these ANME cytochromes could be capable of carrying electrons between the cytoplasmic membrane and outermost layer of the cells. A small six heme-binding cytochrome *c* protein OmcS was recently shown to form the conductive extracellular “nanowires” in *G. sulfurreducens* that are thought to imbue the biofilms with conductive properties (**Fig 9**) (106, 119). No close homolog of OmcS were identified in ANME genomes, although any number of these relatively small cytochromes could be carrying out a similar function in the extracellular space between ANME and SRB, consistent with ultrastructural observations made with heme-reactive staining (9, 87).

An 8-heme MHC in *G. sulfurreducens* known as OmcZ was shown to be required for optimal anodic current in biofilms grown on electrodes, and is secreted in biofilms and in culture supernatant (120). It was recently shown that OmcZ will also polymerize, forming conductive nanowires (121). Very closely related homologs of OmcZ were found in multiple ANME-2a, 2b, and 2c genomes (**Fig 10**). The sequence similarity shared between OmcZ and the proteins found in these ANME groups is not limited to CxxCH motifs, but also extensive N- and C-terminal regions with many completely conserved residues. This level of sequence conservation is quite remarkable for homologs found in archaea and bacteria, and suggest an important conserved function of these non-heme binding domains, as well as a relatively recent inter-domain horizontal transfer. The OmcZ homolog in ANME-2c E20 is the 14^th^ highest expressed protein in cultures grown under standard laboratory AOM conditions (**S3 Data**). In ANME-2a and 2b the genes encoding these OmcZ homologs are found next to the enormous MHCs described below. Investigations into the properties of ANME OmcZ homologs, specifically whether they can undergo the same polymerization observed in *G. sulfurreducens*, will be an important future area of research.

**Fig 10.**
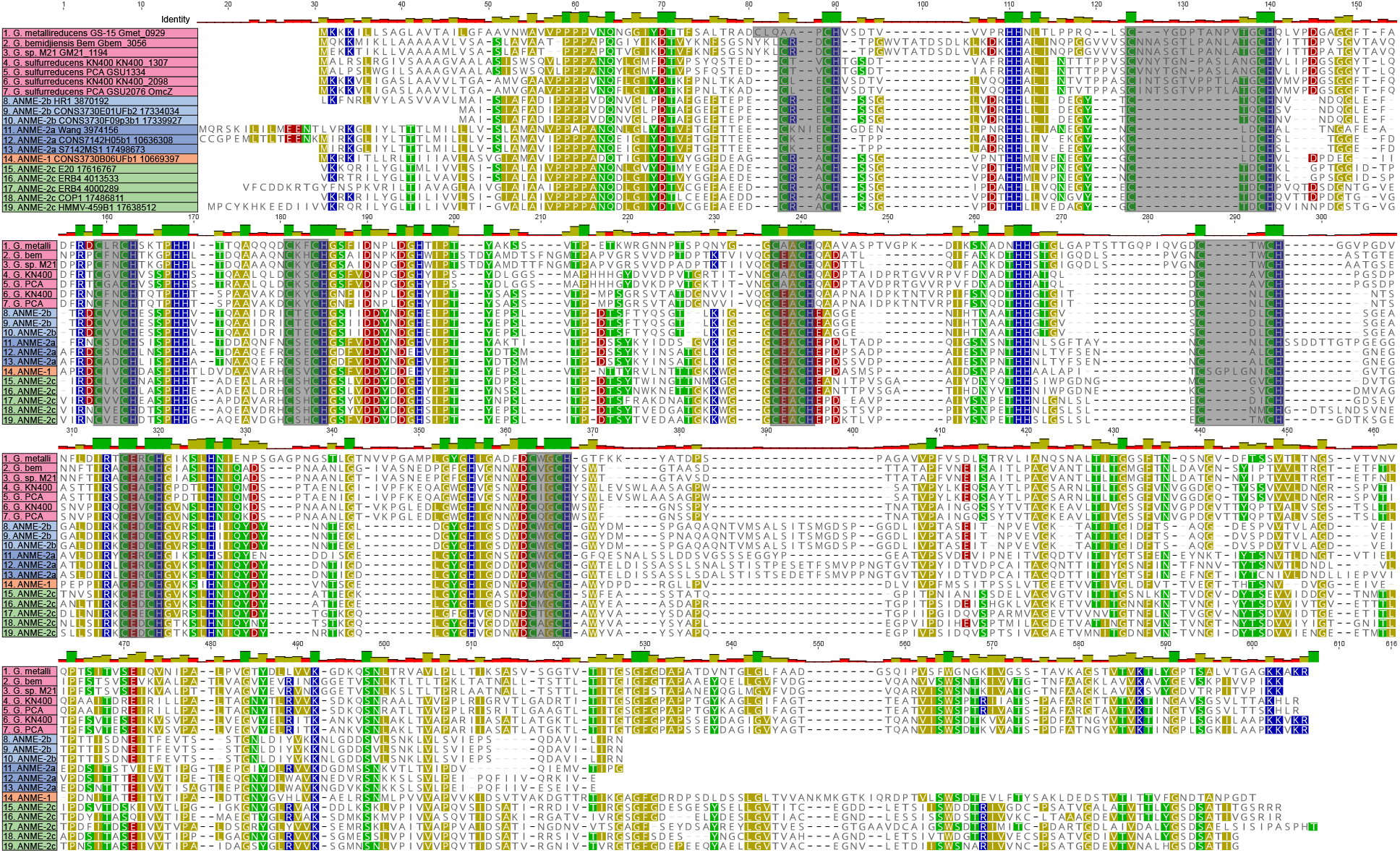
OmcZ homologs in ANME archaea. Protein sequence alignment of OmcZ homologs from various *Geobacter* species and ANME genomes reported here using muscle 3.8.31 with default settings. The eight CxxCH binding motifs are highlighted in gray. Regions of significant sequence identity are present throughout the protein, not just associated with the CxxCH motifs suggesting conserved function. Alignment file can be found in **S1 Data**.

#### S-layer conduits

ANME-2a, 2b, 2c, 2d, and 3 genomes also contain exceptionally large MHCs with 20 and 80 heme binding sites (**Fig 11, S13 Fig**). Similarly large cytochromes in archaea are only found in *Geoglobus* and *Ferroglobus* (87). Both of these groups are capable of extracellular iron reduction (122, 123). The largest and best conserved of these MHCs were classified into three major groups (ANME MHC Type A, B, and C) based on sequence identity, conserved domains, and heme-binding motif distribution (**Fig 11**). The most notable feature of ANME MHC-A and B is the presence of predicted S-layer domains, which commonly make up the outer proteinaceous shell surrounding the cytoplasmic membrane in many archaea (124). ANME MHC-B additionally contain an N-terminal domain free of heme-binding motifs and annotated as “peptidase M6-like domains”. The presence of an S-layer domain in these cytochromes suggests their use for electron transfer through this outermost layer. In *M. acetivorans* the S-layer domain structure has been determined by x-ray crystallography and contains two subdomains connected by a flexible linker (125). In all ANME MHC with S-layer domains, this flexible linker also contains a single heme-binding motif, which would place a heme group within the plane of the S-layer. Both ANME MHC-A and B encode for C-terminal regions predicted to be transmembrane helices that may anchor them cytoplasmic membrane. We suggest that ANME MHC-A and B are functionally analogous to the MtrCAB complexes found in the outer membranes of the EET-capable Gram-negative bacteria (**Fig 9A**) (104). In ANME an alternative mechanism needs to be employed since the outermost layer is proteinaceous.

**Fig 11.**
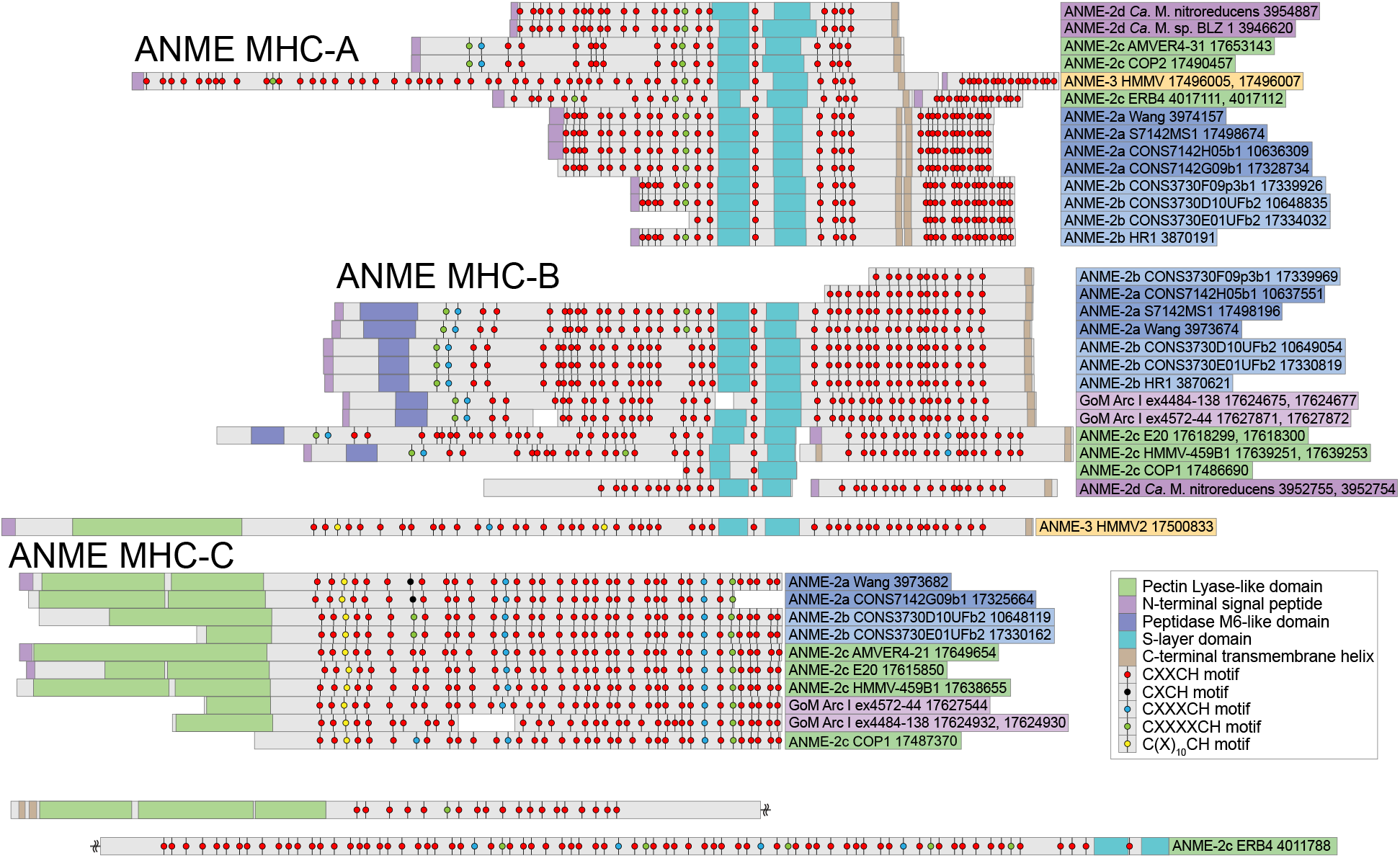
Large multiheme cytochrome c proteins in ANME archaea. Schematic of protein structure highlighting the position of heme-binding motifs and other conserved features of the large ANME-specific multiheme cytochromes. The ANME MHC were divided into three major groups based on sequence similarity and conserved domains structure. ANME MHC-A contains an S-layer domain and C-terminal transmembrane helix. In ANME-2a and 2b these proteins are extended by an additional transmembrane helix and more heme-binding motifs. ANME MHC-B contains an S-layer domain and C-terminal transmembrane helix as well, but an N-terminal region devoid of heme-binding domains has similarity to peptidase M6-like domains. ANME MHC-C do not contain S-layers or C-terminal transmembrane helices, but instead contain a large N-terminal region with a predicted pectin lyase-type domain. Domains predicted with InterProScan and are displayed with colored boxes. Large MHC proteins from ANME-3 sp. HMMV2 and ANME-2c sp. ERB4 that do not clearly fit into these categories are also shown (Note: ANME-2c sp. ERB4 is a single peptide split between two lines due to its size).

ANME MHC type C do not encode for an S-layer domain or C-terminal transmembrane helices, but do encode for a large N-terminal “pectin lyase-like domain”. It is difficult to predict the function of this additional domain. However it is interesting to note that the pectin lyase fold typically occurs in proteins that attach to and/or degrade carbohydrates, and has been found in bacteriophage tail spike proteins for attachment to hosts (126). It is possible then that this domain is involved in recognizing the outer cell wall of partner bacteria. Some ANME genomes contained exceptionally large MHC that did not fall clearly into these three categories. ANME-3 HMMV2 for example contained a single polypeptide containing C-terminal features of ANME MHC-C and N-terminal features of ANME MHC-B (**Fig 11**). A fosmid assigned to ANME-2c ERB4 encoded the largest MHC in our dataset containing an S-layer domain, a pectin lyase-like domain, and 86 heme binding motifs.

In actively growing ANME-2c cultures, genes encoding these large ANME MHC were expressed at lower levels than the enzymes of the reverse methanogenesis pathway and some of the smaller MHCs leading the authors to conclude that these were of little importance to electron transfer in this culture (9). If two proteins are expected to carry out the same function in a pathway, transcription levels may be useful in determining which may be the dominant one operating under certain conditions. But this is not the case for the different classes of MHCs in the model of ANME EET presented here. The small MHCs are thought to serve as intermediates between the inner membrane and S-layer, as well as assist in conveying conductivity to the extracellular matrix, while the large S-layer proteins specifically provide a conduit through the S-layer. The small MHCs may cover longer distances, and would therefore be expected to be numerically dominant, but still could not make a functional EET system if no specific mechanisms existed for the short transfer through the S-layer to the extracellular environment. This is exactly the scenario observed in the electrogenic model bacterium *G. sulfurreducens* that requires MHC conduits through the outer membrane for EET formed by MtrCAB and other analogous porin-cytochrome systems (104). While absolutely necessary for efficient EET, these genes show lower expression levels than those of some of the smaller soluble MHCs encoded in the genome (127). We therefore find this previously reported transcript information to be consistent with the model of ANME-2 and 3 EET presented here in which the S-layer fusions play a key role (**Fig 9B**).

ANME-1 contained fewer MHCs and at most had 10 predicted heme-binding motifs, and these proteins did not contain any identifiable S-layer domains. From the available genomic data, it is unclear how these cytochromes fit into the cell structure of ANME-1. It was recently suggested that ANME-1 lack an S-layer due to the absence of predicted S-layer domain containing proteins (42). However, the same analysis found no *Archaeoglobus* proteins with S-layer domains, yet the *Archaeoglobus* S-layer has been visualized and extensively characterized (128). The S-layer domain in the large MHC from ANME-2 and 3 are only recognizable because they are homologous to the S-layer protein of *M. acetivorans* that was recently crystalized (125). Before this study the S-layer domain was only identified as a domain of unknown function (DUF1608). The ANME-1 ultrastructure is usually cylindrical and highly reminiscent of *Methanospirillum* which contain S-layers (129). How the MHC in ANME-1 fit into this cell structure will require further investigation. Gram-positive bacteria are also capable of EET, and in some cases have been found to use more modestly sized MHC proteins (6-9 heme binding motifs) for electron transport through the outermost cell wall (130). Such a model could very well apply to the cell wall and cytochromes encoded in the ANME-1 genomes.

#### Duplication of Cytochrome *c* maturation machinery

Cytochrome *c* proteins are matured on the outer face of the cytoplasmic membrane through a variety of systems, the most common in bacteria and archaea being the cytochrome *c* maturation (Ccm) system (118). Ccm transports heme B to the positive side of energized membranes and mediates its attachment to the cytochrome *c* apoprotein (**Fig 12**). The Ccm system has been found in archaea closely related to ANME (131), and slight modifications to these systems in archaea have been investigated in detail (132). Most ANME genomes contained CcmABCEF and lacked CcmH a pattern observed previously in most other archaea including methanogens. One exception to this is the freshwater ANME-2d, which contains CcmH homologs similar to those found in *Ferroglobus* and *Geoglobus sp.* (131, 132).

**Fig 12.**
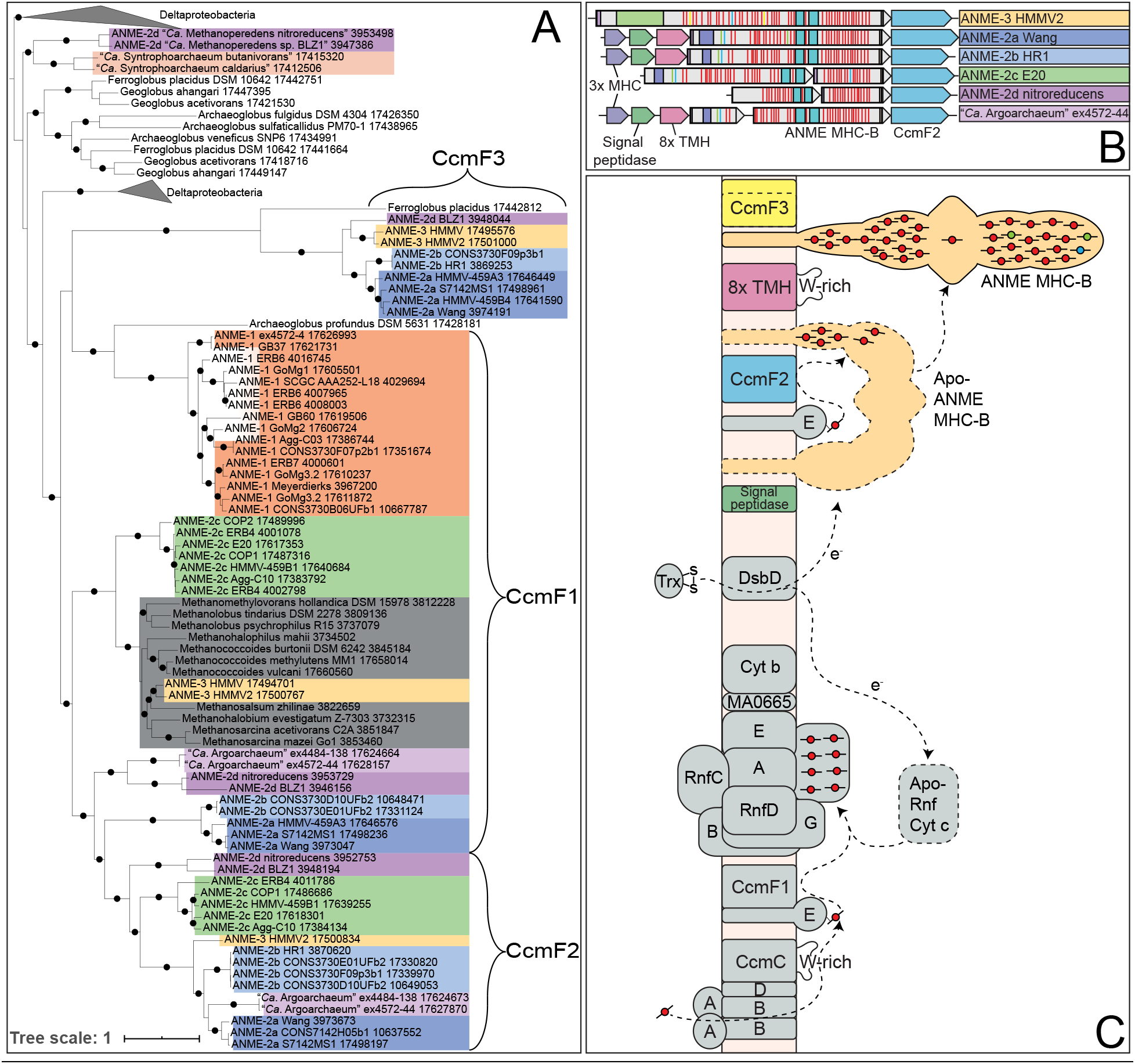
Cytochrome maturation and CcmF duplication. (**A**) Phylogenetic analysis of CcmF homologs from ANME and closely related archaea. CcmF1 cluster contains the CcmF found in the *Methanosarcinaceae*. CcmF2 is a closely related group of homologs found only in ANME and “*Ca.* Argoarchaeum”, and in all cases is found next to the S-layer containing ANME MHC-B. CcmF3 comprises the larger of two encoded genes that appear to be a split CcmF, and are found in ANME and *Ferroglobus*. (**B**) Example gene clusters from all groups that contain CcmF2 illustrating their position with respect to ANME MHC-B. Some genomes additionally contain a signal peptidase and a predicted membrane integral ANME-specific gene in this cluster (8x TMH). Horizontal red lines denote CxxCH heme binding domains, teal represent S-layer domain (see Figure 11 for details of ANME-MHC structure). (**C**) Schematic of cytochrome maturation pathway. CcmA and B comprise ABC transporter module that exports heme B, which is transferred to CcmE via CcmC’s tryptophan (W)-rich periplasmic loop. CcmE is expected to utilize CcmF1 to mature cytochrome *c* proteins found in both ANME and methanogens of the *Methanosarcinaceae*. CcmF2 found only next to ANME MHC-B is expected to be involved in its maturation. The co-occurring signal peptidase is likely involved in cleavage of the N-terminal signal sequence. Closed circles represent branch support values between 80 and 100%. Tree scales represent substitutions per site. Tree construction parameters are found in the Materials and Methods section. Alignment and tree files can be found in **S1 Data**.

A very interesting feature of the ANME genomes reported here is the presence of multiple paralogs of CcmF. CcmF is the largest of the Ccm proteins, and is involved in the transfer and attachment of heme B to the cytochrome *c* apoprotein. One of these highly divergent CcmF paralogs that is conserved in ANME-2a, 2b, 2c, 2d, 3 and “*Ca.* Argoarchaeum” is found in every case next to the ANME MHC-B (**Figs 11** and **12B**). The diversification of CcmF paralogs in ANME and the clustering of one of the CcmF paralogs with ANME MHC-B suggests that special CcmF proteins may be needed to handle the extremely large apoproteins associated with these MHCs. The specialization of CcmF homologs to particular cytochromes *c* is not without precedent, a specific CcmF homolog NrfE is known to be required for insertion of heme B into NrfA at a modified heme binding site with a CxxCK motif (133). These additional CcmF homologs in ANME may be required due to the extreme size of the ANME MHC apoproteins or due to the presence of multiple modified heme biding motifs with increased amino acids between the cysteine residues (**Fig 11**).

Immediately upstream of the CcmF/ANME MHC-B genes in ANME-2a, 2b and 3 are two highly conserved genes also expected to be involved in MHC maturation (**Fig 12B, S11 Fig**). One is a signal peptidase which is likely involved in the cleavage of the N-terminal signal sequence present on most of the encoded MHC proteins in ANME. The second is an integral membrane protein that has no automated annotations (labelled 8x TMH). This later gene is found in every ANME group, “*Ca.* Syntrophoarchaeum”, “*Ca.* Argoarchaeum”, *Geoglobus*, and *Ferroglobus*. BLAST searches against the NCBI nr database revealed no homologs to this protein in any other organisms. While no specific motifs are clearly obvious, a large periplasmic loop is predicted that contains multiple conserved tryptophan residues which is a common feature of the heme handling proteins (HHP) family to which both CcmF and CcmE belong (133). This protein may have a role in heme handling in the maturation process of the MHC proteins in ANME as well. These two genes are also found in a cluster of cytochrome *c* genes highlighted in a previous metagenomic study of ANME-1 (7). Due to their similarity to peptidases and heme handling proteins we suspect these are not membrane anchors for the mature MHC proteins as previously posited. The unusual ANME-1 subclade consisting of GoMg4 and SA contained no cytochrome *c* genes and also lacked CcmABCE, all CcmF paralogs, the signal peptidase, and the integral membrane protein described above. In conclusion, a key genomic difference between canonical methanogens and the ANME archaea are traits linked to direct interspecies electron transfer, especially large, diverse cytochromes and associated biosynthetic machinery. It should be noted that it is possible for engineered syntrophies to occur with methanogenic *Methanosarcinaceae* even in the absence of MHC, although the mechanistic details of how electrons are transferred in this system are lacking (134).

### Anabolic pathways

In addition to energy metabolism reconstruction, the diversity of ANME genomes presented here allows for an evaluation of the anabolic pathways present in the different methanotrophic lineages. The assimilation of isotopically light (^13^C-depleted) carbon into ANME lipids and bulk biomass was crucial to their initial discovery (1–3, 83, 135), and is an important signal used for the interpretation of stable isotope studies of ancient carbonate systems. A better understanding of the precise biochemical pathways available for building this biomass will help better interpret these types of analyses, as well as the results of stable isotope probing experiments. Early experiments also highlighted how isotope signatures serve to identify the syntrophic cooperation between the consortia partners (3, 5). In this regard, long-term symbiotic interactions between organisms has been hypothesized to lead to lasting anabolic impressions on the organisms involved, specifically the adaptive loss of anabolic pathways such that syntrophic partners are left with complementary set of functions such as amino acid biosynthesis (136, 137). Below we highlight some of the key conserved features and differences between anabolic pathways present across the different groups of ANME archaea.

#### Anabolic C1 metabolism

C1 metabolism concerns the enzymes, reaction and cofactors mediating redox transformations of single carbon molecules and the ways these are incorporated into various biomolecules through the formation of carbon-carbon bonds. The single known anabolic C1 pathway present in all ANME is the Wood-Ljungdahl (reductive acetyl-Coenzyme A) pathway. The methyl branch of this pathway is the methanogenesis pathway described in previous sections. In this pathway, methyl groups attached to H_4_MPT are combined with CO_2_ and a pair of electrons from Fd^2-^ to make acetyl-CoA which can then be used in a wide variety of biosynthetic processes. This pathway ANME share with methanogens, some of which are autotrophic, i.e. derive their cell carbon from CO_2_ reduction. Still, an important question for interpreting isotopic studies is whether there are other means of C1 assimilation and, if so, how these alternative mechanisms vary within and between ANME clades.

The H_4_MPT cofactor of the Wood-Ljungdahl pathway was first discovered in methanogenic archaea and was originally thought to be an archaea-specific modified folate substituting in place of H_4_F in all methanogens and their relatives for C1 carrying reactions. This view held until it was shown the H_4_MPT was also present in methylotrophic bacteria (138). These bacteria contain both H_4_MPT and H_4_F and assessing roles of the C1 pools associated with each pathway for either catabolism or anabolism was a major challenge requiring decades of biochemical, genetic and comparative genomic analysis to begin to fully understand (139). In methylotrophic bacteria H_4_MPT appears to be the dominant catabolic oxidation pathway for formaldehyde generated by methanol dehydrogenase, while H_4_F can either serve as a catabolic oxidation pathway for certain methylated compounds or as a mechanism for transferring C1 moieties from formate into the anabolic serine cycle. Because of the nuanced understanding of C1 metabolism in methylotrophic bacteria and the apparent importance of maintaining separate C1 pathways in these organisms, we investigated the extent to which H_4_F-bound C1 pools may be present in the ANME archaea (**Fig 13**).

**Fig 13.**
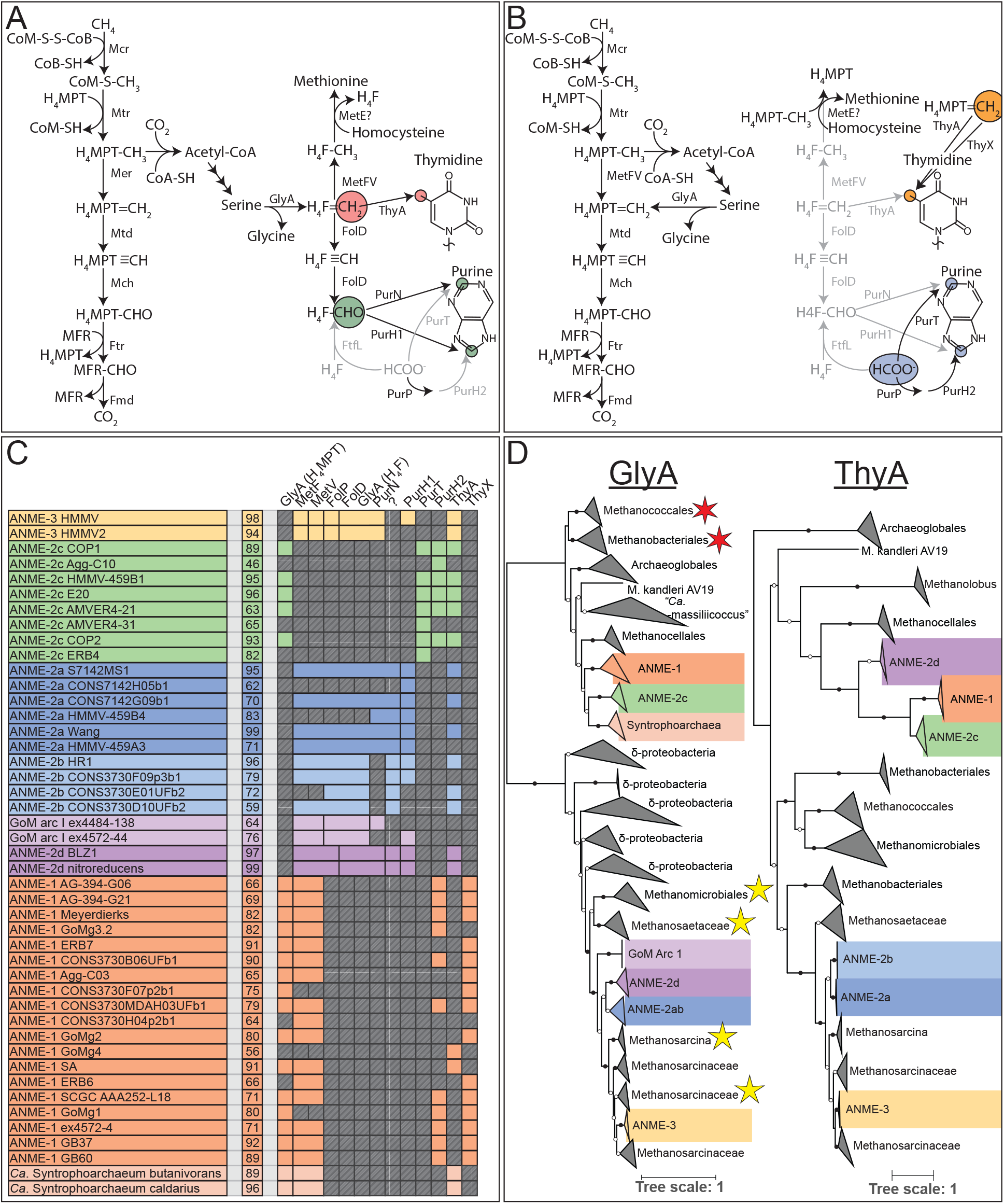
H_4_MPT vs H_4_F C1 metabolisms. (**A**) Pathway of C1 transformations in ANME-2a, 2b, 2d and 3 based on presence of genes and precedent from *M. barkerii*. H_4_MPT is the catabolic C1 carrier, while the H_4_F C1 pool is derived from serine and is used for the biosynthesis of purine, thymidine and possibly methionine. (**B**) Pathway of C1 transformations available to ANME-1 and 2c which lack the vast majority of the H_4_F interacting enzymes in (**A**). H_4_F-CHO-specific enzymes in pyrimidine synthesis are replaced by free formate versions in ANME-2c. While present in these organisms, GlyA appears to be the type that interacts with H_4_MPT instead of H_4_F. Light gray pathways were not found. (**C**) Colored boxes represent presence of various H_4_F-interacting genes in ANME genomes. Missing genes are represented by gray boxes with diagonal line fill. Numbers in the second column represent genome completeness. When genes are together in a gene cluster their boxes are displayed fused together. Gene accession numbers can be found in **S2 Data.** (**D**) Phylogenetic analysis of GlyA and ThyA homologs found in ANME genomes. Red and yellow stars indicate GlyA sequences shown to react with H_4_MPT and H_4_F, respectively. Closed circles represent branch support values of 100%, open circles >50%. Tree scales represent substitutions per site. Tree construction parameters are found in the Materials and Methods section. Alignment and tree files can be found in **S1 Data**.

In ANME-2a, 2b, 2d and 3, MetF is clustered with the genes for MetV, FolP, FolD, GlyA and PurN, all of which are involved in H_4_F-related anabolic C1 reactions (**Fig 13C**). This cluster of genes is also found in other closely related methanogens in the *Methanosarcinaceae* (51). PurH1, encoded elsewhere in these genomes, also utilizes H_4_F. A range of biochemical and isotope labelling studies found that these enzymes are involved in anabolic pathways and have demonstrated their specificity for H_4_F in *M. barkeri*. Importantly, the C1 moieties in the H_4_F pathway are not derived from free C1 compounds, as the critical enzyme formate H_4_F ligase is not present in any ANME genome. Instead, in ANME the carbon atoms in compounds that are derived from methylene-H_4_F will come from the C2 position of acetyl-CoA as was shown in *M. barkeri* (51). Acetyl-CoA derived carbon passes through pyruvate to serine, and through the activity of serine hydroxymethyl transferase (GlyA), the carbon derived from the C2 of acetate is transferred from serine to H_4_F to be used for biosynthesis of purines, thymine and possibly methionine (**Fig 13A**). The methyl group donor for methionine biosynthesis is likely a corrinoid protein, but its identity remains unknown in methanogens (140).

ANME-2c lacks a MetF of either the normal *Methanosarcinaceae* variety, or the ANME-1/uncultured archaeal version. Consistent with the lack of the *Methanosarcinaceae* variety of MetF, all ANME-2c genomes additionally lack MetV, FolP, FolD, PurN and PurH1 (**Fig 13C**). This absence of H_4_F-interacting genes is also true of all the ANME-1 genomes, save for MetV which is found alongside the ANME-1 MetF. This suggests H_4_F is not used in ANME-1 or 2c for biosynthetic reactions (**Fig 13B**)

GlyA homologs are found in all ANME genomes. Since GlyA is expected to interact with H_4_F in ANME-2a, 2b, 2d and 3, we conducted a deeper analysis of GlyA in order to find an explanation for their presence in ANME-1 and 2c since they seemed to lack the other H_4_F-related genes. Phylogenetic analysis of GlyA showed a clear differentiation between GlyA in ANME-1 and 2c vs. those in ANME-2a, 2b, 2d and 3 (**Fig 13D**). GlyA in methanogens without H_4_F react with H_4_MPT (54, 55), while the *Methanosarcinaceae* with H_4_F contain GlyA active towards H_4_F (51). ANME-1 and 2c GlyA sequences cluster with the GlyA that have been biochemically characterized to interact with H_4_MPT, while GlyA from other ANME cluster with those of *Methanosarcinaceae* and would be expected to react with H_4_F (141). This pattern of presence/absence of genes and phylogenetic affiliation of homologs suggests that H_4_F is present in the ANME-2a 2b 2d and 3, H_4_F is present and used in the same manner as *M. barkeri*. In ANME-1 and 2c we expect that GlyA is necessary to produce glycine from serine, but that the resulting C1 moiety is shuttled back into the H_4_MPT-bound C1 pool.

PurH1 and PurN which are absent in ANME-1 and ANME-2c utilize formyl-H_4_F as a source of C1 moieties for two of the carbon atoms in the biosynthesis of purines (**Fig 13A**). An interesting question is how ANME-1 and 2c are able to grow without these enzymes. Alternate enzymes, PurT and PurP/PurH2, can carry out these steps, but use free formate as the C1 donor (142). Notably, ANME-2c contained both PurT and PurH2 which are absent in the ANME-2a, 2b, 2d and 3, while ANME-1 genomes contained PurH2 but lacked PurT. A lack of both PurT and PurN has been observed in *Archaeoglobales* and *Methanobacteriales*, which suggests that there is an as yet unidentified enzyme catalyzing this step since both are known to synthesize their own purines (142). This distribution of genes involved with purine biosynthesis indicates at least an anabolic role for formate in ANME-1 and 2c metabolism.

Another important anabolic C1 reaction is carried out by thymidylate synthase, a crucial step in the synthesis of the thymidine base of DNA, which catalyzes the methylation of deoxyuridine monophosphate (dUMP) to produce deoxythymidine monophosphate (dTMP). This C1 moiety is often derived from 5,10-methylene-H_4_F, and this reaction can be carried out by two non-homologous thymidylate synthase proteins, known as ThyA and ThyX (143). The reactions catalyzed by these enzymes are slightly different, with ThyA using the H_4_F itself as an electron donor to reduce the methylene to methyl, producing a dihydrofolate product. ThyX in contrast utilizes NAPDH in the reaction, leaving H_4_F in the tetrahydro oxidation state.

The genomes of ANME-2a, 2b, 2c, 2d, and 3 contain ThyA, whereas most ANME-1 have ThyX. The only exception to this is the small ANME-1 subgroup comprised of genomes GoMg4 and SA that contain ThyA instead of ThyX, adding to the list of unique features of this ANME-1 subclade. Phylogenetic reconstruction of ThyA homologs revealed a similar split between ANME with and without H_4_F; ANME-2a, 2b, and 3 cluster with their close methanogenic relatives, while ANME-1 and 2c cluster with *Methanocellales*. Curiously ANME-2d also cluster with ANME-1 and 2c although they contain all the other H_4_F interacting proteins. Much less is known about ThyA biochemistry in methanogens. Labeling and biochemical studies have not been carried out to the same degree as with GlyA making it is difficult to predict the cofactor specificity for either of these ThyA clusters. Due to the presumed lack of H_4_F in ANME-1 and 2c the ANME-1, 2c and 2d cluster likely utilizes H_4_MPT, while the ANME-2a, 2b and 3 cluster utilizes H_4_F (**Fig 13A** and **13B**).

The difference between using ThyX or ThyA could be particularly useful because thymidylate synthase has recently been the subject of great interest as a druggable target in *Mycobacterium tuberculosis*. Humans use ThyA, while ThyX is thought to be essential for *M. tuberculosis*, and various ThyX inhibitors are being investigated for clinical antimicrobial uses (144). If ThyX inhibitors are shown to specifically inhibit ANME-1 they could be valuable tools for determining the contribution of ANME-1 to methane oxidation in mixed ANME communities.

#### Apparent amino acid prototrophy

Obligate coupling between the energy metabolisms of syntrophic organisms has been suggested to lead to additional metabolic dependencies such as amino acid or vitamin auxotrophies through adaptive gene loss (136, 137). Yet, the genomes of all ANME clades contain a near complete complement of genes predicted for synthesizing all amino acids, including many of the pathway modifications described from cultured methanogens (**S15 Fig**). Of the enzymes that are widespread in related methanogens, only phosphoribosyl-ATP diphosphatase (EC 3.6.1.31), a step of histidine biosynthesis is not found in any other ANME genomes, with the exception of ANME-3 (**S15 Fig**).

The gene responsible for this step is also absent in *Archaeoglobus fulgidus* and *Nitrosopumilus maritimus*, both of which are capable *de novo* histidine synthesis, suggesting other unknown mechanisms exist for completing this step of histidine biosynthesis (145).

Other isolated steps in amino acid synthesis were also unannotated in ANME genomes, however they were also missing in related methanogens known to complete these steps in an as yet uncharacterized manner. ANME genomes appear to lack phosphoserine transaminase but this is possibly complemented by a broad specificity aspartate aminotransferase as has been described in other methanogens (146). Aromatic amino acids appear to be produced via the 6-deoxy-5-ketofructose-1-phosphate (DKFP) pathway to produce shikimate (147–149), however DKFP synthase was not detected in most of the ANME genomes or in many of the *Methanosarcinales*, again suggesting an as yet unidentified alternate gene may be used in this process (**S15 Fig**). Methionine appears to be synthesized using the partially described pathway converting aspartate-semialdehyde directly to homocysteine, though not all enzymes are known for this pathway (150). The final steps of tyrosine and phenylalanine synthesis did not have annotated genes in many methanogens or ANME genomes but may be complemented by a broad specificity amino acid transaminase. Finally, histidinol-phosphatase (EC3.1.3.15) that is also part of histidine biosynthesis is unannotated in all of the ANME and methanogen genomes, suggesting an alternate, undiscovered gene operating at this step.

#### Incomplete partial TCA cycles for 2-oxoglutarate synthesis

Many biomolecules including porphyrins and amino acids are synthesized from 2-oxoglutarate, and this intermediate comes from the TCA cycle beginning with acetyl-CoA sourced from CH_3_-H_4_MPT and CO_2_ via the acetyl-CoA decarbonylase/synthase complex (ACDS). In methanogens, a complete TCA cycle is not needed. Instead, one of two partial pathways is present to produce 2-oxoglutarate for biosynthesis: either the partial TCA cycle operates in the oxidative direction passing through isocitrate, as found in the *Methanosarcinales,* or it runs in the reductive direction through succinate, as in *Methanococcus* (151)(**Fig 14**). Surprisingly, ANME-2a and 2b largely contain the enzymes of the reductive partial TCA cycle, diverging from their close *Methanosarcinaceae* relatives (**Fig 14**). In contrast, the ANME-1 and 2c universally lack some of the genes in the reductive pathway, but contained the three genes catalyzing the reactions in the partial oxidative pathway. This pathway would involve ATP citrate lyase operating in the direction of ATP production, which is not typical. While some ANME-2a come close to having a complete TCA cycle, no single genome was found that contains all steps in the pathway, supporting the notion that these enzymes are more used as a means to make an important biosynthetic intermediate, not for running a complete reverse TCA cycle for carbon fixation. The ANME-3 genomes lack the complete set of genes for either pathway, and it is currently unclear how they can produce 2-oxoglutarate.

**Fig 14.**
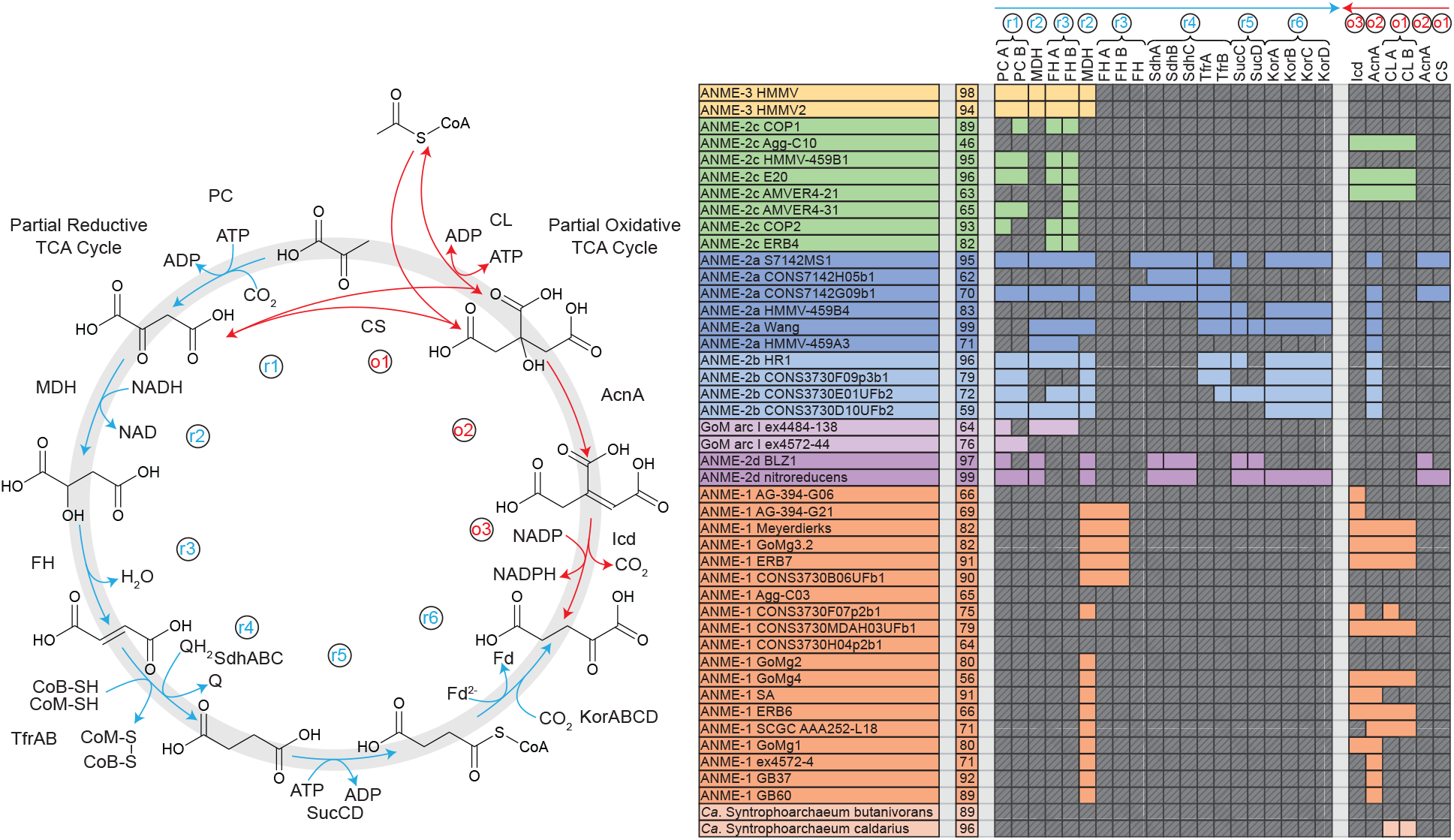
TCA cycle gene present in ANME archaea. (**A**) Schematic of the enzymes and reactants in the partial oxidative and reductive TCA cycles in ANME that lead to 2-oxoglutarate for the purpose of producing anabolic intermediates for biosynthesis. Reductive pathway shown in blue, oxidative pathway shown in red. (**B**) Colored boxes represent presence of various TCA cycle genes in ANME genomes. Missing genes are represented by gray boxes with diagonal line fill. Numbers in the second column represent genome completeness. When genes are together in a gene cluster their boxes are displayed fused together. Note: some steps can be carried out by multiple different enzyme systems. Gene accession numbers can be found in **S2 Data**.

### Additional ANME genomic features of interest

In addition to the catabolic and anabolic pathways described above, we identified a number of other genomic features that appear interesting based either on their unique presence in ANME as compared to closely related methanogens, or their potential for being important in other aspects of the ANME-SRB syntrophy.

#### Nitrogenase in ANME

A phylogeny of the large subunit of nitrogenase, NifD, indicates that the ANME nitrogenases of the “methane seep group” form a monophyletic clade (**Fig 15A**). The closest relatives of this group are from *Roseiflexus castenholzii*, which was also reported for *nifH* gene clones (152). This cluster is notably divergent from the NifD in methanogenic *Methanosarcinales* capable of N_2_ fixation (**Fig 15A**). The ANME NifD, as well as *R. castenholzii* and *Endomicrobium proavitum*, all contain the conserved cysteine and histidine residues for binding the P-cluster and FeMoCo ligands (153), which suggests they may be functional in nitrogen fixation, consistent with *nifH* transcript expression in the environment and correlated FISH-NanoSIMS analysis of ^15^N_2_ assimilation (152). All gene clusters of the methane seep group exhibit the gene order NifHI_1_I_2_DK, and lack the *nifN* and *nifE* genes that function as molecular scaffolds for the maturation of the FeMo cofactor (**Fig 15B)**. These genes were traditionally thought to be crucial for a functioning nitrogenase (154, 155). However, recent reports of nitrogen fixation in *E. proavitum*, which also lack *nifEN*, suggests that these genes are not strictly necessary for nitrogen fixation (156).

**Fig 15.**
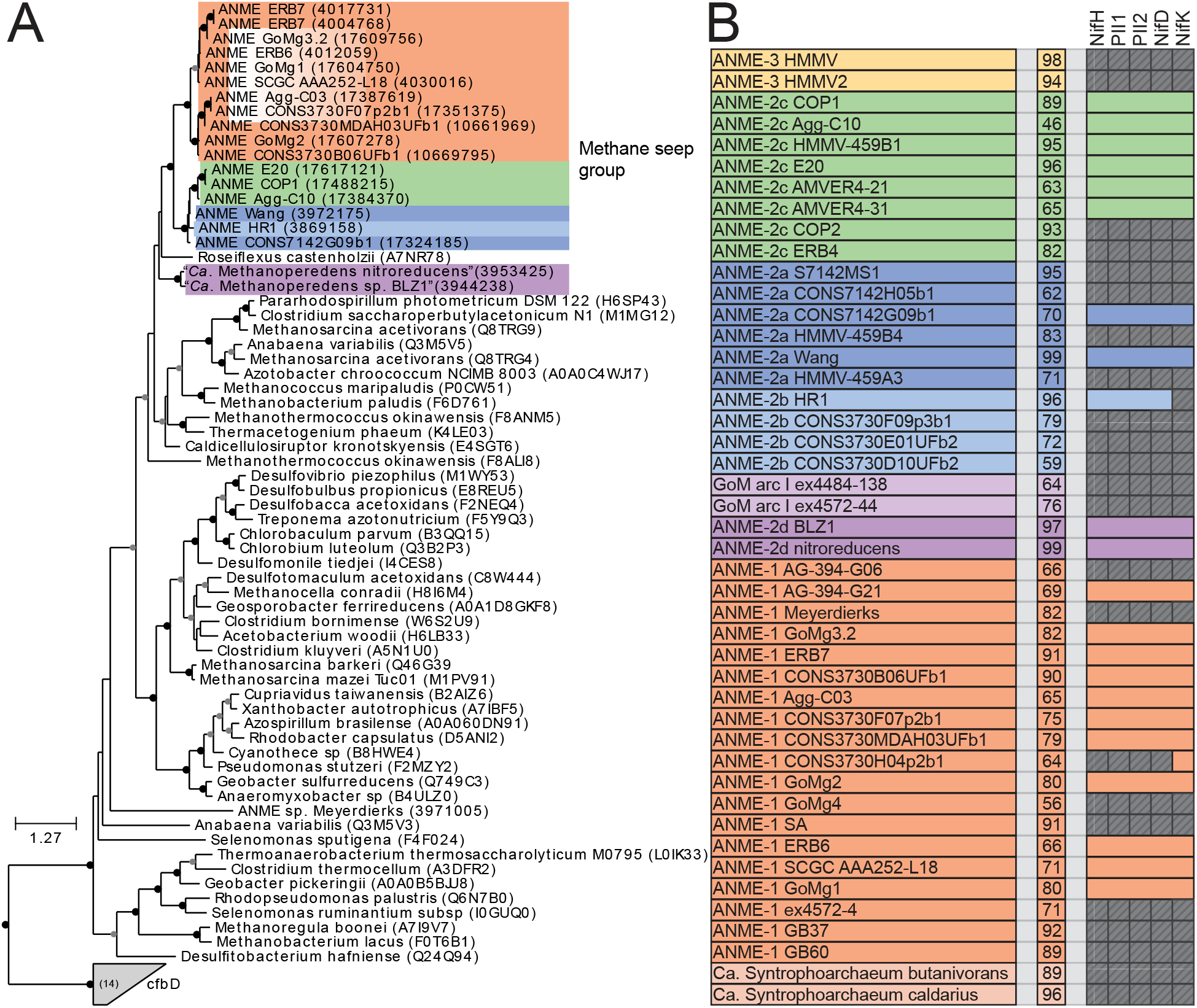
Methane seep group nitrogenase phylogeny and distribution. (**A**) Maximum likelihood phylogenetic tree of NifD amino acid sequences from the “methane seep” group of nitrogenase, with close relatives. Closed circles represent branch support values of 80 to 100%, gray circles between 70 and 80%. Tree scales represent substitutions per site. Tree construction parameters are found in the Materials and Methods section. Alignments and tree files can be found in **S1 Data**. (**B**) Presence of seep group nitrogenase in genomes presented here. Colored boxes represent presence of various nitrogenase related genes in ANME genomes. Missing genes are represented by gray boxes with diagonal line fill. Numbers in the second column represent genome completeness. When genes are together in a gene cluster their boxes are displayed fused together. Gene accession numbers can be found in **S2 Data**.

It was unexpected to find nitrogenases of the “methane seep group” in many of our ANME-1 genomes. Clone libraries of *nifH* amplicons from methane seeps were used to define four major groups present and expressed in these habitats referred to as: methane seep group, methanosarcina-like group, group III and group IV (157). The methane seep group is composed of sequences almost exclusively found in seep environments and was assigned to members of the ANME-2 based on multiple lines of evidence (34, 158). Group IV were detected in ANME-1 fosmids and shown to be poorly expressed by metatranscriptome analysis. This low expression coupled with the fact that group IV NifH were proposed to be involved in F_430_ biosynthesis instead of nitrogen fixation (159) led to conclusion that ANME-1 are likely incapable of nitrogen fixation (7). Group III NifH sequences are largely comprised of sequences similar to those found in *Deltaproteobacteria*, likely corresponding to the ANME’s syntrophic partners (157). The *Methanosarcina*-like group is very similar to sequences found in genomes of the *Methanosarcina* genus, and they were expected to come from low abundance methanogens occasionally present in methane seep environments. The wide distribution of methane seep group nitrogenase in addition to nitrogenases belonging to the group IV in ANME-1 suggest that these organisms may also be capable of nitrogen fixation.

The monophyletic nature of the nitrogenase genes in ANME is notable in the context of the paraphyletic nature of ANME organisms. This form of nitrogenase may have been present in the common ancestor of all ANME, and was subsequently lost in all methanogenic members of the *Methanosarcinaceae* and then replaced with a phylogenetically distinct nitrogenase in some lineages. Alternatively, separate horizontal gene transfer events could have taken place to insert this nitrogenase into the various ANME clades which would seem to imply a strong selection for this specific nitrogenase clade in the ANME metabolism. Notably, ANME-3 is the only ANME clade lacking a nitrogenase suggesting a requirement for fixed nitrogen, such as porewater ammonium (160).

#### ANME-1 genomes harbor many FrhB/FdhB/FpoF paralogs

Since F_420_ is an electron carrier expected to be used by ANME we examined the diversity of genes predicted to be involved in F_420_ redox reactions. A number of proteins that interact with F_420_ have been discussed above, and many of them share homologous subunits that carry out F_420_ redox chemistry such as FrhB (F_420_ dependent hydrogenase), FdhB (F_420_ dependent formate dehydrogenase) and FpoF/FqoF. Other enzymes utilizing this FrhB-like domain are known such as F_420_-dependent sulfite reductase (Fsr), which has recently been examined in detail in ANME (17). Surprisingly, our examination of this protein family revealed many additional unknown paralogs, particularly in ANME-1 (**Fig 16**). These paralogs were generally monophyletic within the ANME-1, suggesting duplication and neofunctionalization within the order *Syntrophoarchaeales* (**S16 Fig**). The gene clusters containing these FrhB paralogs were often well conserved, and seemed to contain genes that code for enzymes not generally expected to interact directly with F_420_. For example, FrhB6 is mainly found in gene clusters containing pyruvate ferredoxin oxidoreductase, the enzyme responsible for the reductive carboxylation of acetyl-CoA to form pyruvate. Similarly, FrhB7 was found in gene clusters with CODH alpha and epsilon subunits. These subunits of the larger CODH/ACS complex are responsible for the carbon monoxide dehydrogenase activity (161).

**Fig 16.**
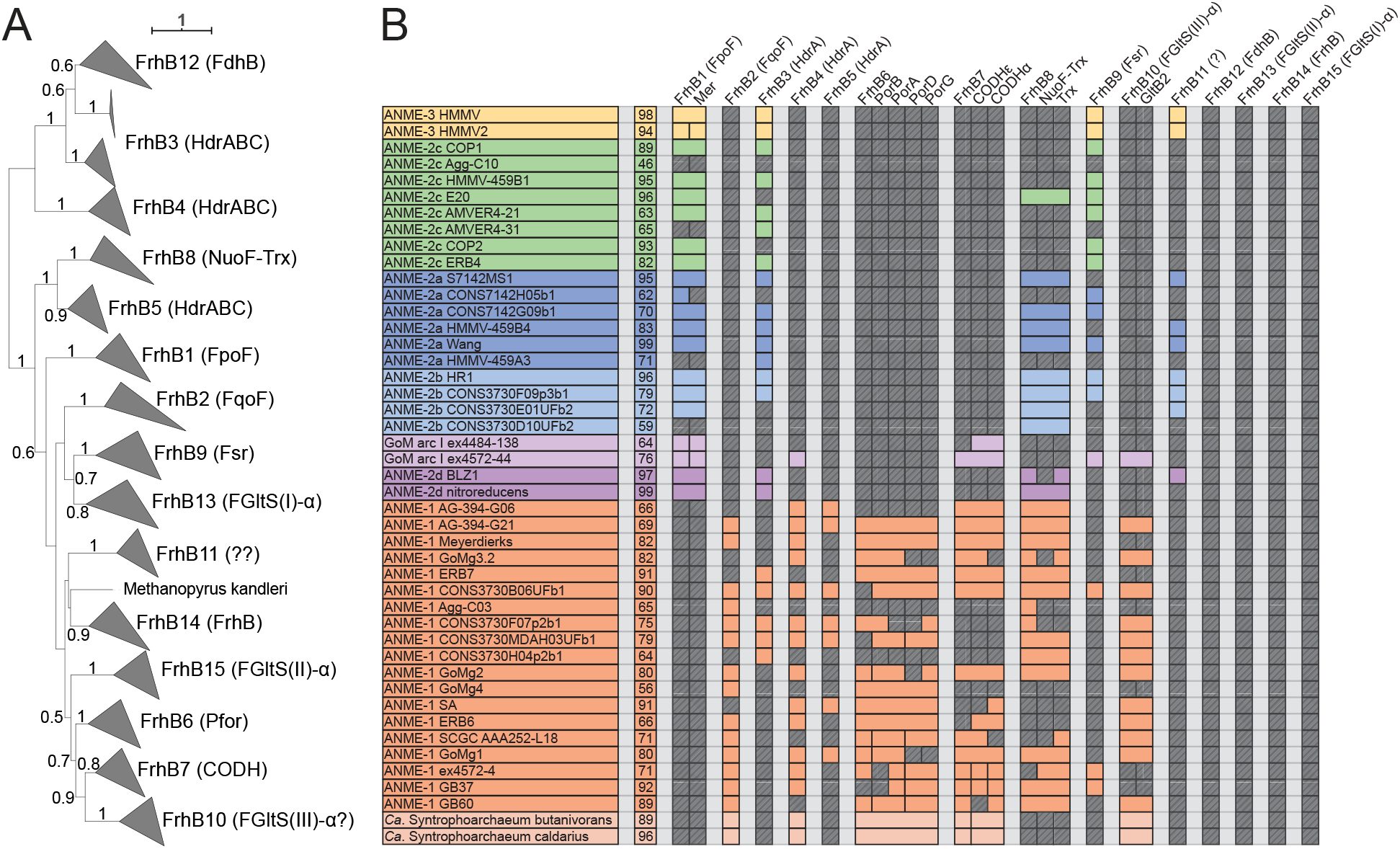
Diversity of FrhB family proteins. Phylogeny and distribution of the FrhB paralogs recovered in our ANME genomes. (**A**) Phylogenetic tree built from all FrhB paralogs in ANME and methanogenic archaea. Major groups have been collapsed (for full tree see **S16 Fig**). Label in parenthesis describe conserved gene cluster; FdhB: F_420_-dependent formate dehydrogenase, FrhB: F_420_-dependent hydrogenase, FqoF: F_420_:quinone oxidoreductase, Fpo: F_420_:phenazine oxidoreductase, HdrA: in cluster with HdrA genes, NuoF-Trx: in cluster with NuoF and thioredoxin genes, Fsr: F_420_-dependent sulfite reductase, FGltS: F_420_-dependent glutamate synthase, Pfor: in cluster with pyruvate ferredoxin oxidoreductase genes, CODH: in cluster with carbon monoxide dehydrogenase alpha and epsilon subunits. Only branch support values >50% are displayed. Tree scales represent substitutions per site. Tree construction parameters are found in the Materials and Methods section. Alignment and tree files can be found in **S1 Data**. (**B**) Presence of various FrhB paralogs in different ANME genomes. Colored boxes represent presence of various nitrogenase related genes in ANME genomes. Missing genes are represented by gray boxes with diagonal line fill. Numbers in the second column represent genome completeness. When genes are together in a gene cluster their boxes are displayed fused together. Gene accession numbers can be found in **S2 Data**.

If these two enzyme systems form complexes with the novel FrhB paralogs as this gene clustering suggests, then they may be operating only in the oxidative direction, using electrons from the CO/CO_2_ couple or the oxidative decarboxylation of pyruvate to reduce F_420_H_2_. Biochemical precedent seems to indicate that F_420_H_2_ cannot be an electron donor for these reactions under physiological conditions. It was initially thought that pyruvate oxidoreductase responsible for the anabolic formation of pyruvate could utilize F_420_H_2_ as its physiological electron donor based on experiments with crude cell extracts (162). Subsequent studies that purified the pyruvate oxidoreductase enzyme found F_420_ to not be a substrate as an electron donor or acceptor (163). Instead, pyruvate oxidoreductase interacted with ferredoxin, a finding consistent with previous results from bacteria. This result also is also more consistent with bioenergetic arguments, the pyruvate/acetyl-CoA+CO_2_ redox couple is expected to be around -500mV and therefore should not be able to be reduced by F_420_H_2_ with a midpoint in the vicinity of -340mV. Thus the pyruvate dependent reduction in crude cell extracts was attributed to secondary reduction of F_420_ by Fd (163). The gene clusters here may represent a dedicated system of F_420_ reduction associated with the breakdown of multi-carbon compounds. Whether this system is used in the breakdown of carbon storage molecules, or is an important part of ANME catabolism remains to be determined.

Additional noteworthy ANME-specific FrhB paralogs include FrhB10, which are found in gene clusters with various subunits of glutamate synthase, potentially representing a third putative group of F_420_-dependent glutamate synthases (164). FrhB8 are found in gene clusters with divergent NuoF homologs and thioredoxin genes, and are almost exclusively found in ANME groups (**Fig 16B**, **S16 Fig**). While our limited transcriptome data shows many of these FrhB homologs to be expressed at moderate levels, FrhB7 in ANME-1 GB60 is an exception, with expression levels near those of the methanogenesis pathway (**S3 Data**). As with the diversity of HdrA-containing gene clusters, this expansion of FrhB-containing gene clusters in the ANME-1 points to a wide range of novel electron transport capabilities in ANME-1.

#### Extensive Dockerin/Cohesin domain-containing proteins

Recently a protein domain study of three marine ANME genomes reported that ANME-1 and ANME-2a contained a surprising number of proteins predicted to have dockerin or cohesin domains (42). These domains are best known from their role in the construction of the cellulosomes which are large, multiprotein complexes that bind and degrade extracellular cellulose in *Clostridia* and other cellulose degrading bacteria (165). Dockerin and cohesin domains form strong bonds with one another, and by encoding multiple sets of complementary dockerin and cohesin-containing proteins many copies of enzymes with cellulolytic activity can be linked to other proteins containing cellulose binding domain as well as anchored to the cell surface. These multiprotein complexes can be bound to the cell by anchor proteins that contain cohesion and S-layer homology (SLH) domains or via covalent linkage to lipids through the action of sortases.

Cohesin and dockerin domains have been found in diverse bacteria and archaea that are not thought to be involved with cellulose degradation, suggesting functions beyond the well characterized ones in clostridia (166). Early work found proteins containing dockerin and cohesin domains in *Archaeoglobus* and these archaeal versions have been verified to perform specific strong dockerin-cohesin bonds (167, 168). However archaea that contained dockerin or cohein domains did not contain large so-called “scaffoldin” proteins with multiple copies of cohesin domains which are required for the formation of large multimeric complexes (166). Since dockerin and cohesin domains simply facilitate the linkage of functional domains, the implications of these domains are not clear simply by their presence in the genome.

Dockerin and cohesin domain-containing proteins in bacteria vary considerably between closely related species in the number and identity of additional domains, and the proteins in ANME are similarly variable, but a few consistent trends are apparent. In members of ANME-2a, ANME-2c and ANME-1 there are proteins that consist of a dockerin domain and multiple cohesin domains, up to as many as five. These scaffoldin-like proteins therefore have the ability to localize multiple copies of whatever proteins contain their cohesins’ complementary dockerin domains. Common dockerin containing proteins in ANME are the periplasmic substrate-binding components of ABC transporters of various types, such as the nickel/dipeptide/oligopeptide (NikA/OppA) system and the TroA system used for metal ion uptake (169, 170). In most ANME-2c genomes there are proteins encoding S-layer domains and dockerin domains, suggesting dockerin/cohesin pairing may be important for attaching functional proteins to the outer layer of the cell and possibly mediating interactions with bacteria. Using experimental methods to determine which dockerin and cohesin domains bind to each other will lead to a better understanding of composition of the ANME extracellular space. A complete list of ANME proteins containing dockerin or cohesin domains can be found in **S4 Data**.

#### Phage-like protein translocation structures

Phage-like protein translocation structures (PLTSs) are large multiprotein complexes that share structural and functional similarities to type VI secretion systems, pyocins, and the *Serratia entomophila* antifeeding prophage described in a broad survey of microbial genomes based on sequence identity and synteny (171). This broad family of related complexes have a wide variety of functions, from neighboring cell lysis, to morphologic transformation of targeted eukaryotic cells (172), stabilizing symbiotic interactions (173) and mediating sibling conflict in the multicellular aggregate bacterium *Myxococcus xanthus* (174). PLTSs are anchored in the cytoplasmic membrane and upon contraction of the sheath proteins extrude a protein complex with a sharp spike that penetrates neighboring cell membranes and can deliver various effector proteins (175). Recent reviews of the known structure and function of these complexes highlight the conserved features which include phage baseplate-like proteins, sheath proteins, AAA+ ATPases, LysM-motif proteins for peptidoglycan binding, and VgrG and PAAR-domain spike proteins (176).

Gene clusters related to these systems were identified in all ANME groups with gene synteny similar to that of *Methanomethylovorans hollandica* that was described in a recent review (171) (**Fig 17**). This gene cluster was relatively well expressed in ANME-2c (**S3 Data**), and could potentially play an important role in the symbiosis between ANME and partner bacteria, or alternatively defend AOM consortia from invasion. Interestingly, PLTS clusters from closely related ANME-2a and 2b were very different from one another, with the ANME-2b sequences much more closely related to the ones found in ANME-1 (**Fig 17C**). In the two ANME-2d genomes available, one encoded the version like ANME-2a, ANME-2c and ANME-3, while the other ANME-2d genome encoded a version that was very similar to ANME-1 and ANME-2b. This difference is most apparent in the PAAR domain spike proteins, with ANME-2b much more similar to the PAAR domain proteins encoded in ANME-1 (**Fig 17C**). It is interesting that such closely related ANME groups encode such different PLTS systems, and suggests that ANME-1 and 2b may have similar interactions with organisms (possibly syntrophic SRB partners) that are quite different than those in association with ANME-2a, 2c and 3.

**Fig 17.**
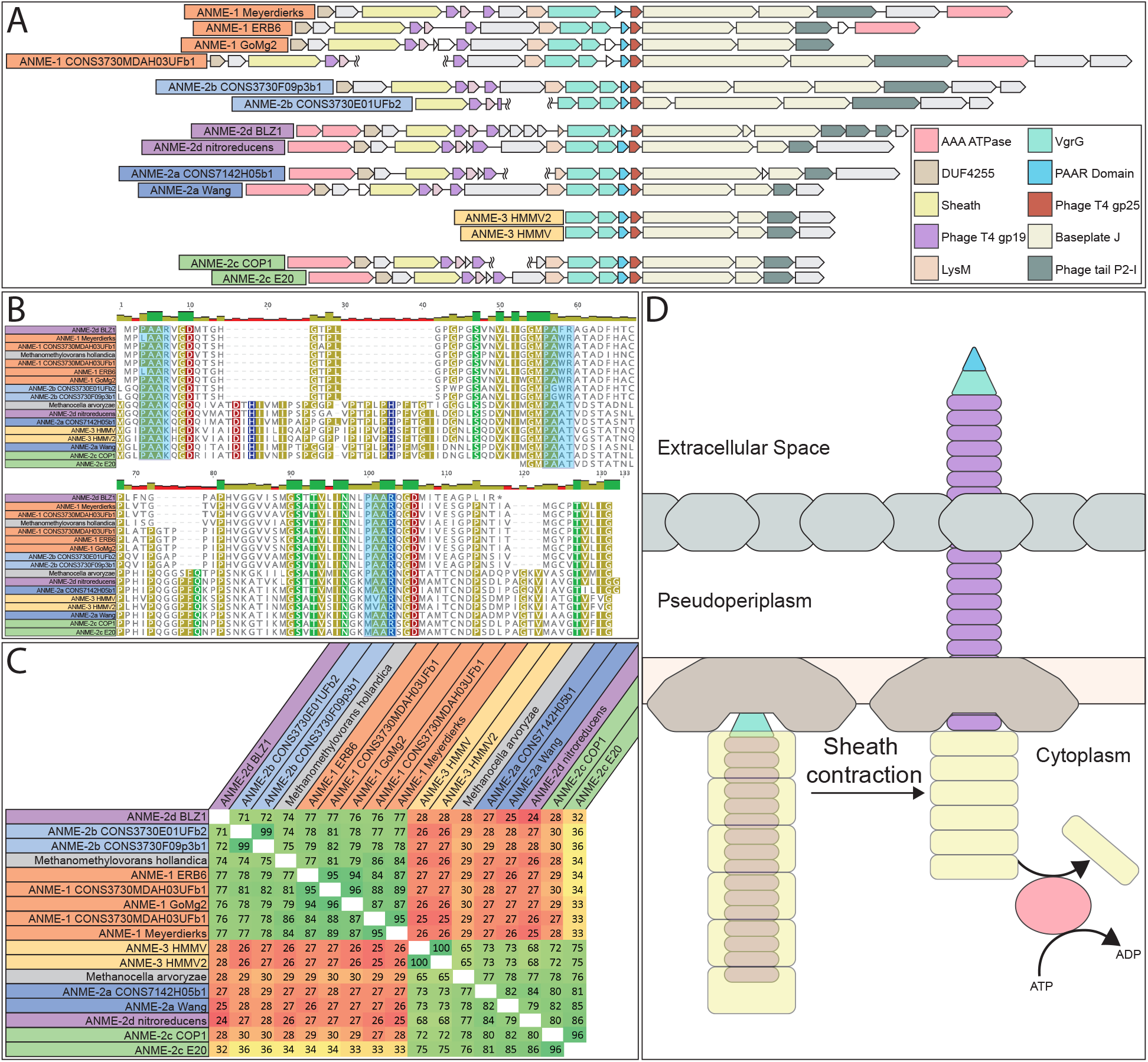
Phage-like protein translocation structures. (**A**) PLTS gene clusters in ANME genomes. (**B**) Alignment of PAAR domain spike proteins. PAAR motifs highlighted in blue. Alignments were made using muscle 3.8.31 with default settings, and alignment file can be found in **S1 Data.** (**C**) amino acid identity of PAAR domain proteins highlighting two clear groupings. Note: closely related ANME-2a and 2b genomes contain PLTS structures belonging to different clusters. (**D**) Schematic of PLTS function.

## Discussion

### The evolution and conserved metabolic features of marine ANME archaea

Our investigation of this expanded set of ANME genomes revealed key features present in ANME that are absent in their methanogenic relatives. While methanogens and methanotrophs use the same set of core enzyme for their catabolic C1 metabolisms, the key difference occurs in the way in which electrons are input or output from this pathway. A unifying feature of the ANME archaea appears to be a multitude of multiheme *c*-type cytochromes that are absent in methanogens. This expansive repertoire of MHCs, some harboring 30+ heme binding domains in ANME-2 and 3, specific cytochrome *c* maturation machinery, and associated conserved hypothetical proteins are features only shared by certain members of the *Archaeoglobales* that are known to conduct EET. The large ANME-MHCs with S-layer fusions and the repurposing of the cytochrome *b* subunit of membrane bound hydrogenases into a potential methanophenazine:cytochrome *c* oxidoreductase complex (Mco) remain some of the clearest examples of bioenergetic novelty in ANME compared to their methanogenic relatives. The study of EET in symbiotic associations is relatively new, and there are no specific marker proteins yet identified that definitively confer EET ability. There is however a general pattern in that the genomes of EET-capable organisms contain many, often exceptionally large, MHCs, as well as systems enabling the transfer of electrons through the various outer layers of the cell. ANME appear to contain all of these genomic features, and this clear EET potential represents an important physiological difference between ANME and methanogens. This electron transfer system, not any modification to the C1-handling enzymes themselves, is likely the major determining factor in the directionality of the “reverse methanogenesis” pathway.

Another common feature of ANME genomes that set them apart from their methanogenic counterparts is the high abundance and diversity of soluble Hdr complexes with divergent formate dehydrogenase (FdhAB) homologs. This is especially true for ANME-1 that lack certain features often found within *Methanosarcinaceae* including HdrDE and, in most cases, the Rnf complex. It is likely that these soluble Hdr complexes are involved in electron recycling and transfer processes in at least ANME-1, but also ANME-2 and 3 based on their conservation and expression levels. So far, experimental evidence suggests that formate is not an electron carrying intermediate in AOM, and the canonical formate transport protein (FdhC) is absent. Hence, the specific substrate and function of these well-conserved complexes remains a fundamental question in ANME metabolism remaining to be addressed. Regardless of the nature of the electron acceptor utilized by these Hdr complexes, the reverse methanogenesis model requires that their combined effect is a net oxidation of CoM-SH/CoB-SH, presumably through electron confurcation.

This expanded set of ANME genomes also helps us better understand their evolutionary history. The paraphyletic nature of ANME suggests that the transition between methanogenic and methanotrophic metabolisms has occurred multiple times. The order of these events is not immediately evident, as multiple scenarios could lead to the observed pattern of metabolism and gene content. This is particularly true in the case of ANME-3, which contains multiple features identified in ANME-2 genomes including large MHC proteins, specific duplicated CcmF proteins, the *b*-type cytochrome associated with Rnf, divergent FdhAB associated with HdrA genes, among others. The presence of these bioenergetic genes within ANME-3, which appear to be the most recently diverged from within the *Methanosarcinaceae*, strongly suggests that this repertoire is critical to the methane oxidation phenotype. With the present data we do not believe there is a clear parsimony-based argument to distinguish between the possibilities that these ANME-specific features represent a gain of function through horizontal gene transfer into the ancestor of ANME-3, or whether they were lost in the methanogenic members of the *Methanosarcinaceae*.

It has long been assumed that ANME evolved from methanogenic ancestors and our current phylogenies support this scenario, as many groups of hydrogenotrophic methanogens emerge from the archaeal tree before the rise of the deep branching ANME-1 (**Fig 1**). With this expanded set of ANME genomes, another interesting evolutionary possibility presents itself: the entire group of methylotrophic methanogens within the *Methanosarcinaceae* may be derived from a methanotrophic ancestor. These methanogens run six of seven steps of the methanogenesis pathway in reverse. This fact was used early on to suggest “reverse methanogenesis” was reasonable for ANME, since they would only have to reverse one additional step. The order of our discovery of these metabolisms may not reflect the order with which these metabolisms evolved, and the complete reversal of the methanogenesis pathway may date back to the last common ancestor of ANME-1 and the other ANME. In this evolutionary scenario, methylotrophic methanogenesis in the *Methanosarcinaceae* evolved by the simple acquisition of methyl transferases and the subsequent loss of ANME-specific systems of EET. Instead of transferring electrons from methane oxidation to the outside of the cell, the *Methanosarcinaceae* electrons are funneled from methyl group oxidation back into the cytoplasm to reduce heterodisulfide.

### The “Methanoalium” group of ANME-1 and the potential for methanogenesis in ANME

The clade within ANME-1 represented by the SA and GoM4 genomes is an exceptional group which requires further detailed study. This group has previously been referred to as the “freshwater ANME-1 clade” and has been found in 16S rRNA gene surveys of various terrestrial and marine environments (23, 177, 178). A recent report constructed the first genome from this group of ANME-1 from a Tibetan hot spring (THS) (12). We recognized similar characteristics between the THS genome and the GoM4 and SA genomes reconstructed here, including the bacterial-type Rnf and the absence of MHC. Although not mentioned previously, we find the same MvhA-type hydrogenase in the ANME-1-THS genome and an absence of cytochrome maturation machinery. As a final check on the novel characteristics of this unusual ANME-1 clade, we reconstructed one additional genome from a recent metagenomic dataset of the Lost City hydrothermal field. This site has reported 16S rRNA genes and McrA sequences belonging to this freshwater ANME-1 clade (12, 177–179). This additional Lost City ANME-1 MAG was 88% complete with 2% contamination and notably had all the hallmark genomic features observed in GoM4, SA, and THS (**Fig 18**).

**Fig 18.**
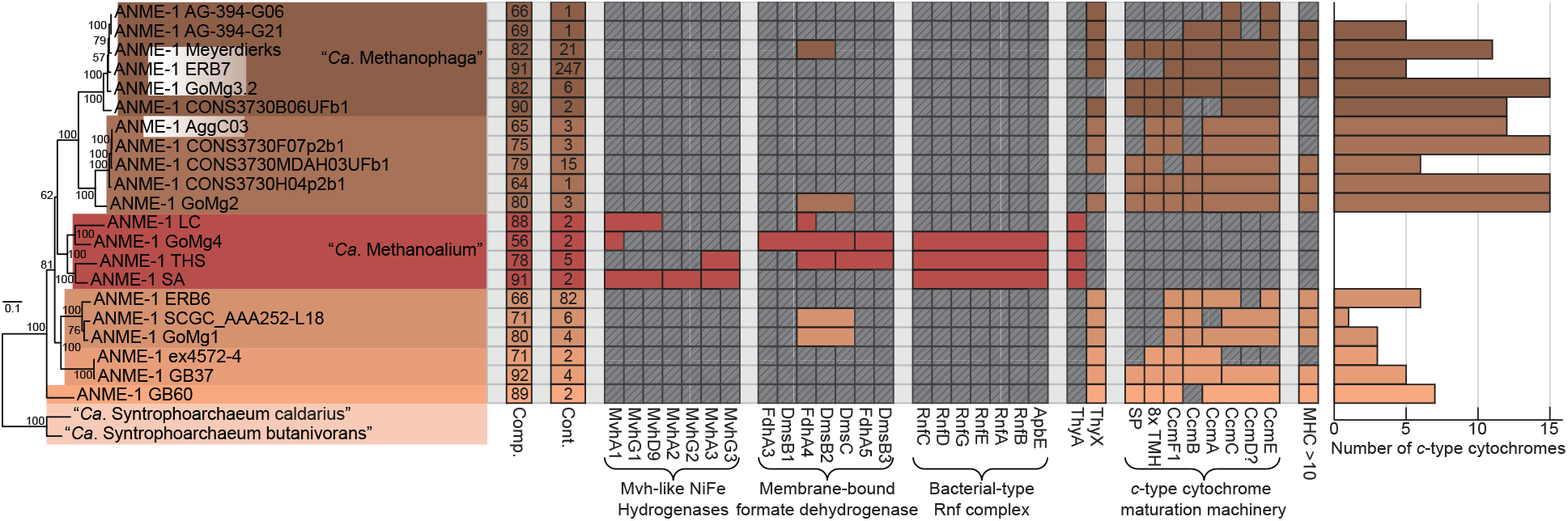
Comparison between “*Ca*. Methanoalium” and other ANME-1 genera. Phylogenomic tree based on concatenated marker proteins highlighting individual ANME-1 genera, two of which have been assigned “*Candidatus*” names. Estimated genome completeness and contamination are shown in the first and second columns. Comparison of the presence of hydrogenase, membrane-associated formate dehydrogenases, Rnf complexes and cytochrome maturation machinery highlights important differences in electron flow between these genera. Genomes encoding MHC with more than 10 CxxCH heme-binding motifs are marked in the final column. Bar chart on the far right demonstrates the number of *c*-type cytochromes per genome. Only branch support values >50% are shown for clarity. Tree scales represent substitutions per site. Tree construction parameters are found in the Materials and Methods section. Alignment and tree files can be found in **S1 Data**, and gene accession numbers can be found in **S2 Data**.

A number of investigations of ANME-dominated environments have concluded that some ANME lineages, particularly ANME-1, maybe have net methanogenic capabilities (180–183). Genomic analysis alone cannot falsify the hypothesis that ANME archaea can gain energy from methanogenesis, due to the similarities in the main methanogenic pathway enzymes. Yet, incubations of methane seep samples with methanogenic substrates has only succeeded in stimulating the growth of low abundance traditional methanogens, never ANME (93). The evolution between methanotrophy (e.g. ANME-2a, 2b and 2c), and methanogenesis (canonical members of the *Methanosarcinaceae*) as well as the apparently recent transition back to methanotrophy in ANME-3 appears to include biochemical innovation and genomic adaptation which occurs on evolutionary timescales, but not ad hoc in the environment. Likely, in a group as old and diverse as ANME-1 (representing at least 6 genera), such a transition may have also occurred. We hypothesize that this clade of ANME-1 (SA, GoM4, THS, and LC) that encode for hydrogenases and lack *c*-type cytochromes may be bona fide methanogens. However, the little evidence currently available for representatives of this clade from the terrestrial subsurface suggests that members of this clade may carry out methane oxidation as well (23). Whether this process occurs in association with a syntrophic partner is currently unknown. What is clear is that this clade’s genomic content and environmental distribution sets them well apart from other ANME. We propose the genus “*Ca*. Methanoalium” for the “freshwater ANME-1 clade” in recognition of the differences in their metabolic potential and unusual environmental distribution (see **S1 File** for etymology).

### Anabolic independence of the ANME archaea from their syntrophic partner

The tight metabolic coupling between syntrophic ANME and SRB could be expected to result in reductive biosynthetic pathway loss in one of the partners as predicted in the “Black Queen Hypothesis” (137). Unfortunately, the partner SRB have not yet been cultured, and thus it remains unknown if these show specific genomic adaptations to a consortia lifestyle. However, we do not observe obvious evidence of this occurring at the level of major ANME lineages for amino acids. If this sort of reductive evolution does occur in the ANME-SRB symbiosis, it may be occurring at the species or strain level, and would be very difficult to confidently detect in coarse-grained genomic analysis with partial genomes. More detailed investigations of the question of complementary biosynthetic pathways between syntrophic partners will be best studied in sediments or enrichment cultures that are dominated by a single ANME-SRB pairing, where near complete genomes of both partners can be generated. But, on the broad scale of ANME evolution to their syntrophic lifestyle, there does not seem to be a concerted loss of anabolic independence in any of the major lineages. This result is distinct from that described in some other syntrophic communities such as those performing the anaerobic oxidation of hexadecane (136).

### Biogeochemical and microbiological consideration of ANME carbon signatures

Bacterial methylotrophs and methanotrophs have traditionally been defined as organisms that derive both their carbon and energy from the oxidation of C1 compounds or methane, respectively (184, 185). ANME were originally called methanotrophs because it appeared that their energy metabolism was based on methane oxidation, and the ^13^C-depleted isotope signature of their lipids was thought to be evidence of the assimilation of ^13^C-depleted methane carbon. Biogeochemical studies additionally show that the respiration of methane to CO_2_ dominates carbon turnover in most ANME environments, and thus that the dissolved inorganic pool of carbon is mostly methane-derived. As a consequence, the carbon assimilated for biomass production is only a few percent of the methane oxidized and thus also methane-derived from a biogeochemical perspective (186, 187). However, in vivo isotope-probing experiments showed that ANME biomass is effectively labelled by ^14^CO_2_ or ^13^CO_2_, leading to the conclusion that ANME assimilate a mixture of CO_2_ and CH_4_ (187, 188), or almost only CO_2_ (189). Short pulse-chase experiments with ^14^CO_2_ and ^14^CH_4_ also support this latter interpretation (93).

Microbiological considerations for the classification of this C1 metabolism were best articulated by Leadbetter and Foster in the context of aerobic methane oxidizing bacteria (190):

> *“Although there is no universally accepted definition of the nature of the autotrophic mode of life, the ability to grow at the expense of CO_2_ as the exclusive source of carbon for cell synthesis remains as the cardinal consideration in the concept of an autotroph*.
>
> The question needing study is whether, during growth on methane, these bacteria dehydrogenate the methane molecule to CO_2_ and “active” hydrogen, following this with a reductive assimilation of the CO_2_ with “active” hydrogen.”

The dominant bacterial methanotrophs in most environments assimilate methane-derived carbon at the oxidation state of formaldehyde for a large portion of their biomass using either the serine or ribulose monophosphate pathway (RuMP) (185). Some organisms have been discovered that use the oxidation of C1 compounds for their source of energy, but use a Calvin-Benson-Bassham (CBB) cycle for the fixation of CO_2_. These organisms presented a problem for nutritional nomenclature because the definition of methylotroph/methanotroph normally involves the incorporation of cell carbon. For methylotrophs, *Paracoccus denitrificans* uses a CBB cycle when growing on C1 compounds, and this has earned them the label of “autotrophic methylotroph” (185, 191). More recently, the methane-oxidizing bacteria *Methylacidiphilum fumariolicum* and “*Candidatus* Methylomirabilis oxyfera” have been found to operate a CBB cycle for carbon assimilation from CO_2_ (192, 193). These authors have taken a similar naming scheme, referring to

> *M. fumariolicum* as an “autotrophic methanotroph” (192). These examples from bacteria suggesting that the “methanotroph” label should be conserved for ANME, and the only question remaining is whether to adopt the “autotroph” label as well.

The predominant labelling of ANME biomass by ^14^CO_2_ or ^13^CO_2_ is intriguing, and suggests that they could either act as true autotrophs, or produce biomass by a combination of CO_2_ assimilation and the incorporation of carbon at other oxidation states, as has been described for some other archaea and methanogens. (194, 195). All ANME genomes encode the Wood-Ljungdahl pathway which can be used for carbon fixation, but the methyl branch of this pathway is used for methane oxidation. One option to explain these isotope labelling results is that ANME are autotrophic methanotrophs, and have an additional CO_2_ fixation pathway so that anabolic and catabolic C1 reactions can run in opposite directions simultaneously, such as the CBB cycle in *M. fumariolicum*. However, we have not found evidence for a second carbon fixation pathway in the ANME. Another possibility could be a complete assimilatory Wood-Ljungdahl pathway utilizing H_4_F as found in bacteria, but the H_4_F-interacting enzymes found in some ANME-2 and -3 would not allow them to activate CO_2_ via formate, and are instead are expected to be involved in specific anabolic reactions well downstream of acetyl-CoA production. If the ANME archaea are autotrophs, it remains to be determined how this is carried out from a biochemical perspective.

Another plausible explanation for the predominant labelling of ANME biomass by ^13^CO_2_ could be carbon back flux along the methyl branch on the Wood-Ljungdahl pathway to the point of CH_3_-H_4_MPT, against the net flux of carbon. Metabolisms operating at close to equilibrium can present challenges to isotope labeling studies because the forward and reverse fluxes through that pathway approach equal values (35). AOM operates as close to equilibrium as is thought to be possible to support life, and most of the steps of the methyl branch of the Wood-Ljungdahl pathway are essentially at equilibrium (37). Under these conditions it seems quite possible that the isotope label of the DIC in these incubations could back flux up through the methyl branch of the Wood-Ljungdahl pathway to the point of CH_3_-H_4_MPT, even if the net reaction is in the oxidative direction. Further experiments are required to distinguish between these possible explanations, and determine whether ANME should be referred to as “autotrophic methanotrophs” or simply “methanotrophs”.

### MetF, F_420_-dependent NADP reductase and electron bifurcation complexes

There remains no clear explanation for what evolutionary or bioenergetic pressures may have led to the loss of Mer and presumed replacement by MetF in ANME-1. An important challenge posed by this switch is that MetF interacts with NADPH instead of F_420_H_2_, and it is not immediately apparent how electrons from NADPH are harnessed for energy production in ANME-1. In one sense, this is similar to the case of methanogens that use multicarbon alcohols as electron donors for CO_2_ reduction (196). Some of these methanogens contain alcohol dehydrogenases that utilize F_420_ as the electron acceptor, and therefore the electrons liberated from the alcohols as F_420_H_2_ can immediately be used in the methanogenesis pathway. Other methanogens however contain alcohol dehydrogenases that are NADPH specific and so a secondary oxidoreductase is needed to transfer NADPH electrons onto F_420_. It was discovered that in this latter group of methanogens, high activities of F_420_H_2_:NADP oxidoreductase (Fno) could be found, and the protein responsible was purified, characterized and sequenced (196, 197). If an Fno enzyme was expressed at sufficiently high levels in ANME-1 then the electrons on NADPH derived from MetF could be fed back into the F_420_ pool for catabolism, just as in the alcohol-oxidizing methanogens.

Remarkably, ANME-1 universally lack homologs of Fno although they are present in all other ANME and methanogens that we examined. In most methanogens Fno is present for the reverse reaction, the production of NADPH for anabolic reactions. This conspicuous absence in ANME-1 of what seems to be a nearly universally conserved mechanism of hydride transfer between the F_420_H_2_ and NADPH pools is noteworthy on its own, but is particularly interesting when considered in the context of the Mer/MetF switch. If ANME-1 produce NADPH from MetF, they will have more electrons than they need for anabolism through this reaction since this amounts to a quarter of all methane-derived electrons. If this is how MetF is being utilized in ANME-1, then how these NADPH electrons are transferred back into an energy conservation pathway is an important open question. Most ANME-1 also contain a homolog of what we have designated HdrA9, 10 or 11, which constitute HdrA with large insertions with high sequence identity with NfnB. This subunit interacts with NADPH in the well characterized Nfn systems (78), so if MetF produces NADPH, these modified HdrA/NfnB fusions may represent a novel path back into the catabolic energy metabolism of ANME-1, confurcating electrons from NADPH and CoM-SH/CoB-SH.

The prevalence of many diverse copies of HdrABC systems in ANME-1 may offer an alternative explanation for how MetF-derived electrons are directed back into central catabolism. Recently a protein complex containing MetFV, HdrABC and MvhD was discovered in *Moorella thermoacetica* that is suggested to carry out electron bifurcation with NADH, CH_2_=H4F and other unknown reactants (52). Genomic analysis revealed the genes coding for these six proteins were found in a gene cluster together in *M. thermoacetica* and our examination of this HdrA homolog in this complex is much like HdrA 3 and 4 described above, with two fused HdrA domains with complementary Cys197/N-terminal domain loss. While none of the ANME-1 genomes analyzed here contain MetF in a gene cluster with HdrA homologs, the two “*Ca.* Syntrophoarchaeum” genomes both contain gene clusters containing MetFV, HdrA and MvhD. If MetF catalyzes the CH_3_-H_4_MPT oxidation step in ANME-1, it may form complexes with one of the many HdrA paralogs. This would allow electrons to funnel directly into a bifurcation/confurcation reaction without passing through the F_420_H_2_ pool. In any case, electron flow through ANME-1 is likely to involve a significant amount of novelty that will require detailed biochemical studies to test these genomic predictions.

### An energetic argument for both chemical diffusion and direct electron transfer in ANME-SRB syntrophy

Two strategies for syntrophic electron transfer could be widespread in the ANME-SRB syntrophy based on the conserved features of energy metabolism described here, EET utilizing the abundant MHC proteins, as well as diffusion of soluble electron carriers potentially produced by the diverse HdrABC complexes. A possible explanation for the conservation of these two systems in the genomes of ANME is that a mixed electron transfer through both systems may best suit an equal division of energy between the ANME and their sulfate reducing syntrophic partners.

If ANME-2a were to use Fpo, Rnf and HdrDE to oxidize F_420_H_2_, Fd^2-^, and CoM-SH/CoB-SH, our expectation would be that all eight electrons from methane oxidation would end up on MpH_2_ (**Fig 19A**). This would conserve abundant energy for the ANME through the Fpo and Rnf complexes, but the redox potential of the Mp/MpH_2_ couple (E_0_’ = -165mV) is more than 50mV more positive than the SO_4_^2-^/HS^-^ redox couple (E_0_’ = -217mV). If all electrons were to pass through MpH_2_ to the SRB their entire metabolism would be endergonic by about 40kJ/mol sulfate. For an equitable sharing of energy between ANME and SRB, the average redox potential of electrons during transfer should be approximately halfway between the CH_4_/CO_2_ (E_0_’ = -240mV) redox potential and the SO_4_^2-^/HS^-^ couple (E_0_’ = -217mV). The MpH_2_ pool would need to be more than 90% reduced to drop its midpoint potential to ∼ -230mV from -165mV, but this would render the HdrDE reaction impossible to run in the CoM-SH/CoB-SH oxidizing direction since the heterodisulfide E_0_’ is approximately 100mV more positive than this. The situation would be even more difficult for ANME-1 which are predicted to use a menaquinone which is very likely at a higher potential than MpH_2_, requiring the membrane-bound electron carrier pool to be greater than 99% reduced for the SRB to yield any energy accepting all eight electrons from menaquinone. In addition, whatever the MpH_2_ or QH_2_ pool midpoint potential is, it will be lower than the potential the SRB will receive electrons at, since the transfer from ANME to SRB through MHC must be down a redox gradient to maintain electron flow.

**Figure 19.**
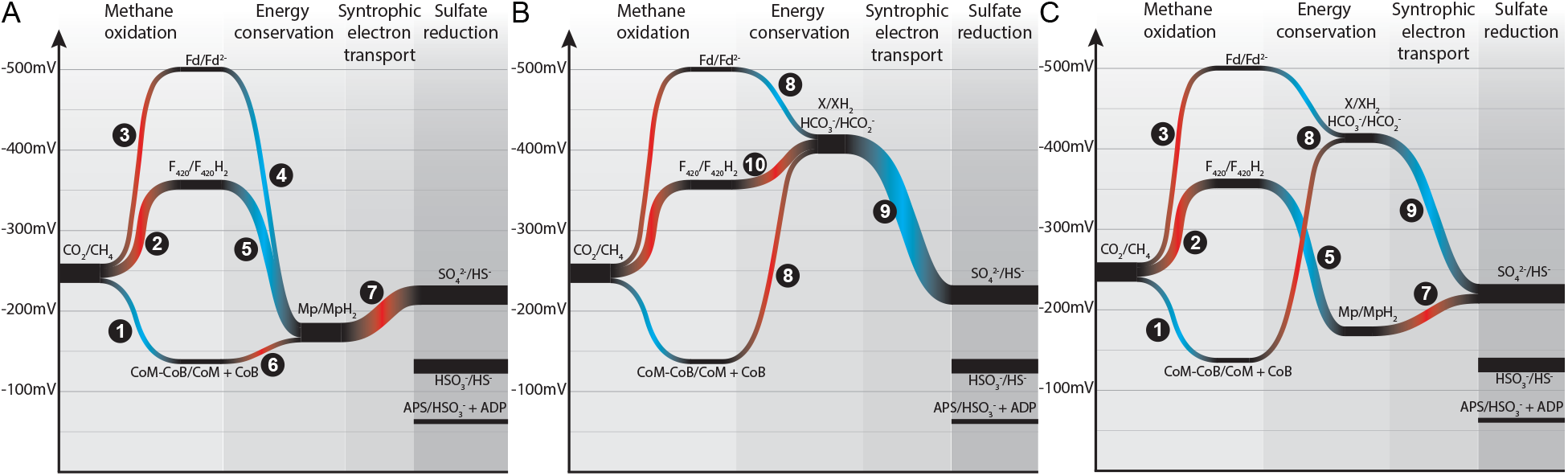
Energetics of mixed electron transfer. Energy flow diagrams showing the redox potential (y-axis) of electrons as they travel from methane through the ANME energy metabolism to the SRB partner (x-axis is reaction progression). Width of paths correspond to number of electrons. Endergonic electron flow (uphill) shown in red, exergonic (downhill) in blue. Steps labelled with numbers are carried out by the following enzymes; 1: Mcr, 2: Mer/Mtd, 3: Fwd/Fmd, 4: Rnf, 5: Fpo/Fqo, 6: HdrDE, 7: Mco and ANME-MHC, 8: HdrABC and ANME-specific FrhB, FdhA, 9: uncatalyzed diffusion of electron carrier 10: FdhAB, possibly confurcation through HdrA complexes. X/XH_2_ represent a hypothetical low-potential electron carrier that could be used as a diffusive electron shuttle. (**A**) electron transfer based entirely on EET from the methanophenazine pool. (**B**) electron transfer based entirely on soluble electron shuttle produced in the cytoplasmic space. (**C**) Mixed model with half of the electrons passing through each pathway.

On the other hand, if all methane-derived electrons were used to produce a low-potential soluble electron carrier like formate through the action of cytoplasmic HdrA-based confurcation then the SRB could readily grow with electrons at this potential for sulfate reduction. In this scenario however the ANME’s entire energy metabolism would contain no steps that could be coupled to energy conservation. Energy would be lost at the sodium pumping Mtr step, and there is no mechanism for recovering this energy since these soluble Hdr/Fdh complexes do not produce transmembrane potential or carry out substrate-level phosphorylation (**Fig 19B**).

A combined model of cytochrome-based EET and diffusion of low potential intermediates could alleviate the need for extremely reduced quinol pools to overcome the energetic inequalities between ANME and SRB, and may help explain the conservation and high expression of these two seemingly opposing methods of syntrophic electron transfer (**Fig 19C**). Four electrons transferred from two F_420_H_2_ through MpH_2_/QH_2_ and MHC-based EET could provide just enough energy to the ANME to overcome the losses at Mtr and support growth. This would utilize the Fpo and Fqo homologs that are universally conserved in all ANME groups. If the remaining four electrons are confurcated to formate or a similarly low potential electron carrier then the effective midpoint potential of this compound could be tuned to reach a balance where the effective redox potential of all eight of the transferred electrons were ∼ -230mV, with the ANME conserving energy via the four electrons passing through Fpo/Fqo, and the SRB conserving energy through the four electrons on the lower potential soluble electron carrier. The SRB would need to continue consuming the higher potential EET electrons, or else the lower potential electrons would cease flowing. Experimental results argue against formate as the predominant free extracellular intermediate in AOM (84, 91–93), but perhaps another small molecule or diffusible organic substrate of sufficiently low redox potential could solve this energetic puzzle.

## Conclusion

In proposing that AOM might be explained by methanogens running in reverse and transferring intermediates to a sulfate reducing bacterial partner, Zehnder and Brock concluded (41):

> “*We do not pretend that our hypothesis provides a complete explanation for the geochemical observations in anoxic marine environments, but our results provide reasonable evidence that methane is not biologically inert in strict anaerobic ecosystems.*”

In a similar spirit, we present these genome-based models of ANME metabolism with the understanding that they represent a series of hypotheses that require biochemical, genetic or physiological experiments performed on purified ANME or ANME-encoded proteins to test. These models in the context of our expanded phylogenetic framework will help address important remaining questions in ANME-SRB syntrophy. Future studies will need to focus on specific ANME subgroups, preferably at the species level, to address questions of energy conservation mechanisms and adaptation to specific niches in anoxic marine environments. Novel methods of analysis such as electrode cultivation may provide important new insights into predicted EET-capabilities. Features specifically supporting syntrophy, including extracellular structures such as pili, specific interactions using phage-like protein translocation structures, or dockerin-cohesin systems will require careful experimental analysis far beyond genomic studies, but may have significant impacts on our understanding of cell-cell interactions in AOM consortia. We expect that explicit consideration of the carbon and electron transfer pathways in specific ANME clades will help address controversial results such as net methane production and the question autotrophic growth. Our continued investigations into the enigmatic origin and function of AOM is important for understanding of the dynamics of methane fluxes in Earth history, and also for the evolution of the archaea and their syntrophic interactions with bacteria.

## Materials and Methods

### BONCAT sorted aggregate assembly and binning

Single aggregates were sorted using bioorthogonal noncanonical amino acid tagging (BONCAT) combined with fluorescence-activated cell sorting (FACS), an activity-based flow cytometry method from samples described previously (14). Two sediment samples were used, originating from Hydrate Ridge, Oregon, USA (sample #3730), or Santa Monica Mounds, California, USA and incubated at 4°C with the artificial amino acid L-homopropargylglycine (HPG) for 114 days for sample #3730, or 25 days for sample #7142. Anabolically active cells were fluorescently labelled using BONCAT followed by FACS sorting using the standard procedure implemented by the Joint Genome Institute’s (JGI’s) single cell sequencing pipeline. Each individual sorted aggregate was lysed and the DNA amplified using multiple displacement amplification (MDA). Metagenomic data was sequenced, assembled and annotated at the JGI. Metagenomic assembly was performed using the jigsaw pipeline v2.6.2 and annotation performed using the standard IMG annotation pipeline (198). Binning of the two partner microorganisms from these single aggregate metagenomes was performed using VizBin (199).

### Gulf of Mexico (GoM) assembly and binning

Sediment samples from the Gulf of Mexico were retrieved by push coring on March 2015 during RV METEOR expedition ME114/2 (quest dive 362) from a gas and oil seep at the Chapopote asphalt volcano, Campeche Knolls (0-10 cm, GeoB 19351-14, 21°53.993’N; 93°26.112’W) (200). This is a cold seep located in the northern part of the Campeche Knolls field in the Gulf of Mexico at 2925 m water depth. On board, sediments were transferred into a Duran bottle, closed with a gas-tight butyl rubber stopper and headspace was exchanged to pure N_2_. Anoxic aliquots of these samples were shipped at 5°C to the Bigelow Laboratory Single Cell Genomics Center (SCGC; https://scgc.bigelow.org) where sequencing took place as previously described (201). In short, sediment samples were diluted and cells were separated from the sediment by a slow centrifugation step. Afterwards, single cells were sorted by high-speed fluorescence-activated and droplet-based cell sorting (FACS) into three 384-well plates and cell lysis followed using five freeze-thaw cycles and KOH treatment. Then, whole-genome amplification via multiple displacement amplification (MDA) was performed with subsequent cell screening and genome sequencing. Cell screening identifying archaeal cells affiliated with ANME was performed by 16S rRNA-gene tag sequencing, while libraries for genome sequencing were created with Nextera XT (Illumina) and were sequenced with NextSeq 500 (Illumina). Reads were quality trimmed using Trimmomatic v0.32 with the following parameters: ‘-phred33 LEADING:0 TRAILING:5 SLIDINGWINDOW:4:15 MINLEN:36’ and screened for contamination against the human genome assembly GRCh38. Individual single cell datasets were assembled by normalizing the kmer coverage of the reads using kmernorm 1.05 (https://sourceforge.net/projects/kmernorm/) using the parameters ‘-k 21 -t 30 -c 3’ and assembled using SPAdes with the parameters ‘--careful --sc --phred-offset 33’. From these assemblies, only contigs greater than 2200 bp were kept for the final assembly.

Eleven single-cell amplified genomes (ASAGs) affiliated to ANME clades from the Gulf of Mexico were identified. These had different levels of completeness and were clustered using the previously amplified 16S rRNA gene fragments and genome comparisons (average nucleotide identity, ANI) in to seven different groups for coassembly in order to improve completeness. Raw reads of the clustered were coassembled using SPAdes 3.9.0 (202) and scaffolds under 2000 bp were discarded from the coassembled genomes. Contamination and completeness was assessed by checkM (203). In total, 4 genome bins were retrieved: ANME GoMg1 (combined from 2 SAGs), ANME GoMg2 (combined from 2 SAGs), ANME GoMg3.2 (combined from 5 SAGs) and ANME GoMg4 (combined from 2 SAGs).

### Haakon Mosby Mud Volcano (HMMV) assembly and binning

The Haakon Mosby Mud Volcano is located on the Norwegian continental slope, west of the Barents Sea at 1,270 m water depth and it is characterized by gas hydrate accumulation in certain areas (204). ANME genomes were recovered from sediment samples collected on two different cruises, and from a laboratory enrichment culture inoculated from sediment collected on a third cruise. Genome ANME sp. HMMV and ANME HMMV2 were obtained from publicly available data in the sequence read archive (SRA) from a 2010 cruise to HMMV, accessions SRR1971623 and SRR1971624, respectively.

Genome ANME HMMV-459A3 was obtained from a sediment sample (459A) collected during cruise PS ARKII/1b in June/July 2007 from station PS70/096-1; 72.0032° N, 14.7205° E at 1263 m water depth; a site covered with a white microbial mat. Directly after recovery, samples were frozen at -20°C until further processing. Genomes of ANME sp. HMMV-459B4 and -459B1 were obtained from an enrichment culture established from sediment samples collected during cruise PS ARKXIX/3b in June/July 2003. Directly after recovery, samples were frozen at -20°C and then were used to established enrichment cultures in the Max Planck Institute of Marine Microbiology (Bremen, Germany). The enrichment cultures were established from sediment samples diluted with artificial seawater (205) with methane as the only energy source as described previously (186). Enrichment cultures were incubated at 4°C in the dark and propagated when sulfide concentration reached values over 12 mM. DNA was extracted from the sediment sample and an enrichment aliquot following the Zhou protocol (206) and sent to the Max Planck Genome Center in Cologne for sequencing. From the extracted DNA, Illumina paired-end libraries were prepared and sequenced using HiSeq machines generating 2x100 paired-end reads with an average insert length of 290 bp. In total, 26,574,291 and 28,965,807 paired-end reads were generated respectively for libraries 459A and 459B. The metagenomic reads from 459A and 459B libraries were independently assembled using SPAdes 3.9.0 (202). The assembled metagenome was binned using MetaWatt 3.5.3 (207) with a scaffold length threshold of 1000 bp.

#### ANME HMMV

SRR1971623 was downloaded from the NCBI sequencing read archive and assembled using Megahit (208). ANME sp. HMMV was binned from the metagenome using differential coverage as described previously (209), using read mapping of SRA runs SRR1971621, SRR1971623 and SRR1971624 with BBmap (http://sourceforge.net/projects/bbmap/) with a 98% identity cutoff. To further improve upon this bin, it was used for 30 iterative cycles of mapping and assembly, to extend the edges of contigs and fill gaps, using BBmap and SPAdes.

#### ANME HMMV2

SRR1971624 reads were recovered from SRA, trimmed, and quality filtered using Trimmomatic (210). The reads were assembled with Metaspades, version 3.9.0 using the default parameters (211). Scaffolding and gapfilling of the metagenome assembly was performed using the ‘roundup’ mode of FinishM v0.0.7 (https://github.com/wwood/finishm). Population genomes were recovered from the assembled contigs using the groopm2 (https://github.com/timbalam/GroopM) binning method (212).

### Amon Mud Volcano (AMV) assembly and binning

Genomes of ANME sp. AMVER4-31 and AMVER4-21 were obtained from sediment samples coming from Amon Mud Volcano (031° 42.6’ N, 032° 22.2’ E) during cruise R/V L’Atalante on September/October 2003 using the submersible Nautile. Amon Mud volcano is a gas-rich mud volcano located in the Eastern Mediterranean at 1,120 m water depth (213). Directly after recovery, samples were frozen at -20°C and then were used to established enrichment cultures in the Max Planck Institute of Marine Microbiology (Bremen, Germany). The enrichment cultures were established and maintained as described above for the HMMV enrichments with the exception of the incubation temperature that was 20 °C. DNA was extracted following the Zhou protocol (206). DNA was sent to the Center for Biotechnology (CeBiTec, Bielefeld, Germany) for sequencing. From the extracted DNA, two Illumina paired-end libraries were prepared and sequenced generating 2x250 paired-end reads with an average insert length of 550 bp. In total, 3,886,975 and 5,190,719 paired-end reads were generated respectively for each library. The metagenomics reads of both libraries were used for a combined assembly using SPAdes 3.9.0 (202). The resulting assembled metagenome was binned using MetaWatt 3.5.3 (207)with a scaffold length threshold of 1000 bp.

### ANME SA assembly and binning

Raw reads from MG-RAST sample mgm4536100.3, which were sourced from a South African Gold mine (214), were downloaded (https://metagenomics.anl.gov/) and assembled using megahit 1.0.3 (208). Contig coverage was estimated by mapping the reads onto the assembled contigs using BBmap 33.21. Subsequently, the coverage and composition of a contig containing the ANME-1 mcrA gene was used to guide manual binning of the ANME-1 genome using R statistical programming language (R Core Team 2016), as described previously (215).

This initial genome bin was improved by reassembly using spades 3.9. 0 (202). First, all reads from the samples were aligned against the initial genome bin with bwa 0.7.10 using the ‘mem’ algorithm and default alignment settings (216). Reads that aligned to the initial genome bin were separated into proper pairs - which also included reads that did not map to the genome bin but their paired sequence did - and reads without a pair in the dataset. These reads were used as input for spades 3.9.0, which was run with default settings. The scaffolds from the spades assembly were filtered to remove those that were less than 1000bp or those with a coverage of less than 5 or greater than or equal to 30; both of these statistics were taken straight from the contig names produced by spades. Additionally, five contigs were manually removed as they contained partial 16S rRNA genes that were not phylogenetically related to ANME-1, in addition to one large contig which appeared to be a phage genome.

### ANME S7142MS1 assembly and binning

DNA from methane seep sediment incubation #7142 (∼2mL) was extracted using the MoBio Powersoil DNA kit (MoBio) according to the manufacturer’s protocol. DNA concentrations were quantified using the Quant-iT dsDNA HS assay kit (Invitrogen) as per the manufacturer’s instructions. The paired-end library was prepared using the Nextera XT DNA library preparation kit (Illumina, USA) for DNA extracted. The libraries were sequenced on a NextSeq500 (Illumina, USA) platform generating 2×150 bp paired-end reads with an average insert length of 300 bp. In total, 168,382,029 paired-end reads for the metagenome were generated from the library. Bulk metagenome reads were trimmed and quality filtered using Trimmomatic (210) and BBMerge (http://sourceforge.net/projects/bbmap/) using default settings. Low-abundance k-mer trimming was applied using the khmer (217) script trim-low-abund.py using with the K = 20 and C = 30 parameter and assembled with MetaSPAdes (211), version 3.9.0 using the default parameters. Scaffolding and gapfilling of the metagenome assembly was performed using the ‘roundup’ mode of FinishM v0.0.7 (https://github.com/wwood/finishm). Population genomes were recovered from the assembled contigs using MetaBat (218). ANME sp. S7142MS1 was further refined by removing scaffolds with divergent GC-content, tetranucleotide frequencies or coverage using the outlier method in RefineM v0.0.13 (https://github.com/dparks1134/RefineM).

### ANME SCGC AAA252-L18 assembly and binning

ANME SCGC AAA252-L18 was coassembled from 12 single cell datasets publicly available under the NCBI bioproject accession PRJEB7694, which were sourced from a mud volcano in the Gulf of Cadiz. Reads from these 12 samples were assembled together using Spades 3.5.0 (202) using the single cell (--sc) mode. This combined assembly was binned using VizBin (199) to remove contigs that contained divergent pentanucleotide sequences.

### ANME ERB4, ERB6, ERB7 binning

Fosmid libraries from Eel River Basin were isolated and sequenced in conjunction with a previous study (6). Previously unpublished fosmids were binned using VizBin (199) to create ANME ERB4, ERB6 and ERB7. These fosmids-derived genome bins were quite large due to duplicate regions of the genomes and represent a population of very closely related strains which were difficult to further assemble due to apparent genome rearrangements. Fosmid NCBI accession numbers assigned to specific bins can be found in **S5 Data**.

### ANME-1 LC assembly and binning

A metagenome coassembly from various Lost City hydrothermal fluids (unpublished) was screened for McrA sequences from the ANME-1 SA group of ANME-1 since McrA from this group has been previously reported at this site (179). A close hit ∼85% identity to ANME-1 sp. SA McrA was found on a short contig. Reads from all separate samples were mapped onto this contig and one sample was found to have high coverage of this contig. This sample was reassembled alone using MEGAHIT v1.2.9 (208) using minimum contig size of 1000 and meta-large option. Assembly was binned with MetaBAT2 v1.7 and genome completeness and phylogeny were determined by CheckM v1.0.18 and GTDB_Tk v1.1.0, respectively (203, 219). One bin was highly >80% complete and identified as being ANME-1 by GTDB_Tk. The entire assembly was visualized in R using scripts from mmgenome package and a subset of ∼5000 contigs surrounding those assigned to this bin in a plot of coverage vs. the second principle component of tetranucleotide frequency were exported (220). These 5,000 contigs were subjected to manual binning using the anvi’o workflow for metagenome binning (with only a single sample for coverage information) (221). This improved genome bin was 88% complete and 2% contaminated as estimated by CheckM. This ANME-1 LC MAG was 1,073,849 bp in length and contained 152 contigs. MEGAHIT, CheckM, GTDB_Tk were implemented on the KBase platform (222). The ANME-1 LC assembly has been uploaded to NCBI under Assembly accession number GCA_014061035.1.

### Quality and taxonomic assessment of genomes

All genomes were assessed for their completeness and contamination using checkM 1.0.6 (203) using the ‘taxonomy_wf’ command and the Euryarchaeota marker set containing 188 marker genes. Genome taxonomy was assigned using the GTDB_Tk (219) and naming conventions are consistent with the GTDB release 89.

### 16S rRNA gene sequencing of sorted single aggregates

For sorted single aggregate samples that did not have a 16S rRNA gene in the binned genomic dataset, we attempted to amplify the 16S rRNA genes for cloning and subsequent sequencing. The original MDA amplified samples were used as template to amplify the bacterial 16S rRNA gene using a modified universal reverse primer u1492R (5’-GGYTACCTTGTTACGACTT-3’) and modified bacterial b8F-ym (5’-AGAGTTTGATYMTGGCTC-3’) (223). The u1492R primer was also paired with a modified archaeal a8F-y (5’-TCCGGTTGATCCYGCC-3’) to amplify the archaeal 16S rRNA gene (224). Fifteen microliter PCRs were run for 1 min at 94°C, followed by 40 cycles of 15s at 94°C, 30s at 54°C, and 45s at 72°C, and a final elongation step at 72°C for 4 min. The reaction mix included 0.2 μM forward and reverse primers (Integrated DNA Technologies, Inc. Coralville, IA, USA), 1× ExTaq PCR buffer (Takara, Clontech Laboratories, Inc., Mountain View, CA, USA), 0.75 U ExTaq (Takara), 0.22 mM dNTPs (New England Biolabs, Ipswich, MA, USA), and 1 μl template. No amplicon was visible in triplicate DNA control reactions when 4 μl were run on a 1.5% agarose gel visualized with SYBR Safe (ThermoFisher Scientific; cat. no. S33102). PCR products were plate purified (Millipore Multiscreen filter plates; cat. no. MSNU03010), ligated with the Invitrogen TOPO TA Cloning Kit (ThermoFisher Scientific; cat. No. K457501) and transformed using Top Ten chemically competent Escherichia coli cells. Picked colonies were grown overnight in Luria–Bertani broth (Miller’s modification) and amplified using M13 primers for 30 cycles. The M13 products were visualized to confirm the correct size insert and screened by RFLP digest using HaeIII enzyme (NEB) on a 3% agarose gel. All unique clones were plate purified and sent for sequencing using T3 and T7 primers at Laragen Sequencing (Culver City, CA, USA). Contigs were assembled using Geneious software version R10 (225). Sequence accessions for the clones from these genomes can be found in **S2 Table**.

### Calculation of genome similarity

The average amino acid identity (AAI) was calculated between each pair of genomes was calculated using compareM and its AAI workflow (aai_wf) command (https://github.com/dparks1134/CompareM). Briefly this workflow first identifies all open reading frames were determined using prodigal 2.0.6 (226) and then homologous genes were determined using reciprocal best-hits using 0.7.11 (227). The AAI was calculated by averaging the percent similarity of all proteins that were reciprocal best hits. The genome similarity (GS) displayed in **S1 Fig** was calculated by first calculating the percent alignment between the genomes as the fraction of the genes that are reciprocal best hits (RBH), using the smaller number of total genes between the two genomes (X & Y) as the denominator, and then multiplying the AAI value by that fraction (i.e. GS = AAI*RBH/min(X, Y)). The order of the matrix was calculated using the R statistical programming language 3.3.2 (R Core Team 2016) using the ‘hclust’ function using the default distance metric (‘euclidean’) and the ‘ward.D2’ method of agglomeration. This matrix was visualized using ggplot2 (228).

### 16S rRNA gene phylogeny

16S rRNA gene sequences were extracted from SILVA 128 (229, 230) and new sequences from ANME genomes were aligned using SINA 1.2.11 (231) though the online portal at https://www.arb-silva.de/aligner/. Alignment columns were masked in ARB 6.0.2 (232) using the inbuilt ‘archaea ssuref’ filter and exported in fasta format removing all columns that contained only gaps. The phylogenetic tree in **Fig 1A** was constructed using RAxML 8.1.7 (233) using the following parameters: raxmlHPC-PTHREADS-SSE3 -f a -k -x 12395 -p 48573 -N 100 -T 16 -m GTRGAMMAI. Trees were visualized using ete3 (234).

### Concatenated protein phylogeny

The genome trees in **Figs 1B, 3** and **18** were constructed from a list of 43 hidden markov models (HMM) trained from proteins common to both bacteria and archaea sourced from Pfam and TIGRfam (see above for list). Open reading frames were annotated onto genome sequences using prodigal 2.6 (226) and candidates for the marker genes were identified using hmmsearch 3.1b1 (http://hmmer.org). Only the top match to each maker was retained and aligned to the HMM using hmmalign 3.1b1 (http://hmmer.org). The alignment was trimmed on each side using the stockholm format reference coordinate annotation (line beginning with #=GC RF) to include only residues that matched the reference (columns containing an ‘x’). The protein sequences were then concatenated and a phylogenetic tree was constructed using RAxML 8.1.7 (233) using the following parameters: raxmlHPC-PTHREADS-SSE3 -f a -k -x 67842 -p 568392 -N 100 -T 16 -m PROTGAMMAWAG -o IMG_638154518. Note that the outgroup is specified in the command line (-o IMG_638154518), which corresponds to Sulfolobus solfataricus.

### McrA phylogeny

McrA sequences from cultured methanogenic and methanotrophic archaea and genome sequences from uncultured representatives were downloaded from the NCBI refseq database. These sequences were combined with clone sequences from ANME dominated environments (31, 235) and from the ANME genomes sequenced here. These sequences were aligned using muscle 3.8.31 (236) with the default parameters. The phylogenetic tree in **Fig 1D** was constructed using RAxML 8.1.7 (233) using in the following parameters: raxmlHPC-PTHREADS-SSE3 -f a -k -x 67842 -p 19881103 -N 100 -T 16 -m PROTGAMMAWAG.

### Transcriptome data

Data shown in **S3 Data** was reproduced from the main text or supplemental data of papers reporting on the ANME-2a Wang genome (11), the ANME-2d BLZ1 genome (19), and the ANME-2c E20, ANME-1 GB37 and GB60 genomes (9).

### NifD and FpoH phylogenies

Homologs of NifD and FpoH were identified in ANME genomes using KEGG ortholog IDs K02586 and K00337, respectively. Genes annotated with these KEGG IDs were separately combined with a set of protein sequences from uniprot that were also annotated with these IDs using the following website (in the case of NifD (http://www.uniprot.org/uniprot/? query=k02586&sort=score&format=fasta). A representative set of genes was made using cd-hit 4.6 (237) by removing all sequences that were greater than 95% amino acid similarity to each other. In the case of FpoH most bacterial sequences were omitted. These sequences were aligned using muscle 3.8.31 (236) with the default parameters. A phylogenetic tree was constructed using RAxML 8.1.7 (233) using in the following parameters: raxmlHPC-PTHREADS-SSE3 -f a -k -x 8512 -p 110339 -N 100 -T 16 -m PROTGAMMAWAG. Tree was visualized using ete3 (234).

### Additional protein phylogenies

Phylogenetic trees for RpoB, MtrE, Mer, Mtd, Mch, Ftr, FwdB/FmdB, MetF, HdrA, VhtC/McoA, Small ANME MHCs, CcmF, GlyA, ThyA, FrhB were constructed based on alignments using muscle 3.8.31 (236) with the default parameters. Trees were built with PhyML 3.0 (238) implemented on the website (http://www.atgc-montpellier.fr/phyml/) with automatic substitution model selection based on the Akaike Information Criterion as implemented by SMS (239). Branch support was provided by the aLRT SH-like fast likelihood-based method. Trees were visualized with iTOL (240).

### Determination of cytochrome orthologs

Homologous proteins displayed in **S13 Fig** were determined using proteinortho 5.16 (241) using the option ‘-singles’ to keep even unique proteins. Orthologs containing multiheme cytochromes were determined by counting the number of CxxCH motifs in the amino acid sequences. Orthologous groups where the median count of heme binding domains was greater than one were included for analysis. These groups were then curated to resolve any erroneous placement of individual proteins. Each group of cytochromes was aligned using muscle 3.8.31 (236) and individual sequences that did not share conserved residues or contained unusual insertions or deletions were moved into separate orthologs. Proteins that were originally categorized as unique were also compared using blastp 2.6.0+ (242) to determine if they could be placed into an orthologous group.

### Analysis of ANME-MHCs

Cytochromes with greater than ten heme binding motifs were annotated using Interpro (243, 244) and were categorized based on the presence of an S-layer domain, transmembrane helixes, and the presences and type of putative sugar binding/degrading motifs.

### Identification of dockerin/cohesin domains

Proteins containing dockerin and/or cohesin domains were identified by predicting domains in all hypothetical proteins in all ANME genomes using the KBase platform’s Annotate Domains app (v.1.0.7) with the “All domain libraries” function (drawing from COGs, CDD, SMART, PRK, Pfam, TIGRFAMs and NCBIfam) (222). All domain annotations were downloaded as csv files, and any protein containing a dockerin or cohesin domain was exported along with other domains therein to a separate csv file. These csv files can be found in **S4 Data**.

## Acknowledgments

We thank Stephanie A. Connon and Alice Michel for help sequencing 16S rRNA genes from some single aggregates. The work conducted by the U.S. Department of Energy Joint Genome Institute, a DOE Office of Science User Facility, is supported under Contract No. DE-AC02-05CH11231. We acknowledge the support of all crews of the expeditions mentioned here for sampling and work at sea, and the work of Susanne Menger for the cultivation of AOM consortia. Funding for this study was received by the DFG Leibniz grant and by the Max Planck Society to AB. Genomes of BONCAT-FACS sorted ANME-consortia were generated via a JGI Director Discretionary Project Award (to R.H. and V.J.O.). We thank Chief Scientist Susan Lang, the scientific party of the 2018 Lost City Expedition, and NSF support to Lang and Brazelton (OCE-1536702/1536405). Grayson L. Chadwick is supported by the Miller Institute for Basic Research in Science, University of California Berkeley.

The authors declare no competing interests.

---

**S1 Fig.**
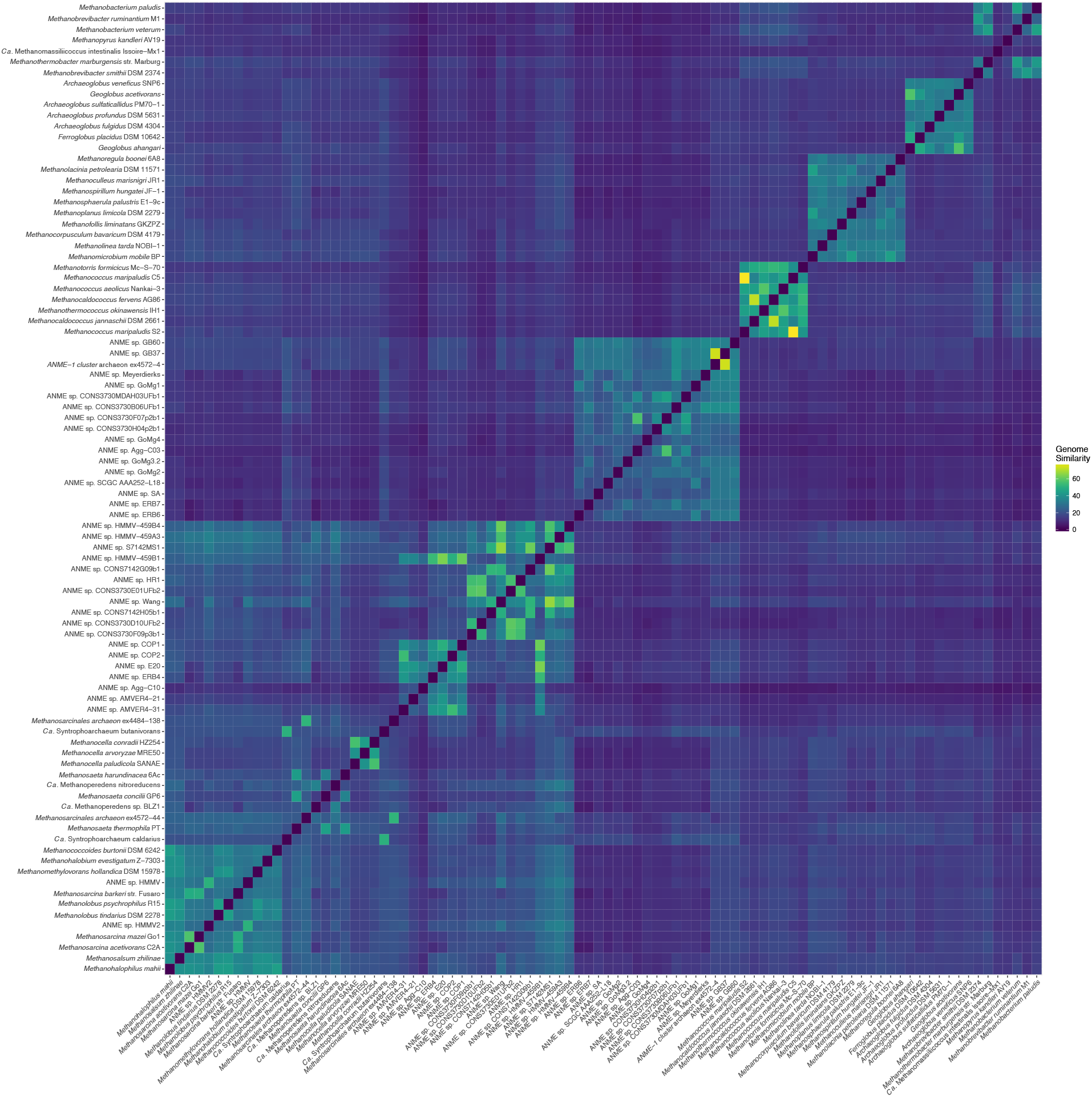
Genome similarity. Genome similarity between methanogen and ANME genomes reported here. Details of similarity calculations can be found below in the Materials and Methods.

**S2 Fig.**
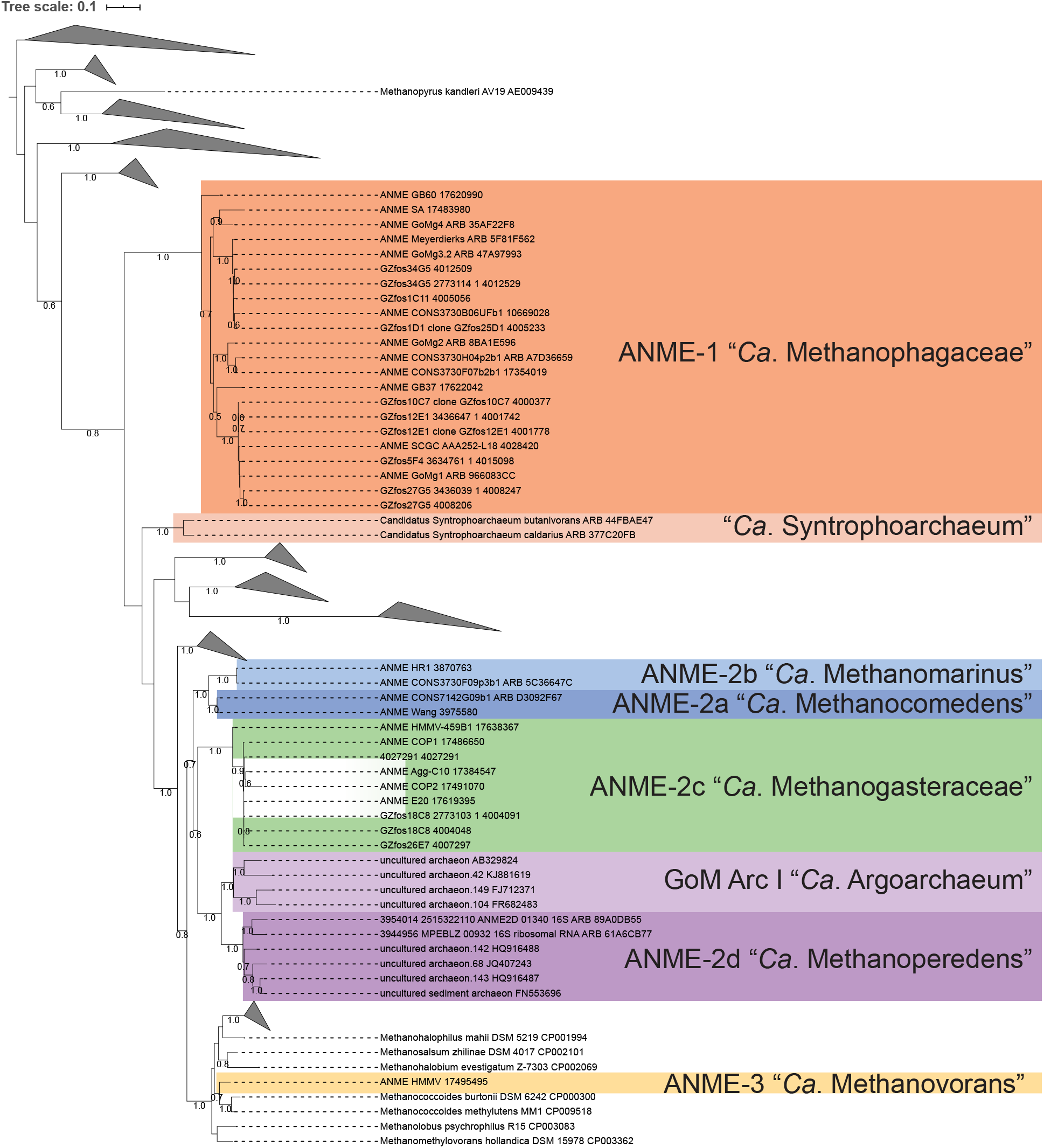
Expanded 16S rRNA gene tree. Tree expanding compressed clades in **Fig 1A.**

**S3 Fig.**
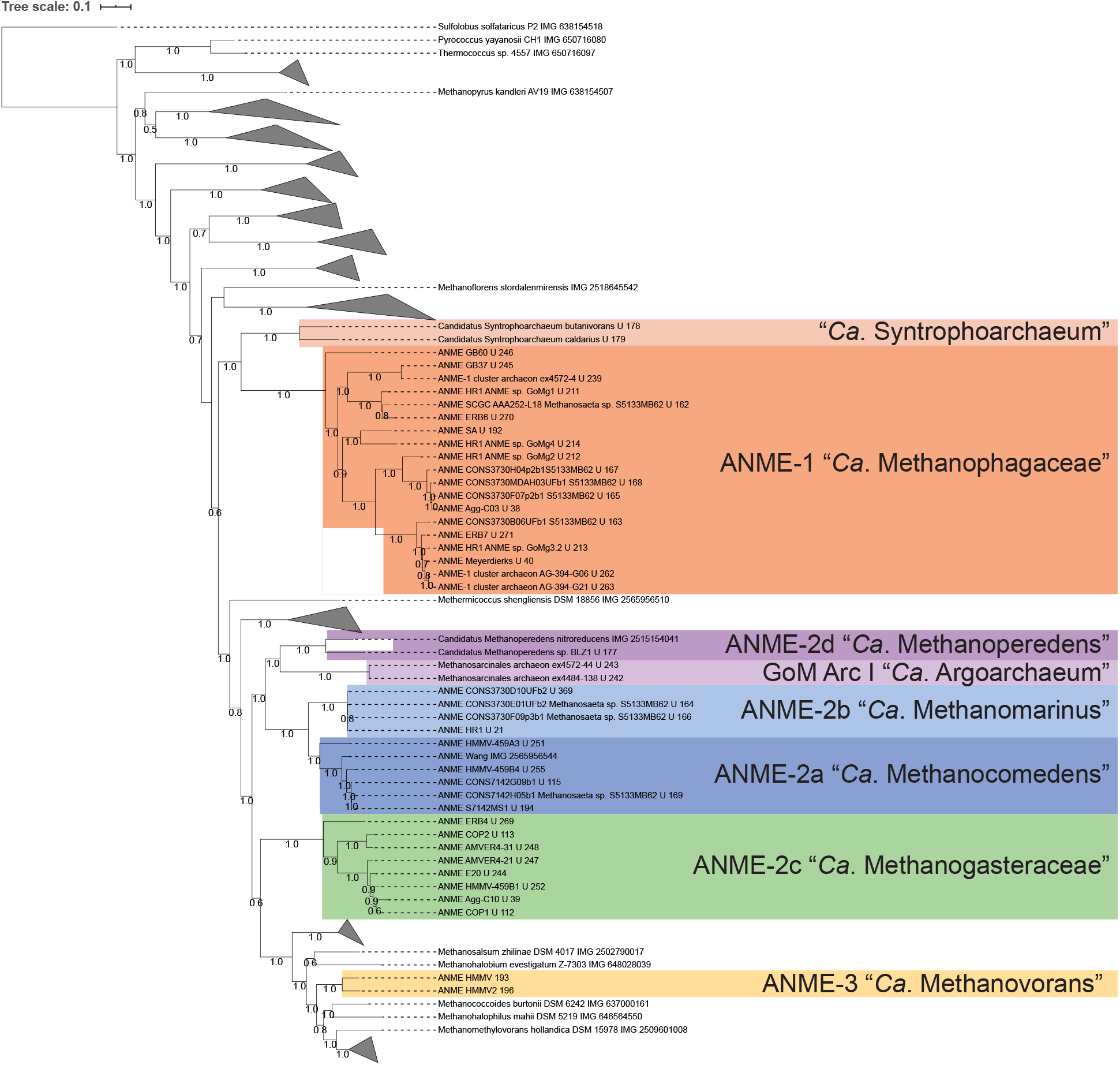
Expanded concatenated marker protein tree. Tree expanding compressed clades in **Fig 1B.**

**S4 Fig.**
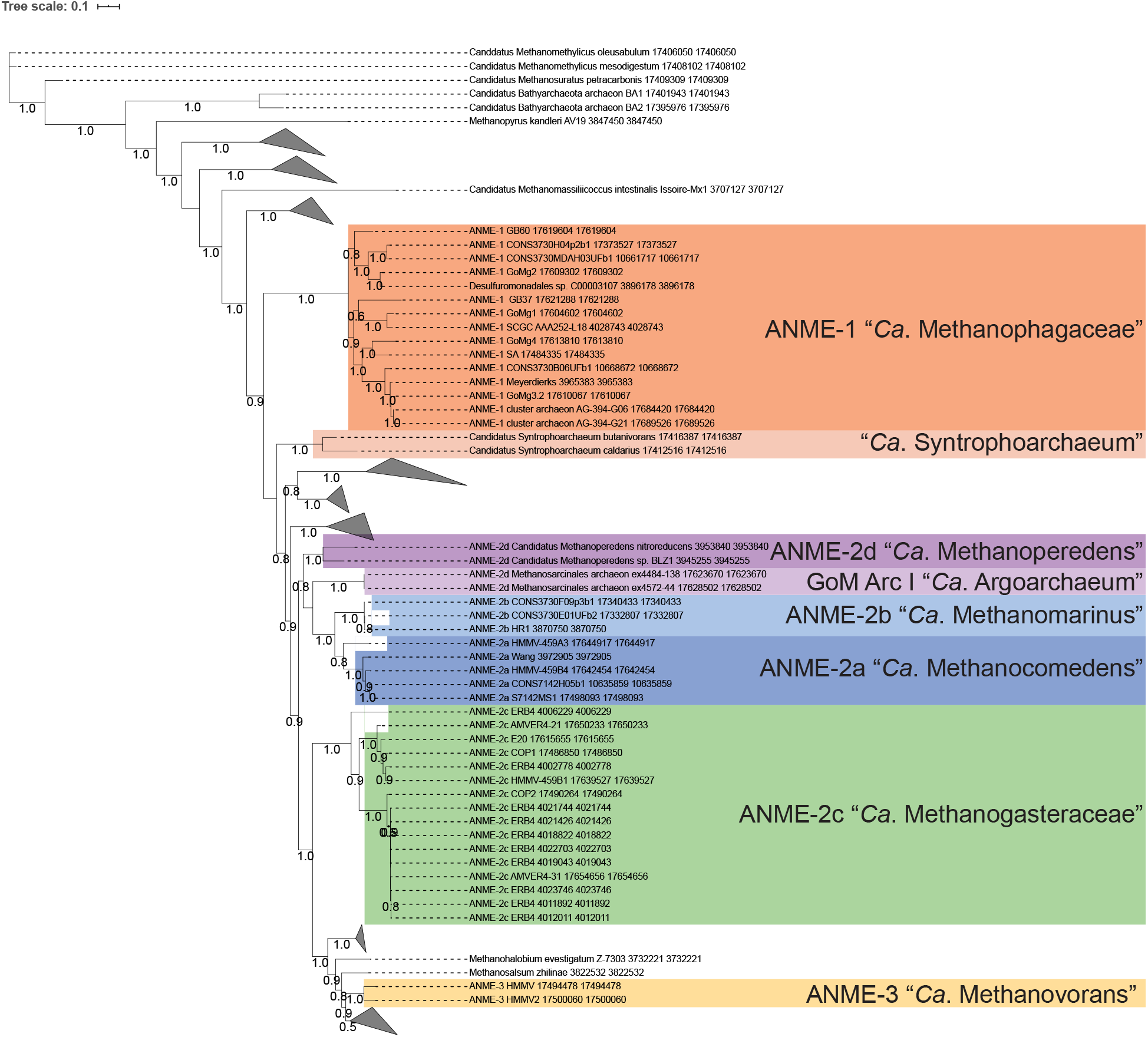
Expanded RpoB protein tree. Tree expanding compressed clades in **Fig 1C.**

**S5 Fig.**
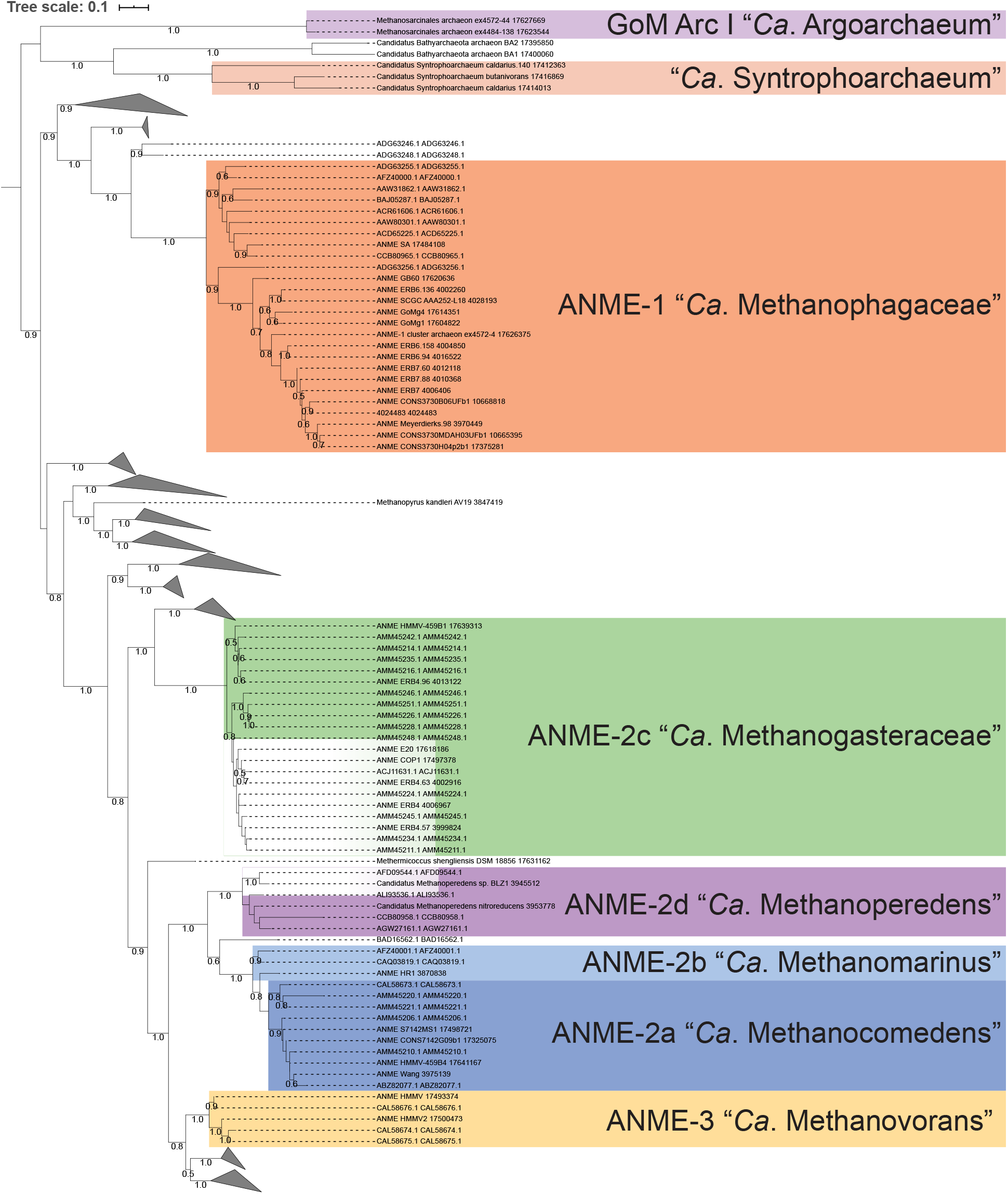
Expanded mcrA protein tree. Tree expanding compressed clades in **Fig 1D.**

**S6 Fig.**
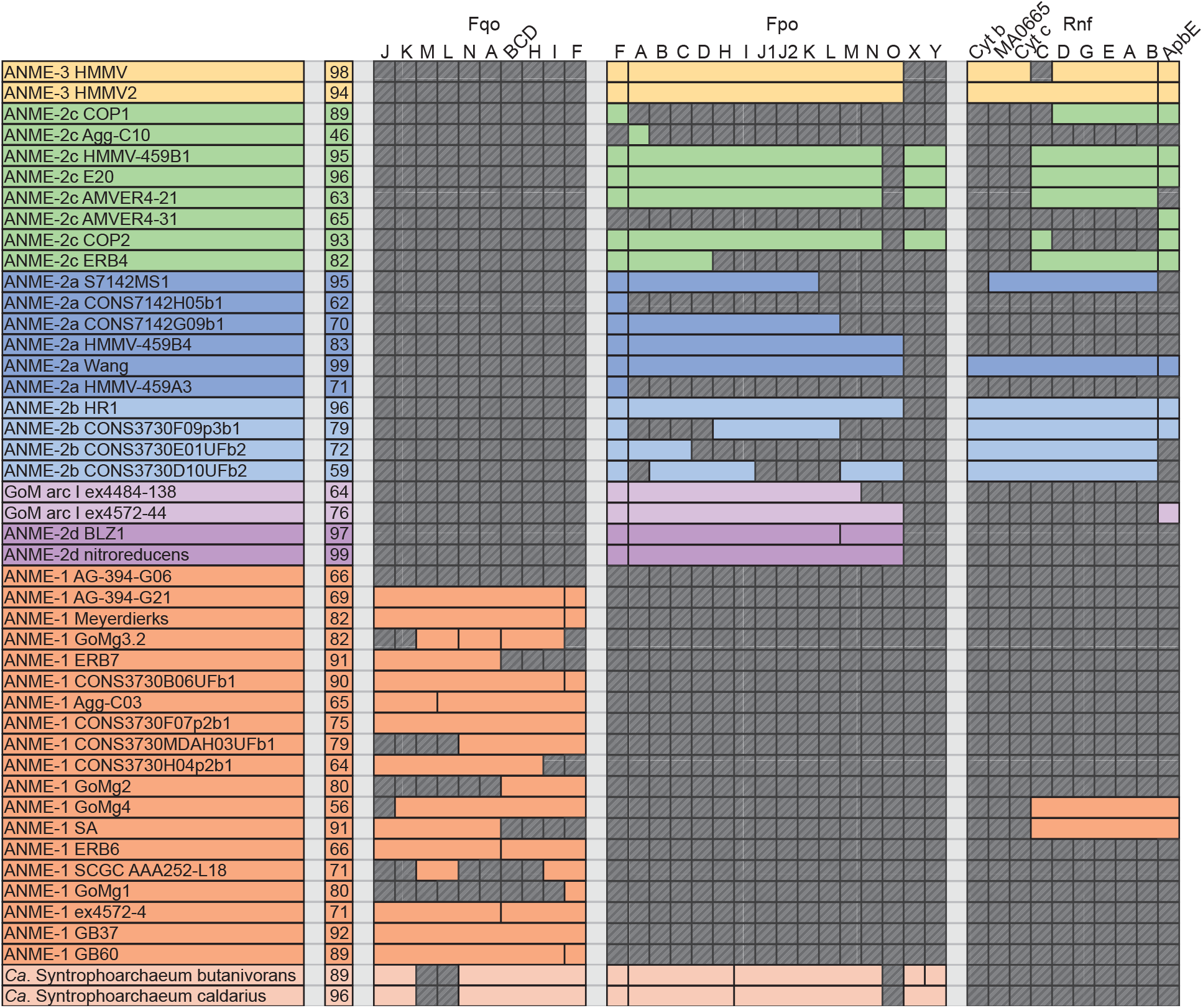
Presence of Fd^2-^ and F420H2 oxidation systems. The presence of each subunit of the oxidation complexes outlined in **Figure 6** is represented by colored boxes. ANME-2c and “*Ca.* Syntrophoarchaeum” lack the FpoO subunit found in other ANME and methanogens, and instead are followed by two highly expressed and conserved proteins denoted FpoX and Y. ApbE is a flavin transferase involved in Rnf biosynthesis. Missing genes are represented by transparent gray boxes with diagonal line fill. Numbers in the second column represent genome completeness. When genes are together in a gene cluster their boxes are displayed fused together. Gene accession numbers can be found in **S2 Data**.

**S7 Fig.**
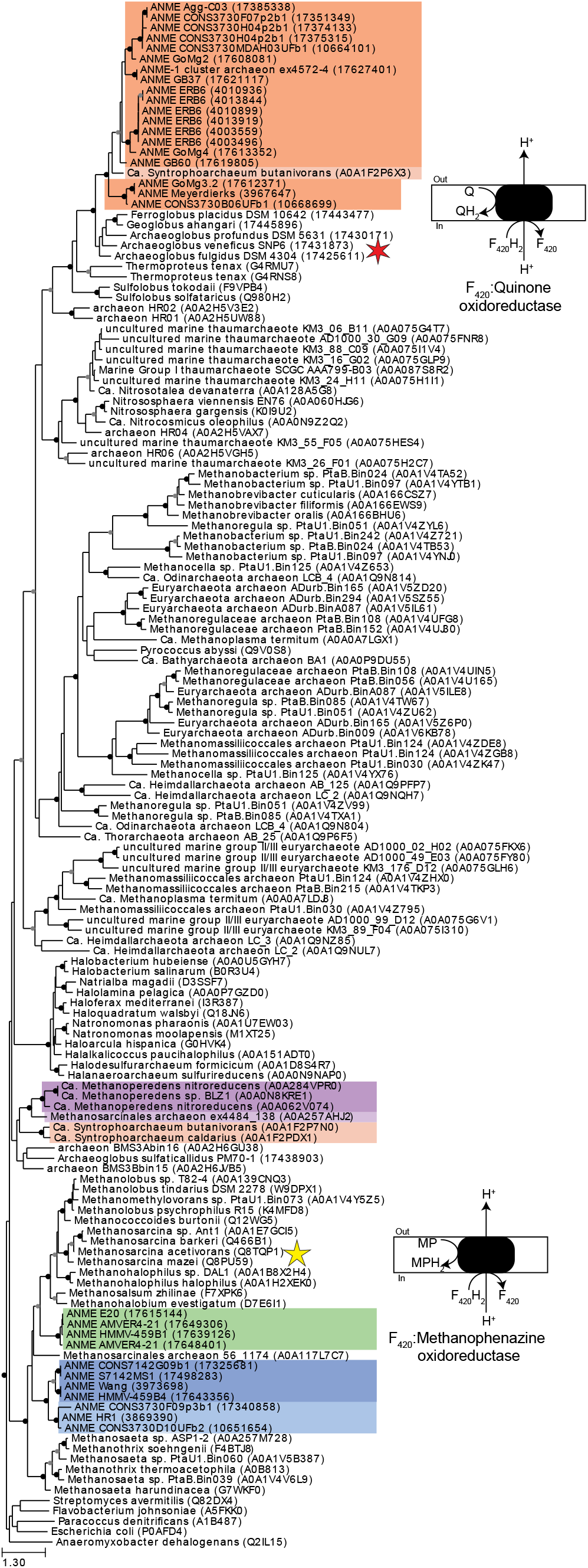
FpoH phylogenetic tree. Phylogeny of the H subunit of Fpo/Fqo complexes found in ANME and related organisms demonstrates the large evolutionary distance between these complexes. See methods for details of tree construction. Sequences labeled with five and six-sided stars represent the biochemically characterized Fpo and Fqo complexes, respectively. ANME groups are highlighted. Closed circles represent branch support values of 80 to 100%, gray circles between 70 and 80%. Tree scales represent substitutions per site. Tree construction parameters are found in the methods section. Alignment and tree file can be found in **S1 Data**.

**S8 Fig.**
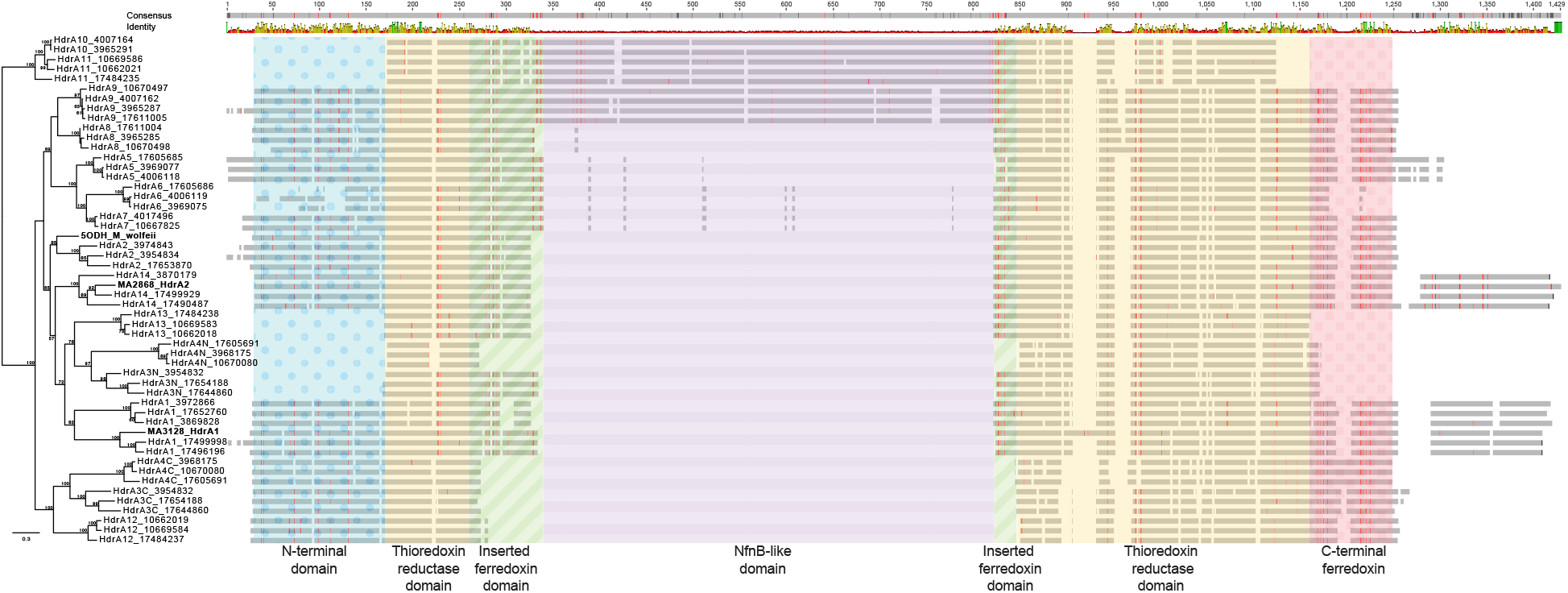
Alignment of HdrA sequences. Protein alignments of representative ANME HdrA sequences were made and overlaid with the domains as colored in **Figs 7 and 8** based on the domains identified in the *M. wolfeii* structure. Cysteine residues are highlighted as red lines. Large insertion in the middle corresponds to the NfnB-like domain that interrupts the inserted ferredoxin domain in some ANME-1 HdrA. “HdrA1 (MA3128)” and “HdrA2 (MA2868)” from *acetivorans* (numbering from Buan and Metcalf, 2010) are included as references. Branch support values >50% are reported. Tree scales represent substitutions per site. Tree construction parameters are found in the methods section. Alignment and tree files can be found in **S1 Data.**

**S9 Fig.**
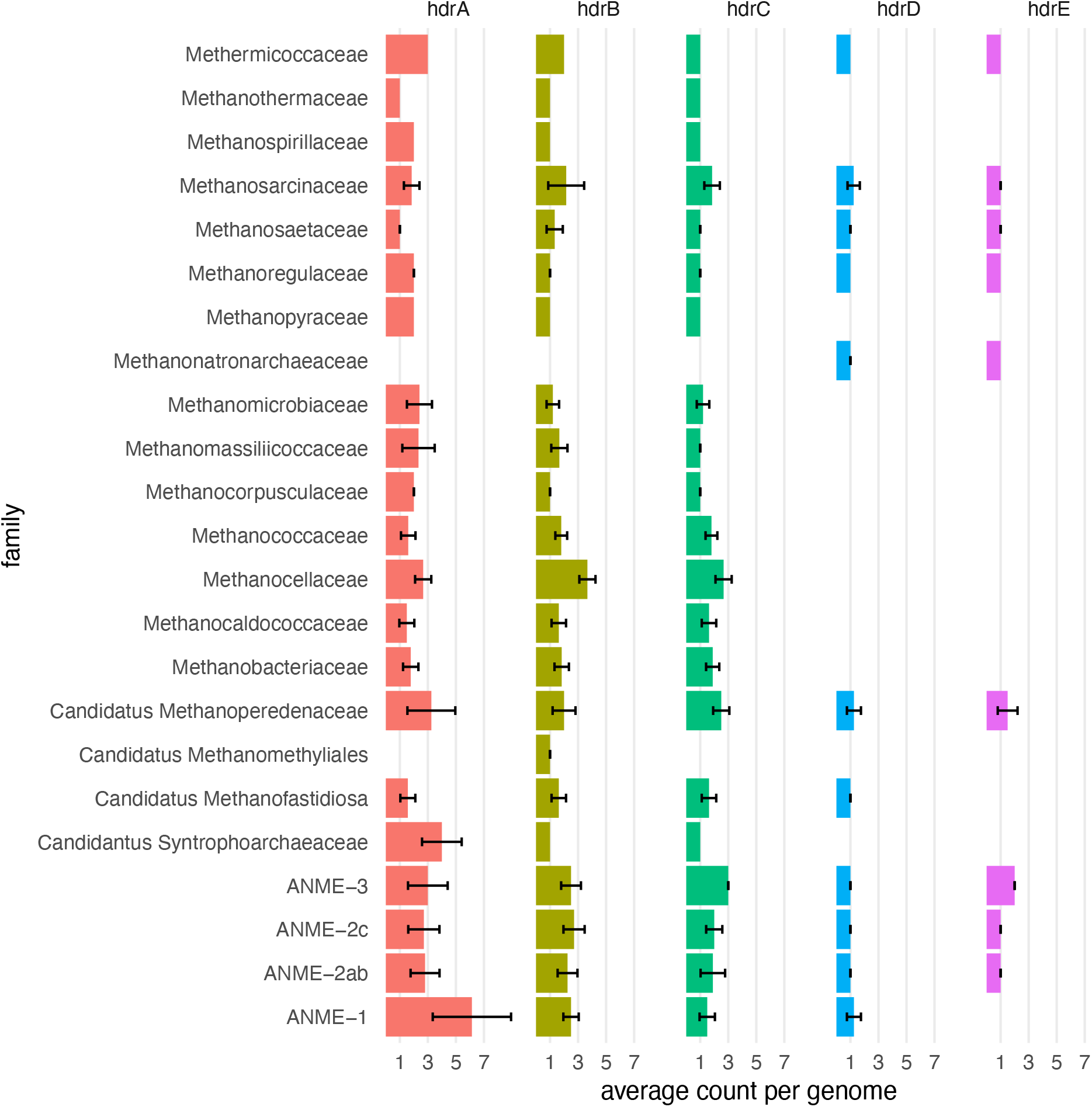
Hdr counts per genome. Most ANME and methanogen genomes contain 1-2 copies of *hdrABC* genes, however ANME-1 have a much greater abundance of *hdrA* homologs that are not accompanied by an increase in the number of *hdrB* or *C* homologs.

**S10 Fig.**
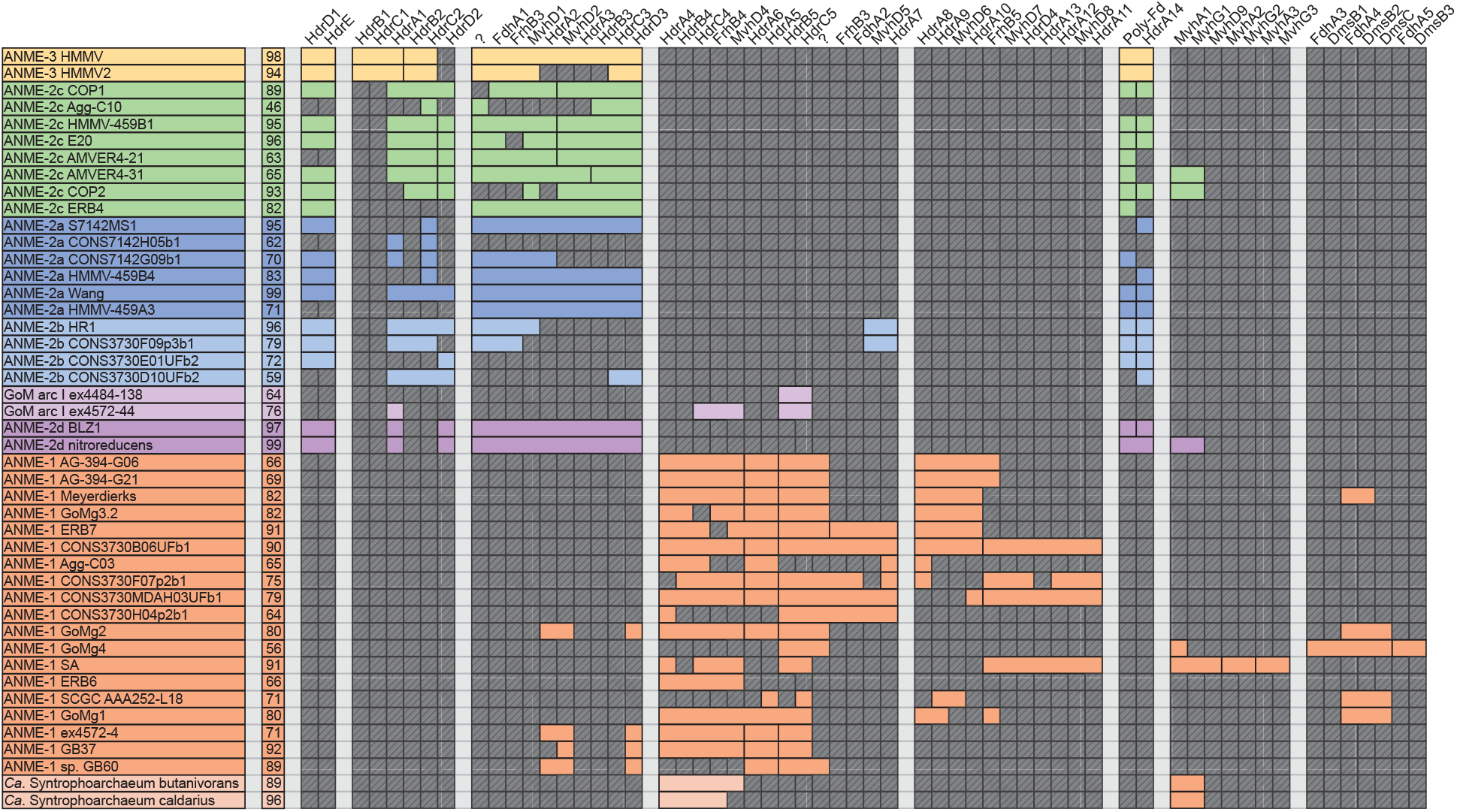
Hdr Presence/Absence. Colored boxes represent presence of Hdr and associated genes in ANME genomes. HdrA1 in ANME is the same group as HdrA1 (MA3128) from *M. acetivorans*. Interestinly, only in ANME-3 is HdrA1 paired with HdrB1C1 (numbering from Buan and Metcalf, 2010). In other ANME HdrA1 is paired with HdrB2C2. HdrA2 from *M. acetivorans* (here called HdrA14), is found in ANME-2 and 3 and largely lacks clear gene clustering, although its associated polyferredoxin is found elsewhere in most genomes. Question marks represent genes with no known annotation but that are conserved in Hdr gene clusters. Missing genes are represented by gray boxes with diagonal line fill. Numbers in the second column represent genome completeness. When genes are together in a gene cluster their boxes are displayed fused together. Numbers were assigned to distinct clusters of paralogs, matching those labels found in **Fig 8**. Gene accession numbers can be found in **S2 Data**.

**S11 Fig.**
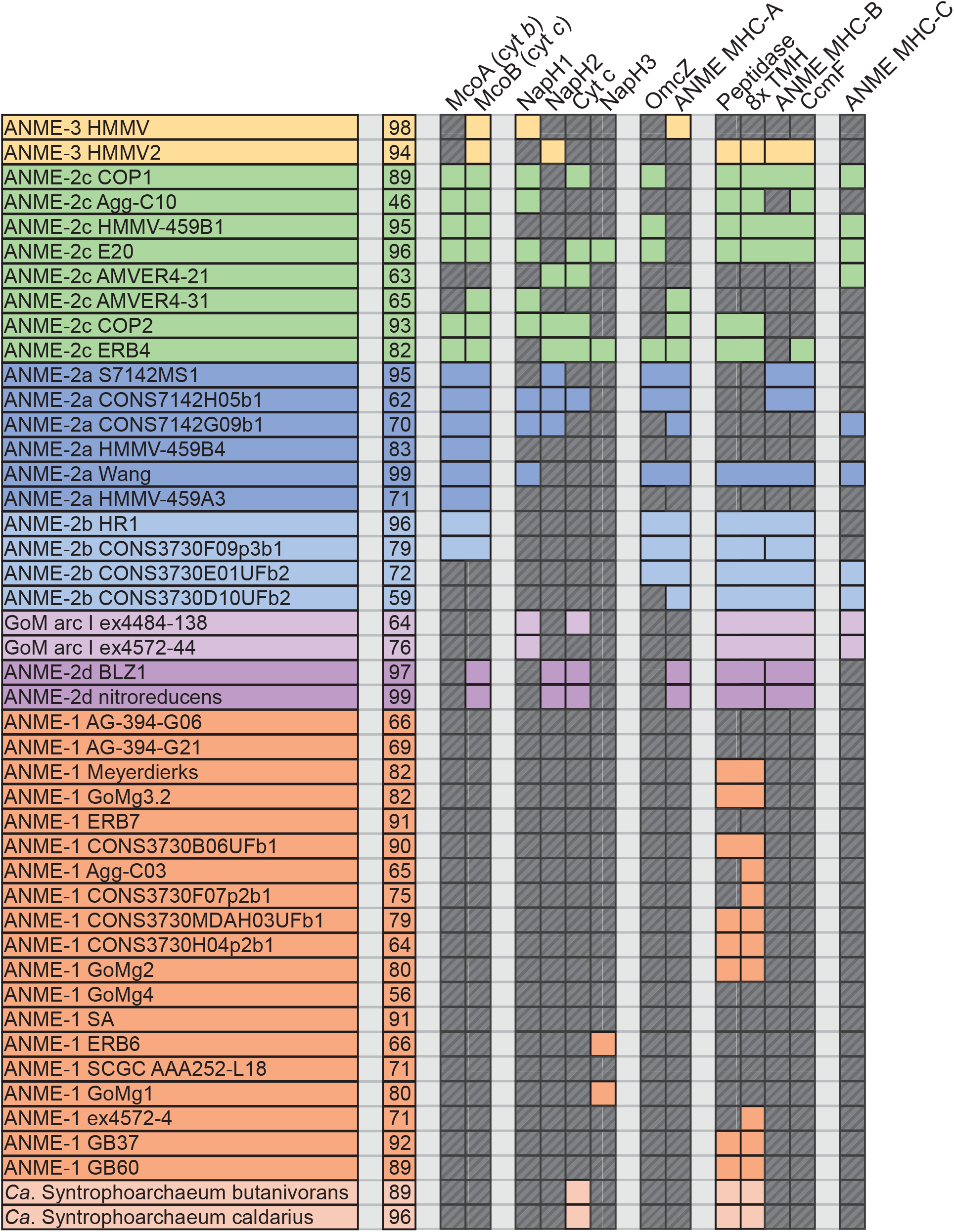
EET presence/absence. Colored boxes represent presence of various key hypothetical EET genes in ANME genomes. Peptidase and 8x TMH denote a signal peptidase involved in cytochrome maturation and a hypothetical membrane protein with 8 predicted transmembrane helices that are found only in ANME and EET-capable *Ferroglobus* and *Geoglobus sp*. Missing genes are represented by gray boxes with diagonal line fill. Numbers in the second column represent genome completeness. When genes are together in a gene cluster their boxes are displayed fused together. Gene accession numbers can be found in **S2 Data**.

**S12 Fig.**
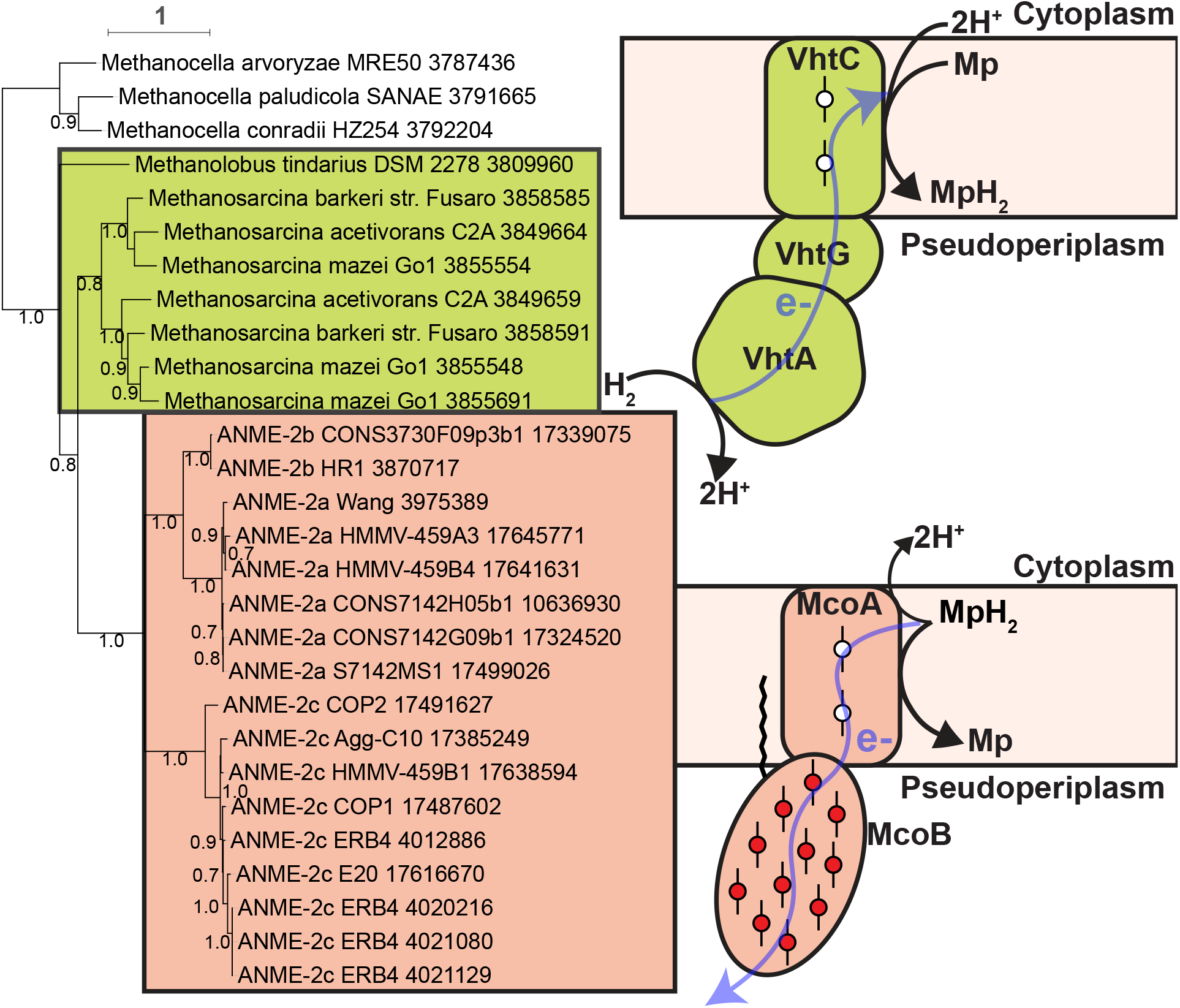
Mco phylogeny. VhtC and McoA homologs from a selection of methanogens and the ANME genomes reported here were used to build a phylogenetic tree. ANME VhtC homologs clustered with multiheme cytochrome genes in ANME-2a and 2b, whereas in methanogens they always clustered with the VhtAG gene encoding the hydrogenase enzyme used by these organisms. Branch support values >50% are reported. Tree scales represent substitutions per site. Tree construction parameters are found in the methods section. Alignment and tree files can be found in **S1 Data**.

**S13 Fig.**
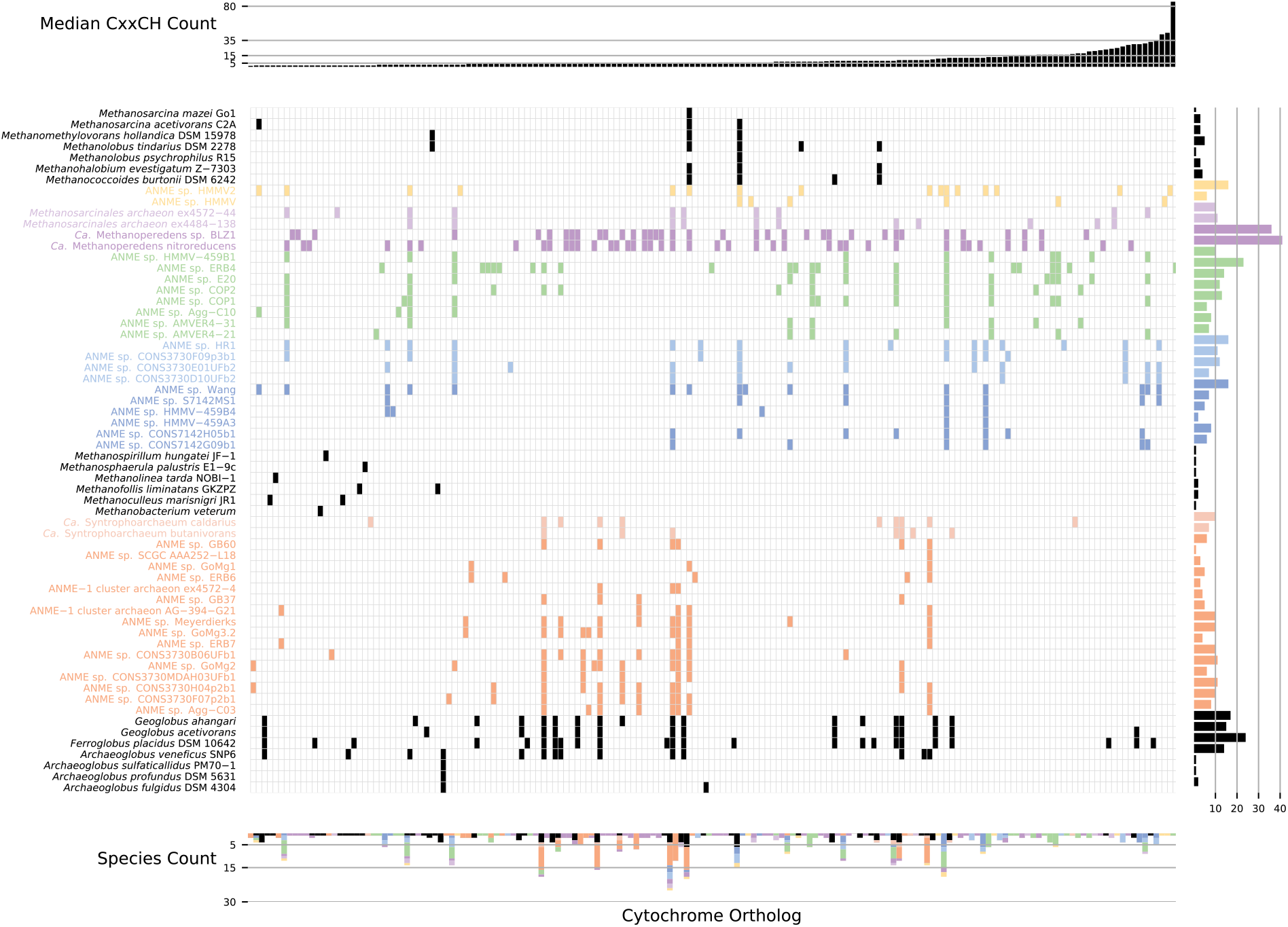
Cytochrome *c* orthologs. Multiheme cytochromes present in ANME, methanogens and *Archeoglobales* genomes. The central panel depicts the presence of cytochromes between genomes, where cytochromes between genomes have been grouped based on reciprocal best-hit blast results. The cytochromes in the central panel are ordered based on the median number of heme binding sites (CxxCH motifs) present in each group (upper bar plot). The clade distribution of each cytochrome is shown in the lower bar plot, while the right bar plot depicts the number of cytochromes present in each organism. Genomes that did not contain any multiheme cytochromes, which were most methanogens, are not shown.

**S14 Fig.**
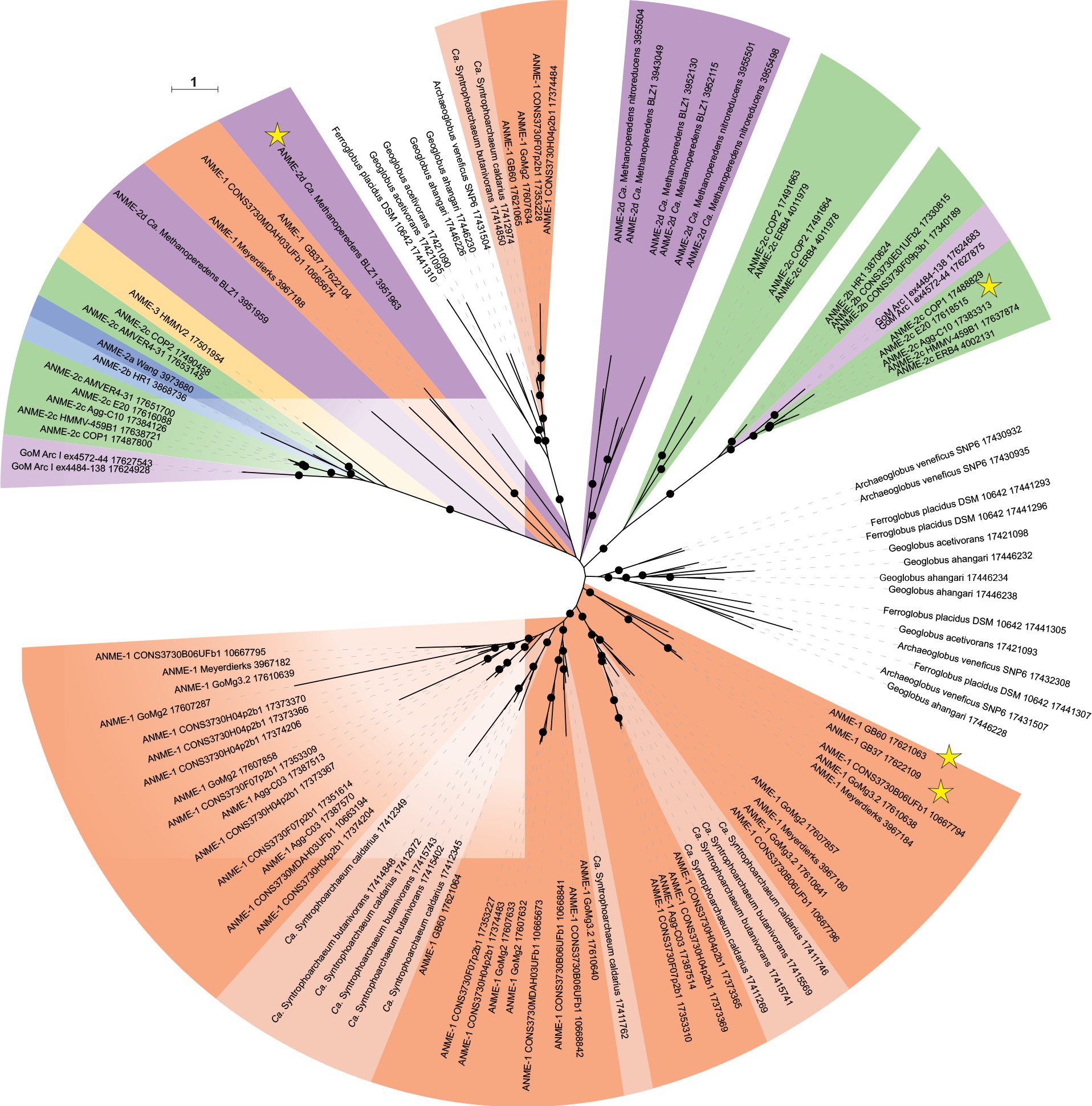
Broadly distributed cytochrome *c* homologs in ANME. Unrooted phylogenetic reconstruction of the most broadly distributed group of MHC proteins found in the ANME archaea. Homologs to this group were only found in ANME, “*Ca.* Argoarchaeum”, “*Ca*. Syntrophoarchaeum” and members of the *Archaeoglobales* known for their EET capabilities. Stars represents the sequence corresponding to the exceptionally highly expressed cytochromes highlighted in **S3 Data**. Branch support values >90% are depicted as filled circles. Tree scales represent substitutions per site. Tree construction parameters are found in the Materials and Methods section. Alignment and tree files can be found in **S1 Data**.

**S15 Fig.**
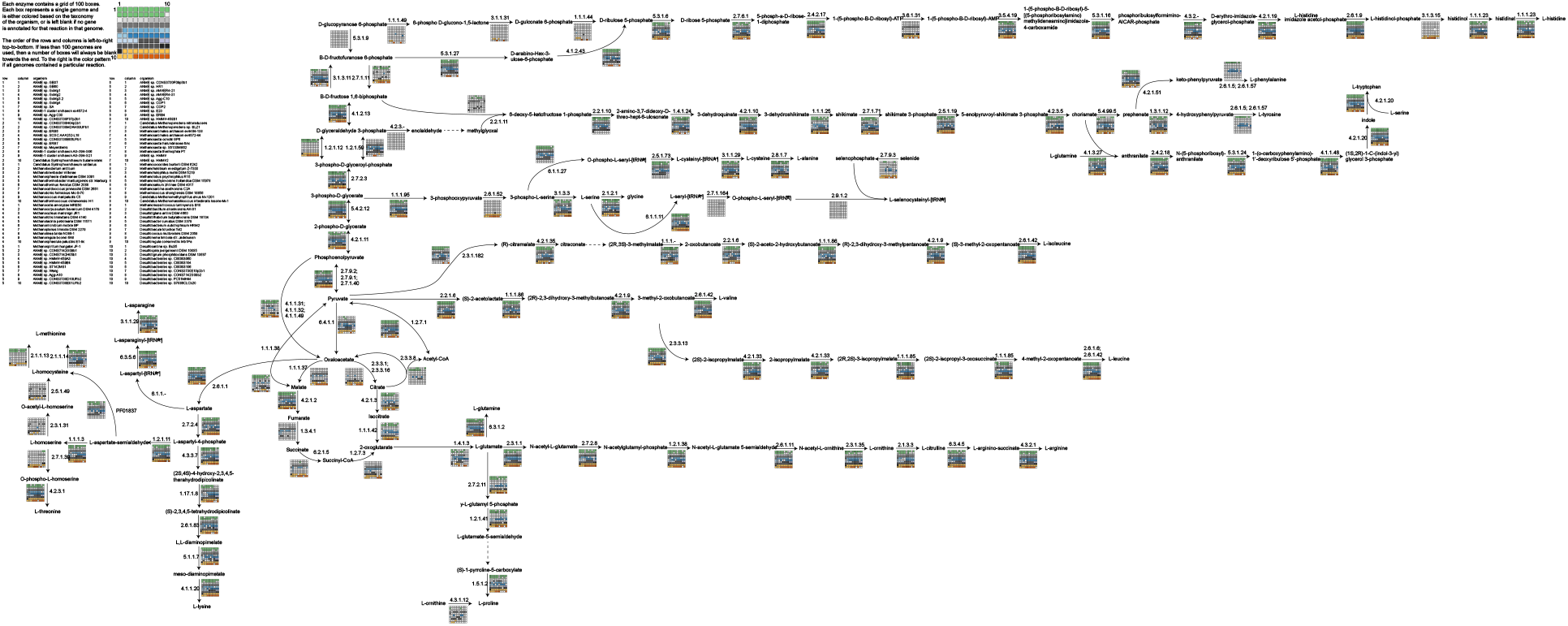
Amino acid pathways. Amino acid pathways present in ANME genomes, methanogens and sulfate reducing bacteria. Each enzymatic reaction is labelled with an enzyme commission (EC) number, or in one case where an EC number is not available, a Pfam accession number and a matrix of small colored boxes. Each colored box represents a single genome, if the box is colored it means that that genome contains an annotation for that enzyme, otherwise it is left blank. Genomes are colored based on their clade.

**S16 Fig.**
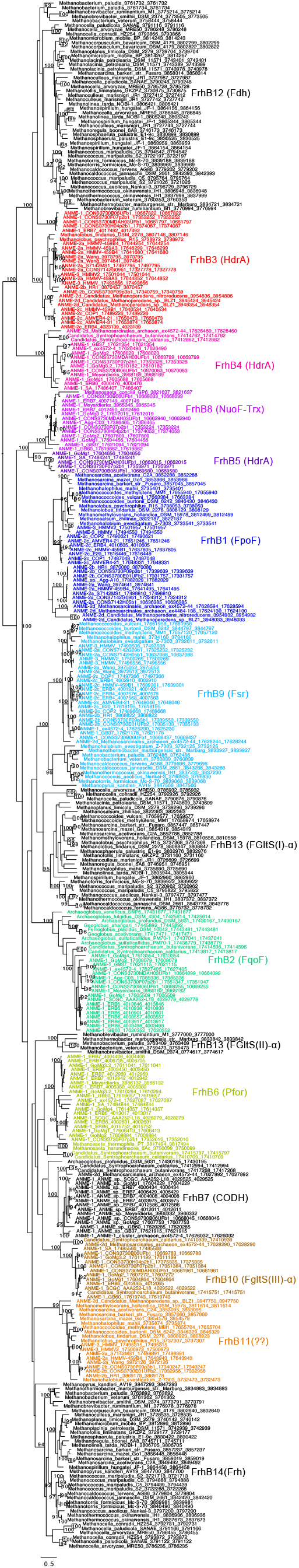
Expanded FdhB protein tree. FrhB paralogs are labelled as in Figure 16. Only branch support values >90% are shown for clarity. Tree scales represent substitutions per site. Tree construction parameters are found in the Materials and Methods section. Alignment and tree files can be found in **S1 Data**.

## Supporting Information

**S1 Data.** Alignments and tree files used in **Figs 1, 4**, **5**, **12**, **13**, **15-18**; and **S7**, **S8**, **S12** and **S14 Figs**.

**S2 Data.** Accession numbers for proteins represented in presence/absence figures presented in **Figs 3, 13**, **14**-**16**, and **18**; and **S6**, **S10** and **S11 Figs**.

**S3 Data.** Compilation of transcriptomic data from the ANME-2a Wang genome (11), the ANME-2d BLZ1 genome (19), and the ANME-2c E20, ANME-1 GB37 and GB60 genomes (9).

**S4 Data.** Spreadsheets listing domain annotations of all proteins in ANME genomes found to have dockerin or cohesin domains.

**S5 Data.** List of accession numbers for fosmids that were assigned to bins ERB4, ERB 6 and ERB 7.

**S1 File.** Formal etymology of proposed ANME genera.

**S1 Table.** Proteins comprising marker set 4 used for concatenated protein phylogenies.

**S2 Table.** 16S sequence accessions determined for single sorted aggregates.

## Notes

### Competing Interest Statement

The authors have declared no competing interest.

