## Supplementary material for "Comparative genomics reveals electron transfer and syntrophic mechanisms differentiating methanotrophic and methanogenic archaea": S1 File

Description of “*Ca.* Methanovorans” (gen. nov.) {ANME-3}

N.L. n. methanum [from French n. méth(yle) and chemical suffix -ane], methane; N.L. pref. methano-, pertaining to methane; L. part. adj. *vorans*, consuming; N.L. mas. adj. Methanovorans, consuming methane.

Description of “*Ca*. Methanogasteraceae” (fam. nov.) {ANME-2c}

N.L. n. methanum [from French n. méth(yle) and chemical suffix -ane], methane; N.L. pref. methano-, pertaining to methane; L. noun *gaster*, belly; N.L. fem. noun Methanogasteraceae, methane eater.

Description of “*Ca*. Methanogaster” (gen. nov.) {ANME-2c}

N.L. n. methanum [from French n. méth(yle) and chemical suffix -ane], methane; N.L. pref. methano-, pertaining to methane; L. noun *gaster*, belly; N.L. fem. noun Methanogaster, methane eater.

Description of “*Ca*. Methanocomedenaceae” (fam. nov.) {ANME-2ab}

N.L. n. methanum [from French n. méth(yle) and chemical suffix -ane], methane; N.L. pref. methano-, pertaining to methane; L. verb *comedo*, consume/devour; N. L. Methanocomedenaceae, consuming methane.

Description of “*Ca*. Methanocomedens” (gen. nov.) {ANME-2ab}

N.L. n. methanum [from French n. méth(yle) and chemical suffix -ane], methane; N.L. pref. methano-, pertaining to methane; L. verb *comedo*, consume/devour; N. L. Methanocomedens, consuming methane.

Description of “*Ca*. Methanomarinus” (gen. nov.) {ANME-2b}

N.L. n. methanum [from French n. méth(yle) and chemical suffix -ane], methane; N.L. pref. methano-, pertaining to methane; L. adj. *marinus*, pertaining to the sea; N. L. mac. adj. Methanomarinus, methane metabolizing organism from the sea.

Description of “*Ca*. Methanoalium” (gen. nov.) {ANME-1}

N.L. n. methanum [from French n. méth(yle) and chemical suffix -ane], methane; N.L. pref. methano-, pertaining to methane; L. adj. *alium*, other; N. L. mas. adj. Methanoalium, a different methane metabolizing organism.
